# iSCORE-PD: an isogenic stem cell collection to research Parkinson’s Disease

**DOI:** 10.1101/2024.02.12.579917

**Authors:** Oriol Busquets, Hanqin Li, Khaja Mohieddin Syed, Pilar Alvarez Jerez, Jesse Dunnack, Riana Lo Bu, Yogendra Verma, Gabriella R. Pangilinan, Annika Martin, Jannes Straub, YuXin Du, Vivien M. Simon, Steven Poser, Zipporiah Bush, Jessica Diaz, Atehsa Sahagun, Jianpu Gao, Samantha Hong, Dena G. Hernandez, Kristin S. Levine, Ezgi O. Booth, Marco Blanchette, Helen S. Bateup, Donald C. Rio, Cornelis Blauwendraat, Dirk Hockemeyer, Frank Soldner

## Abstract

Parkinson’s disease (PD) is a neurodegenerative disorder caused by complex genetic and environmental factors. Genome-edited human pluripotent stem cells (hPSCs) offer a unique experimental platform to advance our understanding of PD etiology by enabling the generation of disease-relevant cell types carrying patient mutations along with isogenic control cells. To facilitate this approach, we generated a collection of 65 human stem cell lines genetically engineered to harbor high risk or causal variants in genes associated with PD (*SNCA* A53T, *SNCA* A30P, *PRKN* Ex3del, *PINK1* Q129X, *DJ1/PARK7* Ex1-5del, *LRRK2* G2019S, *ATP13A2* FS, *FBXO7* R498X/FS, *DNAJC6* c.801 A>G/FS, *SYNJ1* R258Q/FS, *VPS13C* A444P/FS, *VPS13C* W395C/FS, *GBA1* IVS2+1/FS). All mutations were introduced into a fully characterized and sequenced female human embryonic stem cell (hESC) line (WIBR3; NIH approval number NIHhESC-10-0079) using different genome editing techniques. To ensure the genetic integrity of these cell lines, we implemented rigorous quality controls, including whole-genome sequencing of each line. Our analysis of the genetic variation in this cell line collection revealed that while genome editing, particularly using CRISPR/Cas9, can introduce rare off-target mutations, the predominant source of genetic variants arises from routine cell culture and are fixed in cell lines during clonal isolation. The observed genetic variation was minimal compared to that typically found in patient-derived iPSC experiments and predominantly affected non-coding regions of the genome. Importantly, our analysis outlines strategies for effectively managing genetic variation through stringent quality control measures and careful experimental design. This systematic approach ensures the high quality of our stem cell collection, highlights advantages of prime editing over conventional CRISPR/Cas9 methods and provides a roadmap for the generation of gene-edited hPSC collections at scale in an academic setting. Our iSCORE-PD collection represents an easily accessible and valuable platform to study PD, which can be used by investigators to understand the molecular pathophysiology of PD in a human cellular setting.

## Introduction

Parkinson’s disease (PD) is the second most common neurodegenerative disorder with a prevalence of more than 1% in the population over the age of 60^1^. PD is primarily characterized by a progressive loss of dopaminergic neurons in the midbrain and, in most cases, the presence of proteinaceous inclusions (Lewy bodies) in affected cells^2–5^. However, PD-associated pathology is highly variable and can affect a wide range of brain regions^6^. Furthermore, non-neuronal cell types, including astrocytes, oligodendrocytes and microglia play important roles in the pathogenesis of the disease^7^. The precise etiology leading to neuronal cell loss is largely unknown. The discovery of mutations in more than 20 genes linked to rare monogenic forms of PD has revealed a broad spectrum of molecular and cellular pathways that can contribute to PD pathology, including vesicle transport, lysosomal function, mitochondrial function, and endoplasmic reticulum (ER) quality control^4,8^. However, even individuals with PD who carry the same mutation can present with highly heterogeneous clinical and pathological features^5,9^, variable age of onset and highly diverse or, in some cases, complete absence of Lewy body pathology^6^. Recognizing this variability in penetrance and complex pathology, it is widely acknowledged that additional genetic and environmental modifiers can influence disease pathophysiology, even in monogenic forms of PD^4^. Therefore, distinguishing between common PD-associated phenotypic features from those that are specific to a particular mutation remains challenging.

Genome editing of human pluripotent stem cells (hPSCs), including both human embryonic stem cells (hESCs) and induced pluripotent stem cells (hiPSCs), is increasingly utilized to establish isogenic cellular models for human diseases such as PD^10–12^. This approach has provided valuable insights into the molecular mechanisms underlying monogenic forms of this disease^11–30^. It utilizes genome editing technologies such as CRISPR/Cas9 or prime editing systems to generate isogenic cell lines, either by genetically inserting or correcting disease-linked mutations in hPSCs^11,12^. While this approach provides the advantage to analyze the effects of a mutation within a presumably identical genetic background, the extent to which the genome of edited cell lines - beyond the intended genetic modifications - are truly isogenic remains unclear. Various sources of genetic variability can contribute to genetic alterations in hPSCs. These include off-target effects associated with genome editing, genetic drift during cell culture, and founder effects introduced during subcloning^31–35^. However, to date we lack a systematic quantification of the relative contribution of these events, raising the important question: to what extent can observed phenotypes be fully attributed to the intended genetic edits, rather than to additional acquired genetic alterations?

In addition, since most isogenic pairs are currently generated in individual hPSCs with distinct, patient-specific genetic backgrounds, cross-comparison of different mutations is confounded by the effect of genetic modifier loci inherent to each individual’s genome^11,36^. Therefore, a unified genetic interaction map of how different monogenic disease-related genes, their pathogenic mutations and their respective phenotypes interact to drive pathology is still missing. To overcome this challenge, ongoing initiatives, for a variety of diseases, exist with the goal of streamlining the development of isogenic, disease-relevant hPSC collections derived from a common, thoroughly characterized parental hPSC line^36^. Here, we report the generation of such a resource for PD as part of the Aligning Science Across Parkinson’s (ASAP) research network, which we have termed iSCORE-PD (Isogenic Stem Cell Collection to Research Parkinson’s Disease). We used state-of-the-art genome editing approaches to establish a total of 65 clonal cell lines carrying disease-causing or high-risk PD-associated mutations in 11 genes (*SNCA*, *PRKN*, *PINK1*, *DJ1/PARK7*, *LRRK2*, *ATP13A2*, *FBXO7*, *DNAJC6*, *SYNJ1*, *VPS13C*, and *GBA1*), along with isogenic control lines. All cell lines were derived from a well-characterized and fully sequenced female hESC (WIBR3; NIH approval number NIHhESC-10-0079)^37^ and underwent rigorous quality control. Importantly, we performed whole-genome sequencing (WGS) on all cell lines to address the fundamental question of isogenicity of genome edited hPSCs by assessing genetic variability within the iSCORE-PD collection. This collection of isogenic hPSCs is accessible to the community to enable the cross-comparison of disease-related phenotypes and accelerate progress in PD research.

## Results

### Characterization of the hESC line WIBR3

A major goal of this work is to complement ongoing initiatives to establish comprehensive collections of hPSC lines which carry mutations associated with PD and related neurodegenerative diseases^36^. The objective is to reduce genetic variability among cell lines to facilitate the identification of disease-relevant pathophysiological signatures. One example of such efforts is the recently described iPSC Neurodegenerative Disease Initiative (iNDI) from the NIH’s Center for Alzheimer’s and Related Dementias (CARD), which utilized the KOLF2.1J (RRID:CVCL_B5P3) hiPSC line^36^. This cell line, derived from a male donor, is currently recognized as a benchmark reference for neurodegenerative disease research, and facilitates the comparison of disease-associated phenotypes across different laboratories. Although hiPSCs have proven instrumental for disease modeling, concerns remain regarding the presence of genetic alterations in somatic donor cells before reprogramming, reprogramming-induced genetic alterations, incomplete epigenetic reprogramming, and aberrant genomic imprinting^11,38,39^. Given these considerations, and the necessity for incorporating cells from both sexes, we opted to use the female hESC line WIBR3 of European descent (RRID:CVCL_9767, NIH approval number NIHhESC-10-0079)^37^. This cell line has been previously demonstrated to maintain a stable karyotype over prolonged *in vitro* culture and has been widely used to model human diseases, including PD^10,30,37,40–45^. For our study, we acquired early passage WIBR3 cells (P14) and initially generated 3 independent single cell-derived subclones (WIBR3-S1, WIBR3-S2, WIBR3-S3). Both the parental line and its subclones showed regular growth and morphology when cultured on mouse fibroblast feeders (MEFs) and under feeder free conditions in mTeSR™ Plus media (Figure 1A). We validated the pluripotency of all cell lines through the detection of pluripotency markers using immunocytochemistry and qRT-PCR (Figure 1B,C; Supplemental Figure 1A). Furthermore, we analyzed the genomic integrity of all cell lines using standard array comparative genomic hybridization (aCGH) and a modified high density Illumina Infinium Global Diversity Array (GDA) Neuro booster Array (NBA). This analysis confirmed a normal karyotype and the absence of larger structural alterations (> ∼500 kb) in both the parental WIBR3 line and its derived subclones (complete karyotype data is available at https://www.amp-pd.org/).

**Figure 1.**
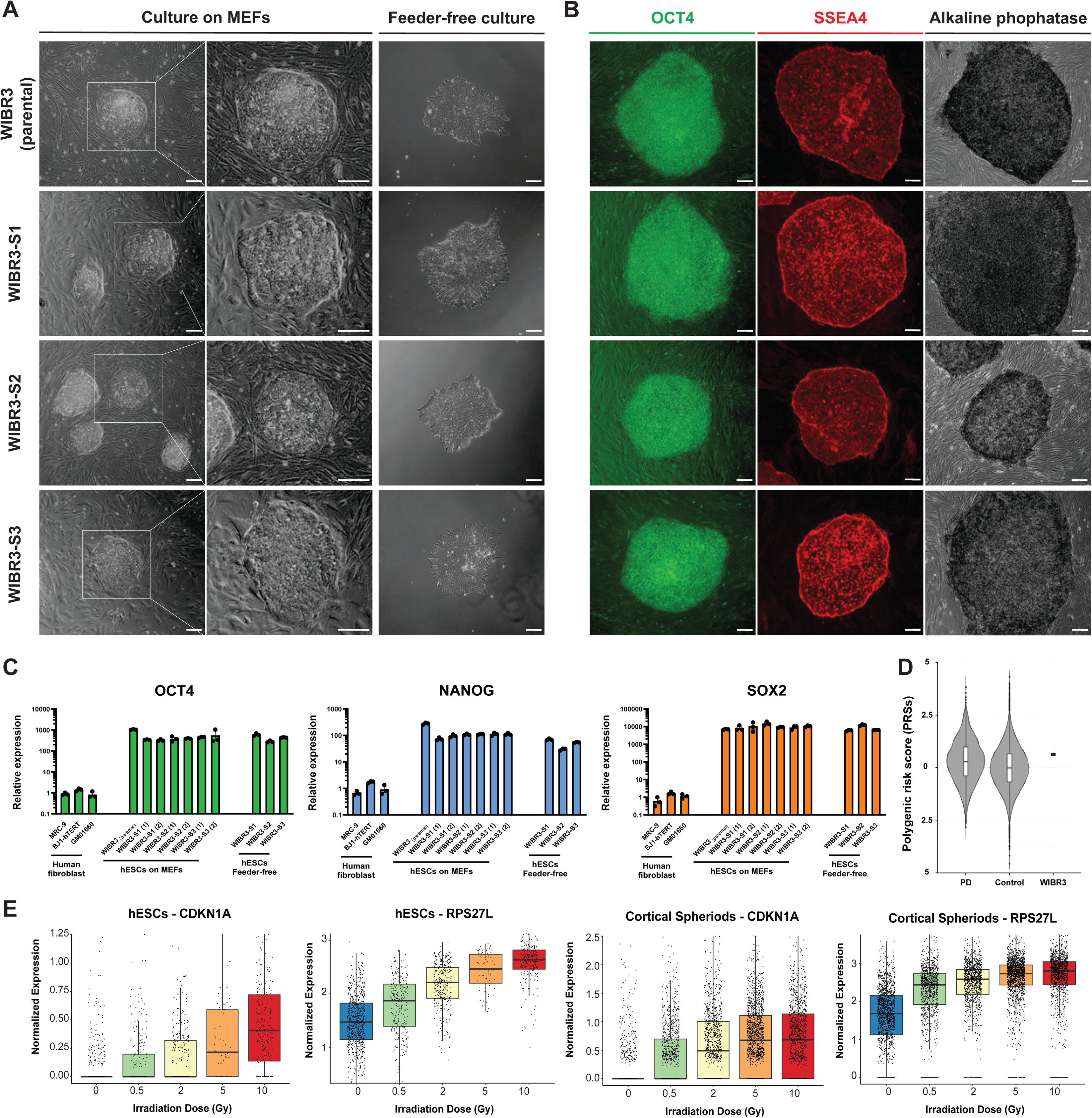
WIBR3 hESC cell line characterization. (A) Phase contrast images of parental WIBR3 hESCs and subclones WIBR3-S1, WIBR3-S2 and WIBR3-S3 cultured on MEFs and in feeder-free conditions. Scale bar 100 µm. (B) Immunocytochemistry for pluripotency markers OCT4 (green) and SSEA4 (red) and staining for alkaline phosphatase (black) of WIBR3 (parental) hESCs and subclones WIBR3-S1, WIBR3-S2 and WIBR3-S3 cultured on MEFs. Scale bar 100 µm. (C) qRT-PCR analysis for the relative expression of pluripotency markers OCT4, NANOG and SOX2 in human primary fibroblasts (MRC-9, BJ1-hTERT and GM01660), WIBR3 (parental) hESCs and subclones WIBR3-S1, WIBR3-S2 and WIBR3-S3 hESCs cultured on MEFs and in feeder-free conditions. Relative expression levels were normalized to expression of these genes in primary fibroblasts. (1) and (2) indicate independent samples. (N=3; Mean +/-SEM). (D) Polygenic risk scores (PRSs) for PD comparing WIBR3 hESCs to population-centered Z score distribution for PD PRSs in individuals with PD and the normal population from the UK Biobank. (E) Assessment of p53 pathway activity following irradiation (0.5, 2, 5 and 10 Gy) of WIBR3 (parental) hESCs (1464 cells) and WIBR3-derived cortical spheroids (5920 cells) by scRNA-seq analysis for the expression of DNA damage response genes CDKN1A and RPS27L (box plot showing interquartile intervals with a line at the median).

To determine the presence of insertions and deletions (indels) at higher resolution and to identify potential pathogenic single nucleotide variants (SNVs), we performed long-read whole genome sequencing (WGS) combining Pacific Biosciences (PacBio, average coverage 28.39X, median read length 18 kb) and Oxford Nanopore Technologies (Nanopore, average coverage 43X, median read length 84 kb). The initial analysis for structural variants applied the Truvari algorithm^46^ (https://github.com/ACEnglish/truvari) to integrate the PacBio and Nanopore datasets and identified a total of 20,561 high confidence structural variants in the WIBR3 parental line compared to the reference human genome [GRCh38/hg38] (Supplemental Table 1). Among these, 109 were localized to coding exons, impacting 102 genes (Supplemental Table 1). The number and distribution of these structural variants is comparable to those observed in the general human population^47,48^. Considering our goal to model PD and related neurodegenerative diseases, we determined that none of these structural variants affect genes with known pathogenic mutations in PD, Alzheimer’s disease (AD), and AD-related dementias (ADRD), or risk genes identified in GWAS associated with these diseases^4,49–51^ (Supplemental Table 1). Additionally, we annotated the integrated structural variant calls using SvAnna^52^(https://github.com/TheJacksonLaboratory/SvAnna) to determine if any variant was of high priority for the phenotype terms HP:0002180 (neurodegeneration) and HP:0000707 (abnormality of the nervous system). None of the structural variants analyzed received high priority scores for either term.

Next, we identified the number and distribution of coding, missense, frameshift and predicted loss of function (LOF) SNVs in the parental WIBR3 cell line compared to the reference human genome [GRCh38/hg38] (Supplemental Figure 1B). This analysis identified 6613 missense SNVs in 3933 coding genes and 120 potential loss of function mutations (including 15 startloss, 54 stopgain, 13 stoploss, 18 frameshift deletion, and 20 frameshift insertion). A full list of variants can be found in Supplemental Table 2, and the full genome is available for broad data sharing (https://www.amp-pd.org/ via GP2 data sharing agreement). Overall, the number and distribution of these variants is comparable to that observed in other sequenced hPSC lines^36^ and within the human population found in the gnomAD database (https://gnomad.broadinstitute.org/)^53,54^.

To determine the presence of potentially pathogenic variants in the parental WIBR3 cell line, we annotated all SNVs using ClinVar^46^ (https://www.ncbi.nlm.nih.gov/clinvar/). Collectively, these analyses revealed 48 variants either listed as pathogenic or having conflicting interpretations of pathogenicity. However, none of these SNVs are linked to a neurological phenotype of interest or are convincingly pathogenic (Supplemental Table 2). As we aim to provide our cell collection to study PD and related neurodegenerative diseases, we analyzed WGS data to calculate the polygenic risk score (PRS) based on the cumulative number of GWAS risk variants associated with PD (Figure 1D)^49^. The analysis indicated that the PRS of the parental WIBR3 line falls within the range observed in the normal population. Subsequently, we focused on identifying high-risk variants in known neurodegenerative disease-associated genes. This analysis revealed that WIBR3 is heterozygous for the *APOE* ε4 allele (worldwide allele frequency of e4 ∼14%), which is a risk factor for Alzheimer’s disease^55^, heterozygous for *rs3173615* (TMEM106B p.T185S) which has been reported to be a modifier of frontotemporal dementia^56^, and homozygous for the *MAPT* H1 allele which is a gene of interest in several neurodegenerative diseases^57,58^.

It is widely recognized that hPSCs can accumulate genetic alterations over time which provide a growth advantage in cell culture. Notably, mutations in the p53 tumor suppressor pathway have been frequently observed in various hPSC lines^59–61^. To evaluate the function of the p53 pathway in the parental WIBR3 cell line, we analyzed the p53-dependent DNA damage response following irradiation (Figure 1E). This analysis confirmed a robust p53-mediated response, as indicated by the dose-dependent expression of the DNA damage response genes CDKN1A and RPS27L in both, undifferentiated hPSCs and *in vitro*-derived cortical spheroids (Figure 1E).

### WIBR3 cells differentiate into PD-relevant cell types

Given that PD is characterized by the chronic progressive loss of dopaminergic (DA) neurons in the substantia nigra, the effective generation of these cell types is crucial for *in vitro* modeling of PD. To address this, we implemented a previously established protocol^62,63^ to differentiate the three independently derived WIBR3 subclones (WIBR3-S1, WIBR3-S2, WIBR3-S3) into midbrain-specified dopaminergic neurons. Briefly, WIBR3 hESCs underwent neural induction via dual SMAD inhibition, combined with sonic hedgehog (SHH) agonist exposure and biphasic WNT activation using the GSK-3 inhibitor CHIR99021, resulting in robust midbrain-specific patterning within the first 11 days. From day 12 onwards, committed midbrain neural progenitors were differentiated into dopamine neurons until day 35 using a cocktail to promote terminal differentiation (BDNF, GDNF, TGFß3, DAPT, cAMP and ascorbic acid) (Figure 2A). For each subclone, we determined the efficacy of neural induction into neural precursor cells and DA neurons by analyzing the expression of midbrain-specific markers by immunocytochemistry (TH and FOXA2) and, qRT-PCR (FOXA2, LMX1A, NR4A2, KCNJ6, TH, PITX3, EN2, AADC and SYN1) (Figure 2B, Supplemental Figure 2). At early time points (day 11 and day 25), the *in vitro* differentiated cultures expressed canonical midbrain floor plate genes at levels comparable to those in concurrently differentiated KOLF2.1J hiPSCs (Supplemental Figure 2B). At day 35 of differentiation, over 80% of cells expressed FOXA2, a marker for early midbrain floorplate neuronal precursors, and approximately 20% expressed the dopaminergic neuron marker tyrosine hydroxylase (TH), indicating the generation of midbrain-specified dopaminergic neurons (Figure 2B; Supplemental Figure 2A).

**Figure 2.**
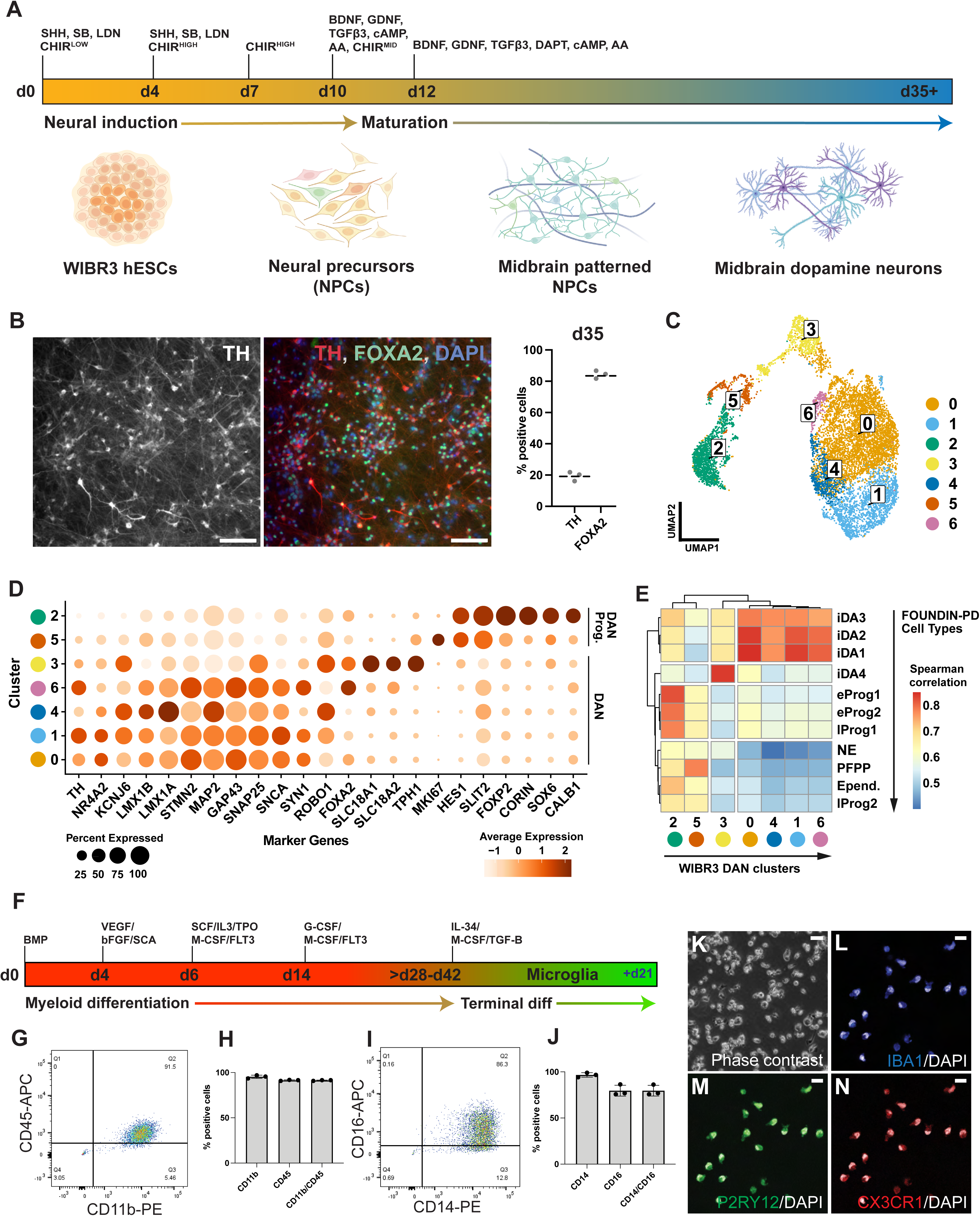
WIBR3 differentiation potential into dopaminergic neurons and microglia subtypes in 2D culture. (A) Schematic depicting the protocol for in vitro differentiation of dopaminergic neurons from WIBR3 hESCs. (B) Immunocytochemistry and quantification of TH and FOXA2 expressing cells in WIBR3 (parental) hESC-derived dopaminergic neurons at day 35. Scale bar 100 µm. (N=3). Check supplemental figure 2A for immunocytochemistry images from WIBR3-S1,2,3. (C) Uniform manifold approximation and projection (UMAP) plot of scRNA-Seq analysis at day 35-37 of dopaminergic neuron differentiation from WIBR3-S1, WIBR3-S2 and WIBR3-S3 hESCs showing 10,097 cells separated into 7 coarse clusters. (D) Dot plot showing expression of key progenitor and mature dopamine neuron marker genes across different cluster identities indicates that clusters identified in (C) represent dopamine neuronal progenitors and dopaminergic neurons at different developmental stages. Dot size indicates the proportion of cells in a cluster expressing a given gene, while color intensity indicates its average expression. Although cluster 2 showed *CALB1* expression, we labelled these cells a progenitor population due to expression of *HES1, SLIT2, CORIN* and absence of mature dopamine neuron markers. (E) Heatmap depicting Spearman correlation coefficients between pseudo-bulk expression profiles of 7 WIBR3 clusters identified in (C) compared to pseudo-bulk expression profiles of 11 FOUNDIN-PD cell types. (F) Schematic depicting in vitro microglia differentiation protocol from WIBR3 hESCs. (G-J) Representative flow cytometry (FACS) analysis (G,I) and quantification (H,J) of CD11b/CD45 and CD14/CD16 expression in hESC-derived iMPs from subclones WIBR3-S1, WIBR3-S2 and WIBR3-S3. (K-N) Representative phase contrast (K, Scale bar 50 µm) and immunostaining (L-N, Scale bar 10 µm) images of in vitro differentiated microglia derived from subclone WIBR3-S1 for microglia-specific markers IBA1 (blue), P2RY12 (green)) and CX3CR1 (red) (terminal differentiation day 14).

To further characterize the DA neuron cultures, we performed single-cell RNA sequencing (scRNA-seq) at day 35-37 post-differentiation from hESCs. Cells were profiled across the three independently assayed subclones, yielding an aggregate dataset of 10,097 cells. Using Uniform Manifold Approximation and Projection (UMAP) for dimensionality reduction, seven distinct clusters were identified in the integrated dataset, each composed of cells from all three subclones (Figure 2C, Supplemental Figure 3A). Among these, clusters 0, 1, 3, 4 and 6 showed strong expression of canonical dopaminergic neuron markers (*KCNJ6, TH, NR4A2*), whereas clusters 2 and 5 displayed strong expression of dopaminergic neuronal progenitor markers (*SLIT2, FOXP2*, *CALB1, SOX6 and CORIN*) (Figure 2D, Supplemental Figure 3B-D)^64,65^. To further compare the differentiation propensity of WIBR3 cells against other cell lines with different genetic backgrounds, we compared our dataset with the recently published Foundational Data Initiative for Parkinson’s Disease (FOUNDIN-PD) data (Figure 2E)^65^. The FOUNDIN-PD dataset includes single-cell RNA-seq data from midbrain DA neuron cultures at day 65, derived from 80 distinct hiPSC lines using a comparable *in vitro* differentiation protocol^65^. This comparison revealed that the clusters representing DA neuron populations (clusters 0, 1, 3, 4, and 6) in our dataset showed the highest Spearman correlation scores with iDA1, iDA2, iDA3 and iDA4 neuron clusters identified in the FOUNDIN-PD data, and clusters 2 and 5 from our dataset correlate more strongly with progenitor populations (Supplemental Figure 4). Together, this analysis indicates a high similarity between the expression profiles of our WIBR3-derived cell types and those in the FOUNDIN-PD dataset, which is currently the most comprehensive and standard data set for *in vitro*-derived midbrain-specific DA neurons. This is relevant, as it should allow the integration of data generated from the iSCORE-PD collection with the FOUNDIN-PD datasets which include 80 hiPSC lines from patients with sporadic and familial PD, as well as age-matched healthy individuals.

Recent data highlighted that the impact of PD-associated mutations extends beyond neurons, affecting cell types such as microglia, which play a critical role in the pathogenesis of PD^66,67^. Chronic microglial activation is suggested to be a key pathophysiological feature of many neurodegenerative disorders, including PD^68^. To this point, we followed a previously described protocol^69^ to differentiate the subclones of WIBR3 (WIBR3-S1, WIBR3-S2, WIBR3-S3) into microglia-like cells (iMGs). In this protocol, hPSCs are initially induced to myeloid intermediates and subsequently differentiated into microglia-like cells through the addition of cytokines, normally secreted from neurons and astrocytes including IL-34, M-CSF, and TGF-β1 (Figure 2F). All WIBR3 subclones robustly generated microglial precursors (iMPs), evidenced by the presence of a high percentage of cells expressing markers tied to the microglial lineage (91.3% CD11b/CD45 and 79.6% CD14/CD16) (Figure 2G-J). Moreover, terminal differentiation yielded cells expressing key markers for mature microglia such as IBA1, CX3CR1, and P2RY12, as confirmed through immunostaining (Figure 2K-N, Supplemental Figure 5). Collectively, these findings underscore the suitability of WIBR3 hESCs as a model to study the contribution of different cell types to PD pathology.

### Genetic engineering of PD-associated mutations requires multiple editing modalities including prime editing

The establishment of a large-scale collection of cell lines carrying disease-associated mutations requires precise and robust gene editing approaches to insert the desired genetic alterations in hPSCs. We previously demonstrated that WIBR3 cells can be genetically modified with high efficiency using either CRISPR/Cas9, TALEN or prime editing-based genome engineering approaches^40,70,71^. CRISPR/Cas9-based editing is effective for introducing targeted genomic deletions and biallelic alterations, while prime editing is highly efficient in introducing heterozygous modifications, which is necessary for modeling dominantly inherited disease-associated alleles^70^. To create cell lines carrying PD-associated genetic alterations in WIBR3 hESCs, we employed two different editing pipelines. Pipeline A (Figure 3A) utilizes FACS-enrichment post-nucleofection to purify effectively transfected cells, followed by clonal expansion and genotyping to allow the isolation of correctly targeted lines. The estimated time for the editing pipeline A is 30-35 days. Pipeline B (Figure 3B) uses nucleofection, limited dilution, and next generation sequencing (NGS)-based genotyping to identify desired edits in a 96-well plate format. This integrated workflow allows for the efficient isolation and purification of correctly edited clonal cell lines, even at low frequency, within a shorter time frame compared to previous approaches (21-35 days).

**Figure 3.**
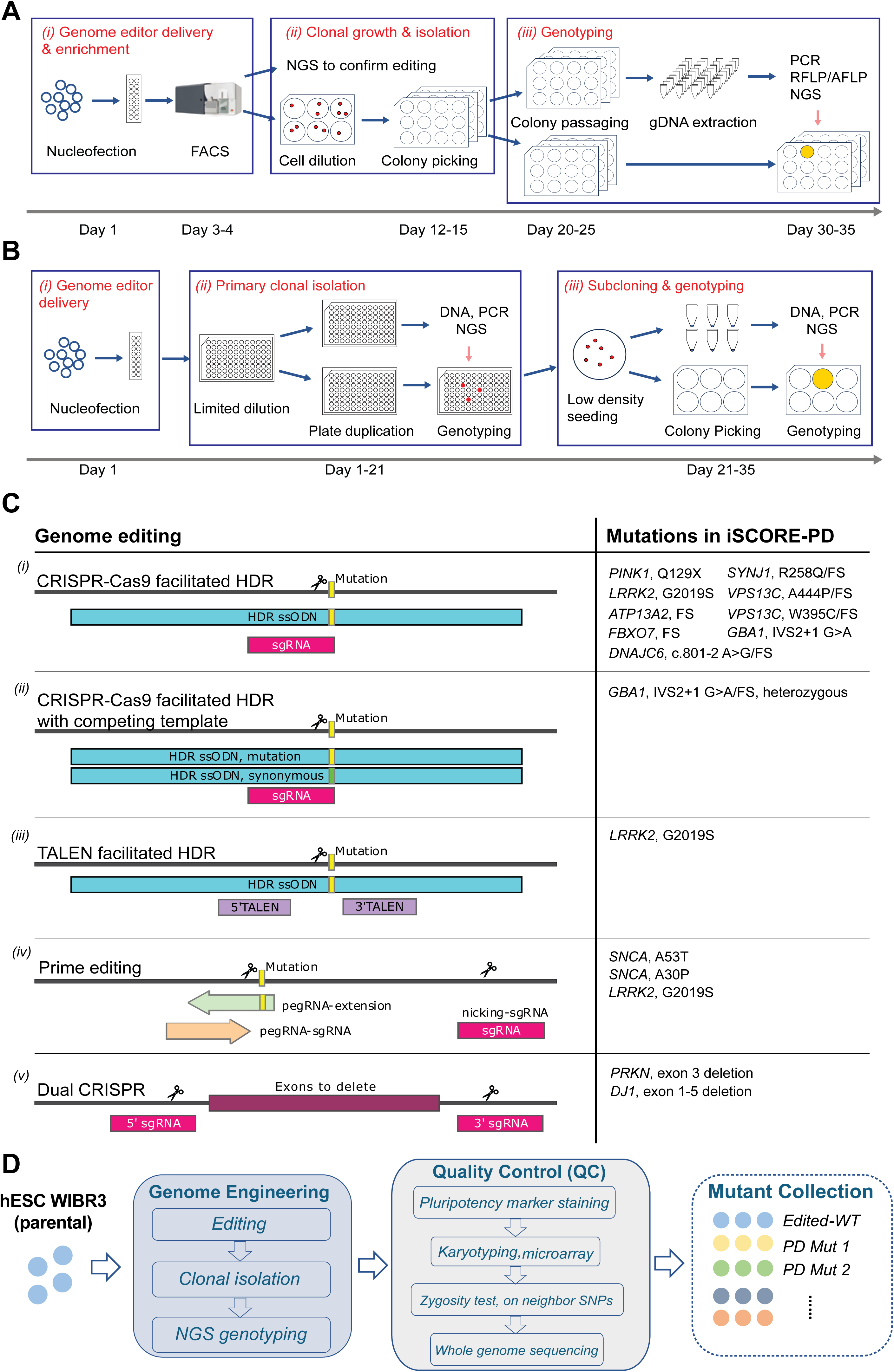
Gene editing workflow to generate iSCORE-PD collection. (A) Schematic illustrating genome editing pipeline A. This approach involves *(i)* FACS-based enrichment of nucleofected cells containing the gene editing reagents including a fluorescent reporter, *(ii)* the isolation of clonally expanded cell lines and *(iii)* the NGS-based genotyping to identify correctly edited cell lines. (B) Schematic illustrating genome editing pipeline B. This approach utilizes a high-throughput cell isolation system. This approach includes *(i)* nucleofection of the gene editing reagents, *(ii)* the plating of cells in a limited dilution (∼10 cells/well) to isolate wells containing correctly targeted cells by NGS and *(iii)* subcloning, expansion and NGS-based genotyping to isolate correctly targeted clonal cell line. (C) Table summarizing the gene editing strategies used to generate the iSCORE-PD collection. These include: *(i)* CRISPR/Cas9 facilitated homology directed repair (HDR) using ssODNs containing the desired genetic modification as repair template for CRISPR/Cas9 induced double strand break. *(ii)* The use of competing HDR templates (ssODNs) containing synonymous mutations in the gRNA-target site to favor the generation of heterozygous over homozygous mutations. *(iii)* TALEN-facilitated HDR using ssODNs containing the desired genetic modification as repair template for CRISPR/Cas9 induced double strand break. *(iv)* Prime editing approach to insert the PD-associated point mutations into hESCs. *(v)* Dual CRISPR approach using 3’ and 5’ sgRNAs flanking the desired deletion to recreate large genomic structural alterations identified in PD patients. (D) Overview depicting genome engineering and quality control steps in the generation of the iSCORE-PD collection.

Given the identification of several disease-causing mutations within most PD-linked genes, the selection and prioritization of specific alleles for gene editing within each gene were based on confidence in pathogenicity for each mutation, allele prevalence, and feasibility of the editing strategy. As described in detail in the supplementary information for each gene (Supplementary Note 1, Supplemental Figures 6-17), we employed three general editing strategies to closely recreate the genomic alterations identified to be causal or high-risk factors for PD. These editing strategies include *(1)* the precise insertion of point mutations using CRISPR/Cas9, TALEN or prime editing approaches to recreate specific PD-associated missense mutations (heterozygous and/or homozygous), *(2)* the insertion of small indels to create frameshift (FS) or premature stop mutations using CRISPR/Cas9, and *(3)* dual guided RNA (gRNA)-mediated CRISPR/Cas9 editing to create genomic deletions identified in PD patients (Figure 3C).

### iSCORE-PD: a cell line collection of isogenic hPSC lines carrying PD-associated mutations

To establish iSCORE-PD, a collection of isogenic hPSC lines carrying monogenic or high-risk variants linked to PD, we initially prioritized engineering mutations in high-confidence PD genes^4^. The specific modifications for each gene were selected based on information in the MDSgene database^72^ (https://www.mdsgene.org) and the currently available literature as outlined in detail for each gene in the supplementary information (Supplementary Note 1). Overall, the current iteration of the iSCORE-PD collection includes 65 clonal cell lines carrying high-risk or causal variants in 11 genes linked to PD (*SNCA* A53T, *SNCA* A30P, *PRKN* Ex3del, *PINK1* Q129X, *DJ1/PARK7* Ex1-5del, *LRRK2* G2019S, *ATP13A2* FS, *FBXO7* R498X/FS, *DNAJC6* c.801 A>G/FS, *SYNJ1* R258Q/FS, *VPS13C* A444P/FS, *VPS13C* W395C/FS, *GBA1* IVS2+1/FS) and isogenic control lines. All cell lines (Table 1, Supplemental Table 3) passed all the below-described quality control steps (Figure 3D) and will be available to the scientific community through the WiCell Research Institute (https://www.wicell.org/). Our intention is to continue to expand this collection in the future. A detailed discussion for each cell line can be found in the supplementary information (Supplementary Note 1, Supplemental Table 3, Supplemental Figures 6-17).

**Table 1.** iSCORE-PD cell line collection summary (targeted genes, mutations generated, genetic inheritance, editing approach, number of cell lines and their genotype). ^1^ mixed inheritance.

|  | MUTATION | INHERITANCE | EDITING | CELLS IN COLLECTION |  |
| --- | --- | --- | --- | --- | --- |
|  |  |  |  | MONOALLELIC -<br>HETEROZYGOUS | BIALLELIC -<br>HOMOZYGOUS |
| **(PARK1) SNCA** | A53T | Autosomal dominant | Prime editing | 3 | - |
|  | A30P | Autosomal dominant | Prime editing | 3 | 1 |
| **(PARK2) PRKN** | EX3DEL | Autosomal recessive | CRISPR/Cas9 (dual guide) | - | 3 |
| **(PARK6) PINK1** | Q129X | Autosomal recessive | CRISPR/Cas9 (HDR) | - | 3 |
| **(PARK7) DJ1** | EX1-5DEL | Autosomal recessive | CRISPR/Cas9 (dual guide) | 5 | 7 |
| **(PARK8) LRRK2** | G2019S | Autosomal dominant | Prime editing, TALEN and<br>CRISPR/Cas9 (HDR) | 4 | 1 |
| **(PARK9) ATP13A2** | FS | Autosomal recessive | CRISPR/Cas9 (HDR) | - | 4 |
| **(PARK15) FBXO7** | R498X/FS | Autosomal recessive | CRISPR/Cas9 (HDR) | - | 2 |
| **(PARK19) DNAJC6** | c.801-2<br>A>G/FS | Autosomal recessive | CRISPR/Cas9 (HDR) | - | 2 |
| **(PARK20) SYNJ1** | R258Q/FS | Autosomal recessive | CRISPR/Cas9 (HDR) | - | 2 |
| **(PARK23) VPS13C** | W395C/FS | Autosomal recessive | CRISPR/Cas9 (HDR) | - | 4 |
|  | A444P/FS | Autosomal recessive | CRISPR/Cas9 (HDR) | 1 | 2 |
| **GBA1** | IVS2+1/FS | Autosomal dominant<sup>1</sup> | CRISPR/Cas9 (HDR) | 4 | 2 |
| **Wild type** | - | - |  | Parental + 3 independent clones |  |
| **Edited wild type (EWT)** | - | - | Prime editing,<br>CRISPR/Cas9 (HDR) | Prime editing (6) + CRISPR/Cas9 (2) |  |

An important consideration in hPSC-based disease modeling is the selection of appropriate control lines. To address this, we provide a set of subclones derived from the parental WIBR3 cell line (WIBR3-S1, S2, S3) (Figure 1, Supplemental figure 1). Additionally, we included WIBR3 cell lines that were isolated as part of the standard genome editing experiment but did not exhibit any genetic modifications at the targeted locus, referred to as “edited wildtype” (EWT) cells. We consider these cells as preferred experimental controls, as they most effectively should account for any non-specific changes induced by the gene-editing process. The EWT cell lines include: EWT1-3 (prime editing controls - Pipeline B), EWT4-5 (CRISPR/Cas9 controls - Pipeline B) and EWT6-8 (prime editing controls - Pipeline A) (Table1, Supplemental Note 1, Supplemental Table 3, Supplemental Figure 6).

### Generation of hPSC collections requires rigorous quality control

A significant challenge for any genome editing approach is the risk of introducing unintended on- and off-target genetic modifications in the edited cell lines. Additionally, it is well-established that clonal expansion and *in vitro* culture of hPSCs can lead to the acquisition of genetic alterations that can provide growth advantages^34,59–61,73–75^. Consequently, there is a consensus in the field that gene-edited hPSC-derived disease models should undergo a rigorous quality control process to validate the pluripotency of edited cell lines, and to ensure the absence of major gene editing- or culture-induced genetic alterations. As part of this collection, all genome-edited cell lines underwent a comprehensive quality control process, as outlined in Figure 3D. Following genome editing and subsequent clonal expansion, individual correctly targeted clones were initially identified using either Sanger sequencing or NGS. All correctly targeted cell lines were subsequently expanded, cryopreserved at a low passage, and assayed by immunocytochemistry for the expression of pluripotency markers OCT4, SSEA4, and alkaline phosphatase. To confirm a normal karyotype and assess overall genomic integrity, genome-edited clonal hESC lines underwent standard aCGH karyotyping and were analyzed using a modified high density Illumina Infinium Global Diversity Array (GDA) Neuro booster Array (NBA). This analysis aimed to identify cell lines with large genome editing-induced structural alterations or complete chromosomal loss compared to the genome of the parental WIBR3 cell line (complete high-density array genotyping data is available at https://www.amp-pd.org/).

A frequently overlooked challenge associated with genotyping approaches based on PCR amplification of the target locus is the common failure to detect loss of heterozygosity (LOH)^76^. LOH result from large deletions or the loss of entire chromosome fragments distal to the site targeted during the genome editing process. To rule out LOH in cell lines that appeared to be homozygously edited based on the detection of a single allele by NGS, we introduced an additional quality control step. We used either a southern blot or SNV-PCR based analysis to validate the presence of two alleles at the targeting site, as described in detail for each gene in the supplementary information (Supplementary Note 1, Supplemental Table 3, Supplemental Figures 6-17). Using this analysis, we identified LOH in 2 out of 18 (11.11%) tested clonal cell lines initially classified as correctly edited with two identical alleles at the target site. These data underscore that LOH is a significant complication arising from genome editing and emphasize the importance of incorporating LOH testing as an important component of the quality control process for genome-edited hPSC lines. Any cell lines showing alterations in any of the quality control assessments described above were removed from the collection.

A summarized list of the genes, mutations, and number of cell lines in the iSCORE-PD collection is provided in Table 1. For detailed information regarding the gene editing process and quality control of all analyzed clonal cell lines in the generation of the iSCORE-PD collection, see Supplemental Note 1, Supplemental Table 3, 4 and Supplemental Figures 6-17. Overall, 19.75% (16 out of 81) of isolated clonal cell lines with correct NGS-confirmed genotype were excluded from our collection. As summarized in Supplemental Table 4, reasons for exclusion include chromosomal and structural alterations (16.05% - 13 out of 81 lines analyzed), lack of pluripotency marker expression (1.47% - 1 out of 68 lines analyzed) and LOH (11.11% - 2 out of 18 lines analyzed). It is important to note that the frequency of chromosomal and large structural abnormalities was higher in clones generated by double strand break-based genome editing (CRISPR and TALEN, 20.34% - 12 out of 59 lines analyzed) compared to prime editing (5.56% - 1 out of 18 lines analyzed). Similarly, LOH at the targeted locus was only observed in CRISPR/Cas9 edited cell lines and absent in prime edited cell lines.

### Genetic variability between cell lines in the collection is largely driven by preexisting spontaneous mutations

Various sources of genetic alterations — beyond the intended genome edits — can contribute to genetic variability in hPSCs that potentially can affect the phenotypic analysis of hPSC derived cells. As outlined in Figure 4A, genetic variation can arise from either spontaneous mutations that result from imprecise DNA replication or DNA repair after damage^34,77^, as well as from non-random off-target effects associated with genome editing^32,33,78^. As most of these mutations — similar to somatic mutations found ubiquitously across normal tissues — do not strongly impact cellular fitness^77^, cell lines comprise a complex mosaicism of subpopulations with fluctuating allele frequencies that are subject to genetic drift during cell culture, and founder effects introduced during subcloning. This inherent genetic variability raises a fundamental question for hPSC-derived disease models: how confidently can we attribute observed phenotypes to the intended genetic edits, rather than to additional acquired genetic alterations?

**Figure 4.**
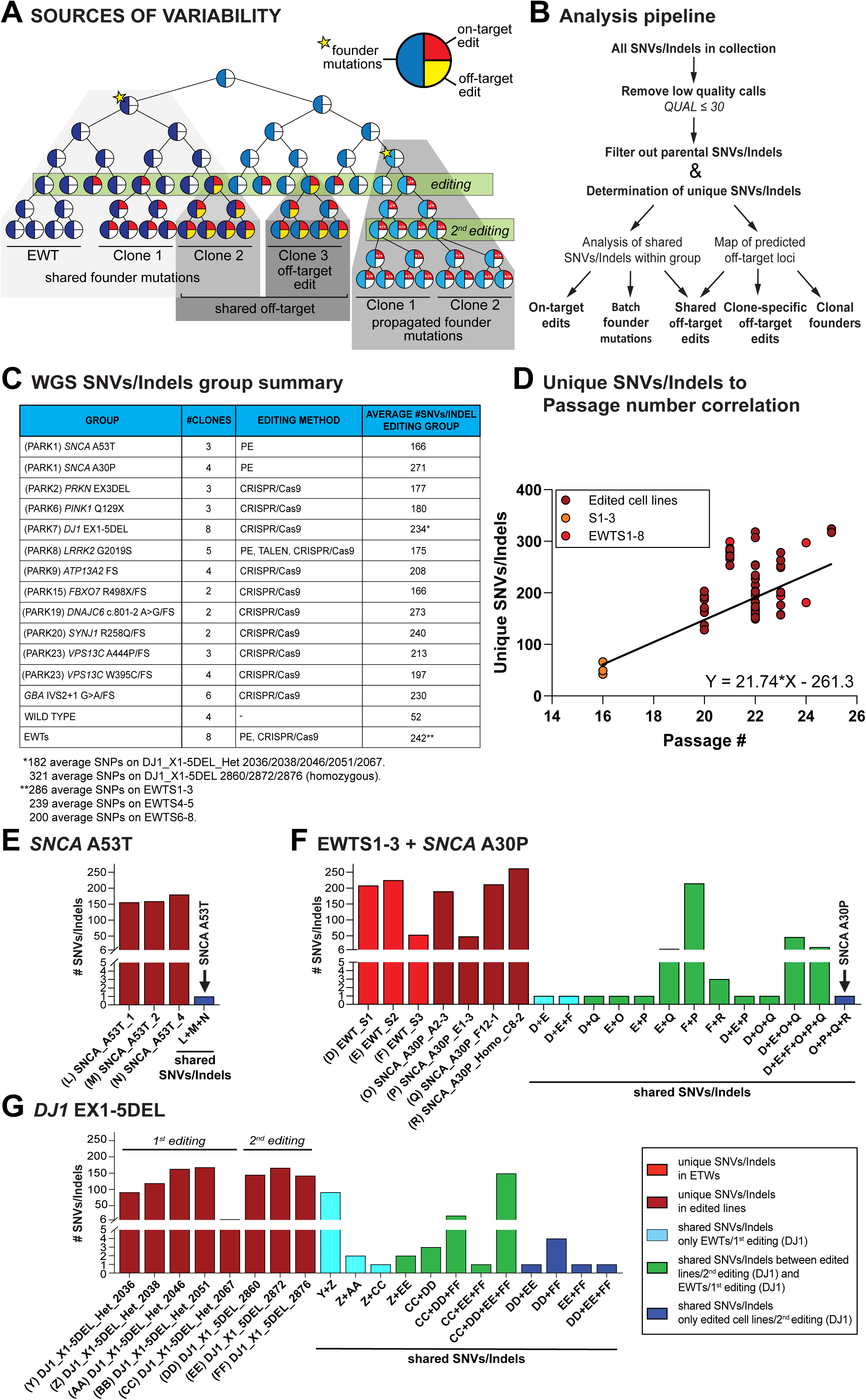
Genetic variation in the iSCORE-PD collection. (A) Schematic representation of the potential source of genetic variability found in the iSCORE-PD collection, including founder mutations from normal in vitro culture, propagated founder mutations during clonal expansion, and genome editing-induced variations (on- and off-target edits). For clarity of illustration, different shades of blue represent distinct sets of founder mutations. (B) Schematic of WGS analysis pipeline to identify unique and shared SNVs/indels for each editing group. Variants (SNVs/Indels) were initially mapped relative to the reference genome (GRCh38) and low-quality calls (QUAL≤30) were removed. To identify SNVs/Indels specific to an editing group, variants present in any other cell line within the collection were removed as a proxy for parental SNVs/indels in WIBR3. These editing group-specific SNVs/indels were then used to classify unique and shared variants within each edited group (on-target edits, founder mutations, and shared off-targets) and to assess their contribution to predicted off-target loci (clone-specific off-targets, clone founders, and shared off-targets). C) Table of all editing groups in iSCORE-PD collection and SNVs/Indels related information: number of clones, editing method used to generate them and average SNV/Indel count. (D) Correlation between the number of unique SNVs/Indels and passage number of last clonal event for each cell line. Regression line: y=22.6x-273.6; R2: 0.3751; 95% CI: 14.93 to 30.26. Orange indicates parental subclone lines (WIBR3_S1/S2/S3), light red indicates WIBR3_EWTS1-8 and dark red all other edited cell lines. (E) Graph showing number of unique and shared SNVs/indels in SNCA A53T group. No shared SNVs/indels beside edited SNCA A53T mutation were detected. (F) Graph showing number of unique and shared SNVs/indels in SNCA A30P group and EWT1-3 isolated from the same editing experiment. Shared SNVs/indels interactions between both edited and unedited cell lines show evidence of a founder effects. (G) Graph showing number of unique and shared SNVs in DJ1_X1-5DEL group. Data shows a significant clonal founder effect between DJ1_X1-5del_Het_2067 and clones DJ1_X1-5del_2860/2872/2876 as a consequence of two successive rounds of editing to generate homozygous lines (compare Supplemental Fig. 10). Different color bars indicate unique or shared SNVs/Indels between different cell lines. Dark red indicates unique SNVs/Indels in edited lines, light red indicates unique SNVs/Indels in EWT lines, light blue indicates shared SNVs/Indels only found on EWTs/1st editing (DJ1/PARK7), green indicates shared SNVs/Indels between edited cell lines/2^nd^ editing (DJ1/PARK7) and EWTs/1st editing (DJ1/PARK7) and dark blue indicates shared SNVs/Indels only found in edited cell lines/2^nd^ editing (DJ1/PARK7).

To comprehensively characterize the genomic variability within the iSCORE-PD collection, we performed whole-genome sequencing (WGS) on the majority of cell lines of the collection (n = 61; complete WGS data is available at https://www.amp-pd.org/) and developed a novel analysis pipeline (Figure 4B). Using DeepVariant (https://github.com/google/deepvariant)^79^ for variant calling and Glnexus for joint-genotyping^80^, we mapped all cell line-specific variants (SNVs and indels) relative to the reference genome (GRCh38/hg38). Each genome-edited line was compared to the parental WIBR3 genome (see materials and methods for details) to identify unique variants for each clone (Figure 4B). This analysis revealed that the genome-edited cell lines in the iSCORE-PD collection, including non-edited controls (WIBR3_EWTS1-8), carry an average of 216.1 ± 55.5 (mean ± SD) SNVs/indels (Figure 4C and Supplemental Table 5). Our analysis pipeline robustly detected unique and shared variants indicated by the consistent identification of all but one (WIBR3_DNAJC6_FS_FS_H10_1) engineered mutation in the clonal cell lines (Supplemental Table 5 and 6). Of the identified SNVs/indels other than the targeted mutations, 1.3 ± 1.3 (mean ± SD, excluding synonymous mutations) variants per cell line were localized to protein coding exons (Supplemental Table 6). Importantly, protein coding SNVs in only five genes (including synonymous variants as described in detail below) were shared among multiple correctly genome-edited cell lines. Notably, the number of unique SNVs/indels in each clonal cell line showed a positive correlation with passage number (R² = 0.3614) (Figure 4D), indicating that WIBR3 cells acquire an average of 21.74 (95% CI: 14.14 to 29.34) mutations per passage during regular cell culture. This rate aligns with previously reported numbers for hPSCs^34^. Importantly, SNV/indel numbers in the non-edited control lines (EWT_S1_8) showed similar trends to those of the other genome-edited cell lines (Figure 4D), indicating that the editing process had minimal impact on the overall number of SNVs or indels per cell line.

#### Common genetic variants are rare and, when controlled for do not confound the phenotypic analysis

An important question is whether genome editing introduces common, non-random genetic variation into engineered cell lines. To address this, we analyzed all clonal cell lines isolated from each targeting experiment to edit a specific PD-associated mutation (referred to as editing group) to identify shared SNV/indel among the cell lines in this editing group. For some editing groups (*SNCA*-A53T, *DNAJC6*, *SYNJ1*, and *VPS13C* W395C), we found no shared SNVs/indels among the cell lines, apart from the engineered mutation itself (Figure 4E, Supplemental Figure 18H, I and K, Supplemental Table 5). However, in the remaining groups (*LRRK2*, *SNCA*-A30P, *PRKN*, *PINK1, DJ1/PARK7*, *ATP13A2*, *FBX07, VPS13C* A444P and *GBA1*), we identified some shared SNVs/indels between cell lines (Figure 4F-G, Supplemental Figure 18 and Supplemental Table 5).

As outlined above, two primary sources of genetic variability in genome-edited clonal lines are: (1) SNVs/indels that arise within the founder cell population prior to editing and become fixed due to targeting associated clonal expansion (founder effect), and (2) non-random genome editing-associated off-target effects. The high number of shared SNVs/indels (up to 215) between individual clonal cell lines strongly suggests that the founder effect is the predominant source of common variants. If this hypothesis is correct, similar SNVs/indels should be shared between edited clones and non-targeted controls from the same targeting experiment. Indeed, we observed a significant overlap of SNVs/indels between correctly targeted clones and non-targeted controls derived from the same experiments in the *SNCA*-A30P and *GBA1* editing groups (Figure 4F, Supplemental Figure 18L). Supporting the founder effect, clones with shared SNVs/indels exhibited significantly fewer unique variants (Figure 4F, e.g., comparing EWT_S3 and SNCA-A30P_E1-3). Together, these data suggest that the shared SNVs/indels represent a subset of the variations typically acquired during cell culture rather than additive editing-mediated variability.

Of note, all initially analyzed DJ1/PARK7 homozygous clones (WIBR3_DJ1_X1-5DEL_2860/2872/2876) share most of their SNVs/indels with the heterozygous WIBR3_DJ1_X1-5DEL_Het_2067 cell line. This is a direct consequence of the targeting strategy, as these homozygous clones were generated through two successive rounds of editing (Supplemental Figure 10). In this approach, the second clonal editing step propagates the genetic variation present in the heterozygous parental line (Figure 4A,G). To account for the potential impact of these shared variants on phenotypical analyses, we screened for an additional homozygous clone that was generated in a single targeting step (WIBR3_DJ1_X1-5DEL_6235). Additionally, we included three homozygous DJ1/PARK7 clones (WIBR3_DJ1_EX1-5DEL_6348/6390/6407), which were generated by retargeting a second heterozygous cell line (WIBR3_DJ1_X1-5DEL_2046) that did not share SNVs/indels with the previously described homozygous clones (WIBR3_DJ1_X1-5DEL_2860/2872/2876). As there is currently no evidence that heterozygous genotypes confer an increased risk of developing PD^81^, we included several heterozygous DJ1/PARK7 lines as experimental controls (WIBR3_DJ1_X1-5DEL_Het_2036/2038/2046/2051/2067) that should account for the genetic variability of the homozygous targeted DJ1/PARK7 clones (Supplemental Figure 10).

Although most common SNVs/indels are non-coding, we analyzed the shared variants that affect protein-coding sequences. As summarized in Supplemental Table 6, we identified heterozygous SNVs in the coding sequence of five genes that were shared across multiple clonal lines within specific editing groups and could affect protein function: (1) *SLC25A51* (nonsense mutation) in WIBR3_SNCA-A30P clones A2-3 and F12-1; (2) *SEPTIN10* (non-synonymous mutation) in WIBR3_DJ1_X1-5DEL_Het_2067, WIBR3_DJ1_X1-5DEL_2860, 2872 and 2876 clones; (3) *SLC35A2* (non-synonymous mutation) in WIBR3_VPS13C_A444P_Homo_C8_2 and WIBR3_VPS13C_FS_Homo_H3_1; (4) *SLC9A4* (synonymous) in WIBR3_LRRK2_G2019S_5_Het, and WIBR3_LRRK2_G2019S_6_Het; and (5) *GUCA2B* (synonymous) in WIBR3_VPS13C_A444P_Homo_C8-2 and WIBR3_VPS13C_A444P_Homo_H3-1. Consistent with a founder effect, the SNV in SLC25A51 was also detected in non-targeted control clones from the same editing experiment (EWT_S1 and EWT_S2), and the SNV in *SEPTIN10* was already present in the heterozygous WIBR3_DJ1_X1-5DEL_Het_2067 parental clone (Figure 4G). A full description of all SNVs/Indels in the protein coding region (splice sites, promoter region, introns, coding exons, 5’ UTR and 3’UTR) or each cell line in the iSCORE-PD collection is provided in Supplemental Table 7. In no case are unintended variants affecting protein coding found in all clones within any specific editing group; thus, they are unlikely to confound phenotypic interpretation when all clones for a given gene are assayed for PD-related pathologies alongside proper control cell lines.

#### Off-targets are rare in CRISPR/Cas9 edited and absent in prime edited iSCORE-PD clones

Genome editing can induce unintended off-target mutations^32,33,78^; however, the frequency and relevance of these mutations for genetically engineered hPSC-based disease models remains unclear. The above analysis indicates that the vast majority of genetic variability observed in the iSCORE-PD collection is driven by the subcloning process of cells that have spontaneously acquired mutations. Nonetheless, we cannot exclude the potential contribution of off-target effects mediated by CRISPR/Cas9 or prime editing. To investigate potential off-target effects, we used Cas-OFFinder^82^ (https://github.com/snugel/cas-offinder) with a relaxed threshold allowing up to five mismatches to generate a comprehensive list of predicted off-target sites for all gRNAs and pegRNAs employed in generating the iSCORE-PD lines. We then identified all SNVs/indels within a 100 bp window surrounding these predicted off-target sites for each genome-editing experiment. Despite the low-stringency threshold, this analysis identified only five SNVs/indels near potential off-target sites across 54 assessed cell lines (Supplemental Table 8). Among these, we considered SNVs/indels at three off-target sites in four cell lines (WIBR3_DJ1_X1_5DEL_2860, WIBR3_DJ1_X1_5DEL_2872, WIBR3_FBXO7_FS_A3_1, WIBR3_PINK1_Q129X_C4_1) as genuine off-target modifications. Consistent with a CRISPR/Cas9-mediated cleavage pattern, these modifications were located 2–5 bases upstream of the NGG protospacer adjacent motif (PAM) (Figure 5A-C, Supplemental table 8). All off-targets resulted in heterozygous modifications. Notably, only one instance of an off-target modification was shared across two cell lines (WIBR3_DJ1_X1_5DEL_2860, WIBR3_DJ1_X1_5DEL_2872). While the number of genuine off-target events was low even in double strand break-based edited clones (4 in 41 analyzed cell lines), it is important to note that no off-targets were detected in prime-edited cell lines. This is consistent with the above-described observation that prime editing induced less SNVs and LOH at the editing site. Thus, off-target effects can occur in genome-edited cell lines, particularly when using the CRISPR/Cas9 system, however they are not the primary driver of genetic variability observed in gene-edited cell lines.

**Figure 5.**
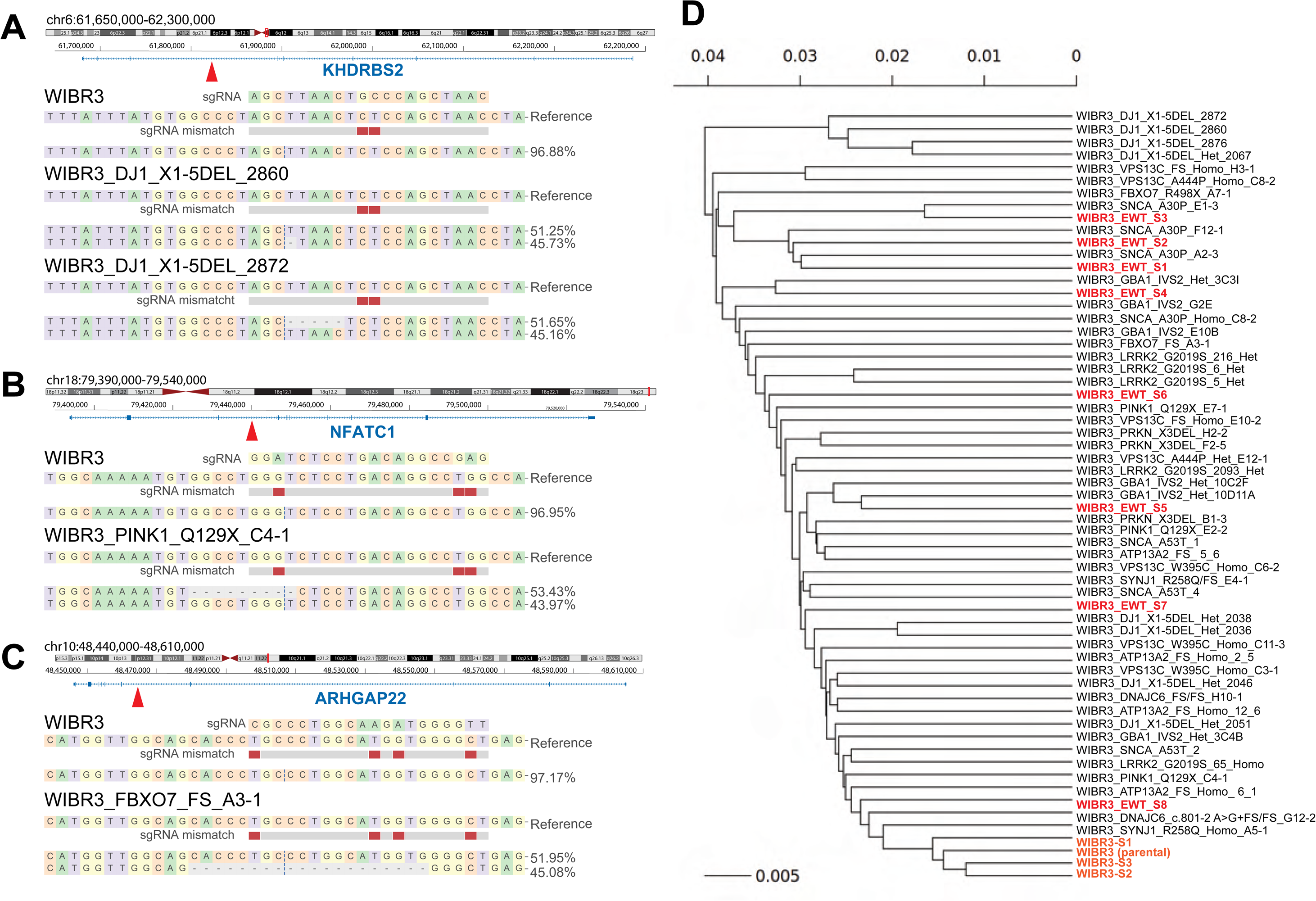
Off-targets and phylogeny. Analysis of off-targets was performed on all gene-edited cell lines and EWT_S1-5. (A-C) NGS results of predicted off-target loci (compare Supplemental Table 8) and reference WIBR3 (parental) showing genomic location and nearest gene. All off-target events described are heterozygous, intronic and non-coding. (D) Phylogenetic relationship of all analyzed cell lines in iSCORE-PD. Cell lines displayed in orange represent WIBR3 (parental) + WIBR3-S1,2,3 subclones and light red represents untargeted EWT_S1-8 cell lines.

Our analysis of WGS data clearly demonstrated that the genetic variation between genome-edited cell lines is very small compared to inter-individual variation in classical hPSC-based disease models^83^, where patient derived cell lines are compared to those from unrelated healthy individuals. However, since it is challenging to predict how the remaining variation could impact cellular phenotypes, it would be desirable that the control cell lines carry a comparable genetic variation. To evaluate how well the control lines (WIBR3_EWT_S1-8) represent the genetic variation within the iSCORE-PD collection, we computed the phylogenetic relationship of all gene edited cell lines (Figure 5D). Consistent with the results described above, correctly targeted clones were most closely related to untargeted controls from the same experiment (e.g., *SNCA*-A30P and EWT_S1-3), suggesting that control cells derived from the same targeting experiment are best to represent the genetic variability of the edited clones. Importantly, this analysis revealed that WIBR3_EWT_S1-8 are genetically distributed across all gene edited cell lines, indicating they cover the genetic variability of the entire iSCORE-PD collection. This phylogeny provides a systematic strategy to select the most appropriate controls for a given experiment based on the closest genetic correlation between controls and gene edited cell lines.

## Discussion

Advances in population genetics and sequencing technologies have greatly enhanced our understanding of the genetic architecture of complex diseases, leading to the identification of numerous genetic variants linked to the development and progression of diseases such as PD. However, revealing the functional role of these variants within a genetically diverse population remains a significant challenge. To overcome this limitation, we have generated a collection of isogenic hESC lines that carry monogenic or high-risk PD-associated mutations. Like the development of inbred animal models, which have proven instrumental in establishing robust genotype-phenotype correlations and have enabled the comparison of phenotypes across research groups, our isogenic cell line collection offers the opportunity to directly compare the phenotypic effect of PD-associated mutations in a genetically controlled system across genes and laboratories.

The establishment of an isogenic cell line collection involves two components, both crucial for the effective implementation of this approach: (1) a thorough characterization of the parental hPSC line and (2) development of a genome editing platform enabling the efficient engineering of genetic alterations similar to those found in patients. Regarding the hPSC line, we have conducted a comprehensive analysis of the parental WIBR3 hESC line and demonstrated that its genomic integrity can be sustained over extended periods in culture. We also show that WIBR3 cells are amenable to multiple rounds of clonal expansion and genome engineering. In addition, high-density genotyping and long-read sequencing show that the WIBR3 cell line does not carry major structural or genetic alterations impacting genes with known relevance to PD. Importantly, we demonstrate that WIBR3 cells can efficiently generate PD-relevant cell types *in vitro* using established differentiation protocols. Thus, WIBR3 cells are a highly characterized female reference hESC line, providing a valuable complement to existing hPSC lines for modeling neurodegenerative diseases.

To generate this collection, we established CRISPR/Cas9, TALEN and prime editing pipelines in hPSCs, enabling the highly efficient and multiplexed introduction of a broad range of disease-associated genetic alterations, ranging from heterozygous and homozygous single nucleotide variants to large structural genomic deletions. During the process of establishing this collection, we made several key observations. Notably, we recognized that all genome editing approaches necessitate a comprehensive quality control (QC) process beyond the validation of the intended genomic modification. Consequently, all the hESC lines described underwent a rigorous quality control procedure, which included the validation of pluripotency and the exclusion of karyotypic and structural aberrations using standard aCGH arrays, high-density genotyping arrays and zygosity analysis at the targeted genomic locus.

Given the observed high level of genetic instability, we decided to investigate the validity of the concept that genome editing can be used to generate isogenic cells that differ exclusively at the intended editing site. Using WGS, we comprehensively assessed the genetic variation within the iSCORE-PD collection and demonstrated that genetic variation between genome-edited cell lines is neglectable compared to inter-individual variation in classical hPSC-based disease models^83^, where patient-derived cell lines are compared to those from unrelated healthy individuals. However, the potential impact of specific variants on cellular and disease phenotypes remains unpredictable even in isogenic experiments and can pose significant problems when comparing just a single pair of genome-edited cell lines.

Our analysis revealed two major findings. First, perfect isogenic cell lines do not exist in *in vitro* cellular systems due to genetic variation, introduced by both cell culture and genome editing. Second, the vast majority of this genetic variation in genome-edited hPSCs arises from preexisting variants in the parental founder cell line acquired during routine cell culture, which become fixed through a founder effect during the clonal expansion process of genome editing (Figure 4A). The observation that many of the shared variants in genome edited cell lines are already present in the parental cell population has three important implications for the use of genome edited cell lines in disease modeling: (1) The best approach to control for this genetic variation is to include multiple independently targeted disease-associated cell lines and controls. (2) Untargeted, clonally derived cells from the same targeting experiment are the best controls, as they most accurately represent the genetic variability of the edited clones. (3) Multi-step cloning strategies carry the highest risk of generating lines with shared variants, as every consecutive editing step propagates the genetic variation present from the preceding manipulation. Together, these findings emphasize the importance of carefully designing genome-editing experiments to account for and mitigate the effects of shared genetic variants on downstream phenotypic analyses.

Finally, by analyzing mutations that are unique in each cell clone we estimate that with each passage hPSCs acquire about 20 additional mutations. Previously, it has been suggested that one way to distinguish mutation-specific phenotypes from alterations coming from unintended genetic variation is to revert the genome-edited cell lines to the wild-type genotype^84^ in a second step. This strategy is highly useful to validate a specific mutation-associated phenotypes. However, our analysis suggests that relying only one cell clone when identifying novel or subtle phenotypes might be insufficient to for account for the genetic variation that is introduced de novo by the continues culturing of cells. Moreover, the mutation correction approach bears complications when comparing phenotypes across different disease-causing mutations as each carries numerous cell lines specific mutations.

In addition, we address the outstanding question whether CRISPR/Cas9 or prime editing is better suited for generating genome-edited hPSC collections. A key finding across the derivation of all cell lines in the iSCORE-PD collection was that the frequency of karyotypic and structural aberrations, as well as LOH of the edited locus, was more frequent in CRISPR/Cas9 than in prime edited cell lines. Moreover, we only detect off-targets in cell lines generated using CRISPR/Cas9. This is the first time that this has been formally reported across a large cohort of gene edits combined with a detailed genotyping approach. This finding is consistent with CRISPR/Cas9-based genome editing introducing a potentially genotoxic double strand break (DSB) at the target site to insert genetic modifications that is frequently processed through complex DNA repair reactions. Instead, prime editing only introduces single-strand DNA nicks, a genetic insult that is more readily repaired by a cell without mutations or genomic rearrangements, driving the repair outcome toward the intended genetic modification^85^.

Furthermore, our analysis confirmed that off-target effects, though rare, can occur in genome-edited cell lines. While the number of genuine off-target events was low even in CRISPR/Cas9 edited clones (4 in 41 analyzed cell lines), it is important to note that no off-target effect was detected in prime edited cell lines. Together, these results confirm our previous observations, that prime editing has substantial advantages over CRISPR/Cas9-based approaches for introducing point mutations and small structural modifications in hPSCs^70^. Furthermore, we strongly recommend including a zygosity analysis, specifically to exclude LOH at the target locus, as a critical step in the quality control pipeline of genome engineered hPSCs.

Given that any genetic alteration induced by cell culture or genome editing can impact the biological properties of hPSCs and disease phenotypes, the in-depth analysis of the genomic integrity and variability of the iSCORE-PD collection emphasizes the necessity for a comprehensive quality control in genome engineering^86^. Considering that clonally derived cell lines are susceptible to cell culture-induced genetic drift and acquire additional genetic and epigenetic alterations over time, we advocate using multiple independently gene-edited clonal lines for each genotype to account for this variability. In addition, we recommend using low-passage number cell lines and performing routine quality control analysis to detect culture induced genetic aberrations, including whole genome sequencing. This approach ensures a robust assessment of disease-relevant phenotypes in vitro, acknowledging the potential variability that may arise during prolonged cell culture and genome editing processes.

While perfect isogeneity remains elusive, the observed genetic variations were minimal compared to those typically found in classical hiPSC experiments comparing cells from patients with those from unaffected individuals. Predicting the impact of this variation on cellular and disease phenotypes is challenging. However, as the majority of these variations are random and predominantly affect non-coding regions of the genome — similar to somatic mutations found ubiquitously across normal tissues — we predict that most of the observed variation is unlikely to strongly impact cellular fitness or disease-associated phenotypes^77^. Therefore, we believe that such variants do not diminish the value of genetically controlled hPSC collections like iSCORE-PD in disease research. Importantly, we provide a roadmap for effectively managing these variations through stringent quality control measures and careful experimental design.

The cell lines described here currently focus on coding risk variants with large effect size linked to monogenic PD^4^. We envision that we and other researchers can expand this collection to eventually incorporate GWAS-identified risk variants with lower effect size. Such an expansion could provide functional insights into how these primarily non-coding sequence variants affect similar cellular and molecular pathways as implicated in monogenic PD. To facilitate such efforts, all generated cell lines will be made available with the support of the Aligning Science Across Parkinson’s (ASAP) initiative and the Michael J. Fox Foundation (MJFF) through the WiCell Research Institute. We anticipate that the subsequent biological analysis of this comprehensive collection and its future expanded forms, involving numerous research groups with diverse expertise, can provide a unified understanding of how genetic risk variants functionally contribute to the pathogenesis of PD. We predict that this collaborative effort has the potential to accelerate the development of novel therapeutic strategies for PD.

## Limitations of this study

Each genome is unique, carrying a distinct combination of sequence variants and genetic alterations that can influence the development and pathology of complex diseases such as PD. Consequently, there is no single cellular model that can fully recapitulate all the molecular and cellular features of such disorders. Given this limitation, it will become necessary to expand the described approach to include additional cell lines with diverse genetic backgrounds to fully dissect the pathobiology of PD. Moreover, our work specifically addresses the genetic variation associated with genome editing and *in vitro* culture of hPSCs. However, it is widely recognized that additional epigenetic modifications acquired during this process can affect the phenotypical analysis of *in vitro*-derived cell types, irrespective of the disease genotype. While using multiple independently edited clonal cell lines alongside continuous quality control measures can mitigate many random genetic alterations, we cannot entirely exclude remaining systematic genetic and additional epigenetic modifications associated with specific gene editing approaches.

## Methods

### hPSCs culture

hESCs were maintained on irradiated or mitomycin C-inactivated mouse embryonic fibroblast (MEF) monolayers as described previously^70^ with daily changes of hESC media (Dulbecco’s Modified Eagle Medium/Nutrient Mixture F-12 (DMEM/F12; Thermo Fisher Scientific) supplemented with 15% fetal bovine serum (Hyclone), 5% KnockOut Serum Replacement (Thermo Fisher Scientific), 1 mM glutamine (Invitrogen), 1% nonessential amino acids (Thermo Fisher Scientific), 0.1 mM β-mercaptoethanol (Sigma) and 4 ng/ml fibroblast growth factor (FGF) (Thermo Fisher Scientific/Peprotech), 1×Penicillin-Streptomycin (Thermo Fisher Scientific). All hESCs cultures were maintained in a cell culture incubator under low oxygen conditions (95% CO2, 5% O2). Cultures were passaged as aggregates every 5-7 days using a collagenase IV solution (Gibco) to detach hESC colonies. All cell lines are tested routinely for mycoplasma. Detailed protocols for culturing of hESCs on MEF feeders can be found on protocols.io (https://doi.org/10.17504/protocols.io.b4pbqvin ; https://doi.org/10.17504/protocols.io.b4msqu6e). All hESCs cultures were adapted to feeder-free culture conditions before starting *in vitro* differentiation experiments. hESCs were maintained on geltrex/matrigel coated plates in mTeSR plus medium (Stem Cell Technologies) in a cell culture incubator under low oxygen conditions (95% CO2, 5% O2) as described previously^70^. Cells were passaged regularly as aggregates either manually or using ReLeSR (Stem Cell Technologies) to detach hESC colonies. Detailed protocols for feeder-free culturing of hPSCs can be found on protocols.io (https://doi.org/10.17504/protocols.io.b4mcqu2w).

### Collecting cell pellets for DNA and RNA extraction

hESCs colonies cultured on MEFs were harvested by collagenase IV and washed twice through an 80 µm cell strainer to further remove MEFs. Collected colonies were pelleted by centrifugation and snap frozen in liquid nitrogen.

### Array genotyping and data processing

Genomic DNA was isolated from cell pellets using the DNeasy Blood & Tissue Kit (QIAGEN; 69504). Genotyping was performed using the Neuro Booster Array (NBA) with best practices guidelines for the Infinium Global Diversity Array^87^. Genotyping data was processed using GenomeStudio (RRID:SCR_010973) and subsequent genotype calls, B-allele frequency and LogR ratio values were used for genomic integrity assessments. When cell lines carrying a genomic edit were present on the NBA, genotype calls were compared to confirm the edit. Genome-wide genotyping calls were compared with the PacBio HiFi WGS variants to assess large genomic events across the two data types using PLINK (v1.9, RRID:SCR_001757)^88^. The B-allele frequency and LogR ratio values were processed and plotted using the GWASTools package in R (v3.6.1, https://www.r-project.org/, DOI: 10.18129/B9.bioc.GWASTools)^89^.

### Long-read sequencing and data processing

#### Oxford Nanopore Technologies DNA extraction, library preparation, and sequencing

Ultra-high molecular weight DNA (UHMW) was extracted from the WIBR3 (parental) hESC line (5 × 10^6^ cells) following the Circulomics/Pacific Biosciences (PacBio) UHMW DNA Nanobind Extraction protocol (Circulomics/PacBio, no longer available) with the Nanobind CBB Kit (PacBio, SKU 102-301-900) and the UHMW DNA Aux Kit (Circulomics/PacBio, NB-900-101-01, no longer available). The extracted DNA was checked using the Qubit dsDNA BR assay (Invitrogen, Q32850) to ensure proper extraction occurred. The extracted UHMW DNA was then taken straight into library preparation for sequencing using Oxford Nanopore Technologies (ONT) SQK-ULK001 Kit and the Nanobind Ultra Long Library Preparation Kit (Circulomics/PacBio, NB-900-601-01, no longer available). The library was split into 3 tubes of 75 µl each, and each tube loaded on a flow cell. After 24 hours, 75 µl of the sequencing library was pulled out of each flow cell and reloaded on a fresh flow cell. This process was repeated one more time for a total of 9 separate R9.4.1 PromethION flow cells.

#### Pacific Biosciences DNA extraction, library preparation, and sequencing

High molecular weight (HMW) was extracted using PacBio’s Nanobind CBB Kit (Pacbio, 102-301-900) from 2 × 10^6^ cells with the Nanobind adherent cultured cells protocol. After extraction, DNA concentration was quantified using the Qubit dsDNA BR assay (Invitrogen, Q32850), sized with a Femto Pulse System (Agilent, M5330AA), and size selected with the PacBio SRE Kit (Pacbio, SKU 102-208-300). Following quality control, the extracted DNA was sheared to a target size of 18-20 kb using the Megaruptor 3 (Diagenode, B060100003). After confirmation of correct sizing, the library preparation was performed SMRTbell prep kit 3.0 (PacBio, 102-141-700) with a PEG wash. The library was sequenced on a Revio flow cell with a 24 h movie time.

#### Long read sequencing Data Analysis

ONT sequencing runs were basecalled on NIH’s HPC (Biowulf) using Oxford Nanopore’s Guppy (v6.1.2, RRID:SCR_022353) in super accuracy mode with the *dna_r9.4.1_450bps_modbases_5mc_cg_sup_prom.cfg* configuration file and the –bam_out option to preserve methylation tags. The basecalled bams were then converted to fastqs using Samtools (v1.17, RRID:SCR_002105)^90^ (samtools fastq -TMm, Ml) and mapped to hg38 using Minimap2 (v2.24, RRID:SCR_018550)^91^ with ONT flags. Data from all flow cells was merged after mapping using samtools (v1.17, RRID:SCR_002105). Then, we used PEPPER-Margin-DeepVariant (v.0.8, https://github.com/kishwarshafin/pepper)^92^ to call small variants (<50bp) and phase our variant calls and alignments. We then used our phased alignment, to produce haplotype-specific methylation calls using Modbamtools (v0.4.8, https://rrazaghi.github.io/modbamtools/)^93^ and Nanopore’s modbam2bed (https://github.com/epi2me-labs/modbam2bed). Lastly, structural variants (SVs) were called using Sniffles2 (v2.2, RRID:SCR_017619)^94^ with default settings. PacBio Revio HiFi data was processed according to general best practices. Data was mapped using Minimap2 (v2.24, RRID:SCR_018550) using PacBio flags. Small variant calls generated by Clair3 (v1.0.4) (https://github.com/HKU-BAL/Clair3) with PacBio flags and SV calls were generated by Sniffles2 (v2.2, RRID:SCR_017619)^94^. Small variants were filtered for DP>15 and GQ>20 using bcftools (v1.17, RRID:SCR_005227)^90^ and annotated with ANNOVAR (v.2022-06-08, RRID:SCR_012821)^95^ to assess the presence of potential pathogenic variants. In addition, Alzheimer’s disease^96^ and Parkinson’s disease genetic^49^ risk scores (excluding UK Biobank summary statistics) were calculated to assess the cumulative risk score using plink (v2.0, RRID:SCR_001757) for disease and compared with participants from the UK Biobank diagnosed with AD and PD^97^. Only SV calls labeled as “PASS” were kept for both ONT and PacBio data. The “PASS” SV calls were then annotated with ANNOVAR (v.2022-06-08, RRID:SCR_012821)^95^ and coding variants were subset. Then, we used Truvari (v4.4.0)^98^ ((https://github.com/ACEnglish/truvari) to merge structural variant calls between the ONT and PacBio datasets both for all variants as well as only coding variants. Numbers on variant type distribution were generated using SURVIVOR (v1.0.7, RRID:SCR_022995). Lastly, the SV overlaps were further annotated using SVAnna^52^ (v1.0.4) (https://github.com/TheJacksonLaboratory/SvAnna with the phenotype terms HP:0002180 (neurodegeneration) and HP:0000707 (abnormality of the nervous system). SVs of interest were plotted using samplot (v1.3.0) (https://github.com/ryanlayer/samplot).

#### WGS analysis pipeline

DNA samples were sequenced with Illumina short-read WGS at Psomagen (Rockville, MD), with a mean coverage of 30x. Data was processed using standard GP2 WGS pipelines. In brief, 150 bp paired-end reads were aligned to the human reference genome (GRCh38 build) using BWA-mem (https://github.com/lh3/bwa) following the functional equivalence pipeline^99^. Sample processing and variant calling were performed using DeepVariant v.1.6.1^79^. Joint-genotyping was performed using GLnexus v1.4.3 with the preset DeepVariant WGS configuration^80^. A detailed description of the WGS pipeline can be found at https://github.com/GP2code/releases/tree/main/BETA-APR2022/wgs_var_calling and https://github.com/GP2code/GP2-WorkingGroups/tree/main/MN-DAWG-Monogenic-Data-Analysis/Terra_wdl/variant_calling/deepvariant.

#### hPSC edited cell line SNV pipeline analysis

To identify common and specific SNVs for each group of cell lines modified by a set of CRISPR/Cas9 or prime editing reagents, the identified polymorphisms identified with the WGS pipeline were first filtered to only consider calls with a GLNexus quality score greater or equal to 30. Then, to identify SNVs/indels specific to only one edited group, the SNVs were further filtered to remove any calls found in any of the other cell lines in the collection. The remaining SNVs are what account for each editing group’s specific SNVs/indels and were used to determine unique and shared variants within each edited group and to further characterized their contribution to coding or non-coding region as well as their effect on the coding sequences by leveraging tools within the BioConductor DeepVariant package^100^ against the TxDb.Hsapiens.UCSC.hg38.knownGene_3.18.0 human transcript annotation package (based on the UCSC hg38 genome based on the knownGene table). A detailed description of the pipeline can be found here (https://github.com/hockemeyer-ucb/pd-sv-analysis). The group unique SNVs/indels were used to generate a phylogeny tree using the BioConductor package fastreeR^101^ and visualized using the R package ape^102^.

#### Off-target analysis

To test whether some of the group specific SNVs could be link to potential off-target effect triggered by the CRISPR/Cas9 or prime editing reagents, all putative off-target sites were identified using Cas-OFFinder^82^ (https://github.com/snugel/cas-inder) and used to identified nearby SNVs/indels that could have been the results of the editing strategies. A detailed description of the pipeline can be found here (https://github.com/hockemeyer-ucb/pd-sv-analysis).

### Molecular cloning

Molecular cloning was carried out as described previously^70^ following standard cloning protocols (https://www.cshlpress.com/pdf/sample/2013/MC4/MC4FM.pdf)^103^. As described^85^, pegRNA plasmids for prime editing were cloned by ligating annealed oligonucleotide pairs (Supplemental table 9) into the BsaI-digested pU6-peg-GG-acceptor (pU6-pegRNA-GG-acceptor was a gift from David Liu. RRID:Addgene_132777; http://n2t.net/addgene: 132777; RRID:Addgene_132777). Prime editing nicking guide plasmids (ngRNAs) were cloned by ligating annealed oligonucleotide pairs (Supplemental table 9) into the BsmBI-digested pBPK1520 plasmid (BPK1520 was a gift from Keith Joung. Addgene#65777; http://n2t.net/addgene: 65777; RRID:Addgene_65777)^104^. For CRISPR/Cas9 based genome editing, the Cas9 expressing gRNA plasmids were cloned by ligating annealed oligonucleotide pairs (Supplemental Table 9) into the BbsI-digested px330-GFP (RRID:Addgene_97084)^40^ or px330-mCherry (RRID:Addgene_98750) as described previously^40^. For TALEN mediated genome editing, we used previously described heterodimeric TALEN pairs to insert the G2019S into the LRRK2 gene^70^. Sequence information for all oligonucleotides (Integrated DNA Technologies, IDT) used to generate plasmids can be found in Supplemental Table 9.

### Genome editing of hESCs

As outlined in Figure 3C, genome editing of WIBR3 hESCs was performed using either plasmid or ribonucleoprotein particle (RNP) based CRISPR/Cas9 or prime editing approaches as described previously^70^ using the following procedures:

#### Nucleofection

hESCs cultured on MEFs were pre-treated with 10 µM ROCK inhibitor (Y27632, ToCris) 1-day before nucleofection (2-3 hours at a minimum is recommended). Cells were collected by collagenase IV (Thermo Fisher Scientific) followed by Accutase (Thermo Fisher Scientific) to dissociate hESCs into a single cell solution. 5 × 10^5^ to 1 × 10^6^ cells were resuspended in 20µL of nucleofection solution (P3 Primary Cell 4D-Nucleofector™; Lonza) and nucleofected (Lonza 4D nucleofector TM Core + X Unit, program CA-137) using the following genome editing reagents for the corresponding edits described in Figure 3: *(1)* Plasmid based CRISPR-Cas9 facilitated HDR: 200 ng gRNA plasmids (px330-GFP), 700 ng ssODN. *(2)* Plasmid based dual CRISPR: 500 ng 3’-gRNA plasmid (px330-GFP) and 500 ng 5’-gRNA plasmid (px330-mCherry (RRID:Addgene_98750)). *(3)* TALEN facilitated HDR: 100 ng LRRK2-TALEN-TA01L and 100 ng LRRK2-TALEN-TA03R, 700 ng ssODN, 100 ng pEGFP-N1 (RRID:Clontech_6085-1). *(4)* Plasmid based prime editing: 500 ng pCMV-PE2-GFP (a gift from David Liu, RRID:Addgene_132776)^85^, 330 ng pU6-pegRNA (RRID:Addgene_132777) and 170 ng pBPK1520-ngRNA (RRID:Addgene_65777). *(5)* RNP-based CRISPR-Cas9 facilitated HDR: 80 pmol purified Cas9 protein (QB3 Macrolab, UC Berkely), 300 pmol chemically modified synthetic sgRNA (Synthego) and 100 pmol ssODN HDR template. *(6)* RNP-based dual CRISPR: 80 pmol purified Cas9 protein, 150 pmol of each chemically modified synthetic 3’-sgRNA and 5’-sgRNA. *(7)* RNP-based CRISPR-Cas9 facilitated HDR with competing templates: 80 pmol purified Cas9 protein, 300 pmol chemically modified synthetic sgRNAs, 50 pmol ssODN HDR template carrying PD mutation and 50 pmol ssODN HDR template carrying a synonymous mutation. *(8)* RNA-based prime editing: 4 μg in vitro transcribed nCas9-RT mRNA, 100 pmol chemically modified synthetic pegRNA (IDT or Synthego) and 50 pmol chemically modified synthetic ngRNA (Synthego). Detailed protocols can be found on protocols.io (dx.doi.org/10.17504/protocols.io.e6nvwkkewvmk/v2).

#### Editing pipelines

We used two different editing pipelines termed *Pipeline A* and *Pipeline B* to generate the iSCORE-PD collection (Figure 3A,B). The editing pipeline used to create each cell line in the iSCORE-PD collection is included in Supplemental Table 3.

*Pipeline A* utilizes FACS-enrichment post-nucleofection to purify effectively transfected cells, followed by clonal expansion and genotyping to allow the isolation of clonal, correctly targeted lines (estimated time for the editing pipeline is 30-35 days). Following nucleofection, the hESCs are plated on MEFs in 10 µM ROCK inhibitor (Y27632, ToCris) containing hESC media (previously described in hPSC culture section) at high density (1 nucleofection/1 well 6 well plate). 48-72 h after nucleofection, Accutase-dissociated single cells are FACS-sorted for the expression of the respective fluorescent marker protein and either directly used for bulk NGS based validation of the desired genome modification or subsequently plated at clonal density (250 to 350 cells/cm2) on MEFs in hESC media supplemented with 10 µm ROCK inhibitor (Y27632, ToCris) for the first 24 hr. Individual colonies are picked and grown 7 to 14 days after electroporation. Correctly targeted clones were subsequently identified by Sanger or NGS sequencing. A detailed protocol can be found on protocols.io (dx.doi.org/10.17504/protocols.io.b4piqvke).

*Pipeline B* (high throughput hPSCs genome editing) involves low cell number nucleofection, limited dilution, and NGS-dependent genotyping to identify desirable edits in a 96-well plate system. This integrated workflow allows the efficient isolation of correctly edited clonal cell lines, even at low frequency, within a shorter time frame compared to previous approaches (21-35 days). As described previously^70^, the nucleofected cells are directly seeded onto MEFs in 96-well plates, at seeding densities of 1000 cells/plate in hPSCs media containing 10 µm ROCK inhibitor (Y27632, ToCris). After individual colonies appear around day 14, plates are duplicated for *(1)* maintenance and *(2)* DNA extraction for NGS-based identification of wells that contain cells with the desired genetic modification. To duplicate plates, cells are washed with PBS (Corning) and treated with 40µL 0.25% trypsin for 5 min at 37 C. 60µL hESC media containing 10 µM Rock inhibitor (Y27632, ToCris) is added to each well to inactivate trypsin. Cells are gently dissociated, and half (50 µL) of the cell suspension is reseeded to a new MEF containing 96-well plate pre-loaded with 100µL hPSC media containing 10 µM Rock inhibitor (Y27632, ToCris) and cultured for another 7 days with hPSC media.

#### NGS-based identification of validation of targeted clonal lines

50µL of cell suspension/well obtained during plate duplication is transferred to a 96-well PCR plate pre-loaded with 50µL 2X lysis buffer (100mM KCl, 4mM MgCl2, 0.9% NP-40, 0.9% Tween-20, 500µg/mL proteinase K, in 20mM Tris-HCl, pH 8) for DNA extraction (50 °C overnight incubation followed by 95 °C 10 min [proteinase K inactivation]). A ∼300bp genomic region covering the designed mutation is amplified (Supplemental table 9) containing NGS barcode attachment sites (GCTCTTCCGATCT) from 2ul cell lysis from each well with Titan DNA polymerase. Amplicons were purified at the UC Berkeley DNA Sequencing Facility, then i5/i7 barcoded in indexing PCR, pooled and sequenced on 150PE iSeq in the NGS core facility at the Innovative Genomics Institute (IGI). CRISPResso2 (RRID:SCR_021538)^105^ in prime editing mode was used to analyze the NGS data to identify wells containing the designed mutation, with the following criteria. Heterozygous candidates: number of reads aligned >100, 70% >mutant allele frequency >20%, indels frequency <5%; homozygous candidates: number of reads aligned >100, mutant allele frequency >70%, indels frequency <5%. Wells containing the desired editCells in those identified wells were single cell subcloned once and genotyped clonally to confirm cell line purity to ensure clonality. Detailed protocols for high throughput hPSCs genome editing (dx.doi.org/10.17504/protocols.io.b4mmqu46) and genotyping by next generation sequencing https://doi.org/10.17504/protocols.io.b4n3qvgn) can be found on protocols.io. For clarity, NGS results reported in any of the figures of this publication showcase only representative reads. Any NGS reads below 1% of the total result were removed. The full NGS report can be found with the rest of raw data files (10.5281/zenodo.14907986 or AMP-PD data repositories).

### Zygosity confirmation by SNV detection

The SNV closest to the editing site for each genetic edit was identified from the whole genome sequencing data of parental WIBR3 hESCs. A genome DNA region flanking the SNV and the editing site was amplified by PCR (Supplemental Table 9) and sequenced by Sanger sequencing or NGS. Clones showing LOH were removed from the final collection.

### Cortical spheroid differentiation

hESCs were differentiated into early cortical spheroids following an adaptation of a published protocol (doi.org/10.1016/j.tcb.2019.11.004, DOI: 10.1038/nmeth.3415)^106,107^. In brief, hESC colonies were dissociated and plated into pre-coated 6-well Aggrewell 800 plates at a concentration of 18M cells per well in hESC media with 10 µM Rock Inhibitor (ToCris). The next day (Day 1), the aggregates were removed, sedimented, and added to an ultralow adherence plate with hESC media supplemented with 5 µM Dorsomorphin (SelleckChem) and 10 µM SB431542 (SelleckChem) (media changed daily). On Day 6, media was replaced with Neural Precursor Expansion Media (Neurobasal medium + B27 supplement without vitamin A (2% vol/vol) + Penicillin-Streptomycin (100U/ml) + GlutaMAX (1% vol/vol) + HEPES Buffer (1% vol/vol) + FGF2 (20 ng/ml) + EGF (20 ng/ml)) (media is changed every day until Day 16 and then every other day until Day 25). For specific details consult published materials on protocols.io: https://doi.org/10.17504/protocols.io.5jyl8po57g2w/v1.

### Cell irradiation

hESCs were transferred from MEFs to feeder-free matrigel substrate with conditioned media for 2 weeks previous to this experiment. hESCs at 50% confluence or cortical spheroids on day 25 of differentiation were irradiated at 0, 0.5, 2, 5, and 10 Gy using a discrete cesium source. 24 hours post irradiation, cells were collected and dissociated for MULTI-Seq barcoding and sequencing. For specific details consult published materials on protocols.io (https://doi.org/10.17504/protocols.io.bp2l6xwbzlqe/v1).

### MULTI-Seq Barcoding and Single-Cell Library Preparation of Irradiated Samples

*hESCs:* Each irradiation condition was labeled with a lipid-modified barcoded MULTI-seq oligo following a previously described protocol (DOI: doi.org/10.1038/s41592-019-0433-8)^108^. In short, cells in PBS were incubated with a 1:1 molar ratio of lipid-modified Anchor Oligo:Barcode Oligo for 5 min on ice. Then an equimolar amount of lipid-modified co-anchor was added for an additional 5 min incubation on ice. Then cells were washed twice with ice cold PBS (Corning) + 1%BSA (Fisher) to sequester the anchor oligos, strained, counted, and pooled for single-cell sequencing. 10x single-cell RNA sequencing was performed according to manufacturer’s instructions using the Chromium Single Cell 3ʹ Reagent Kits v3 with Feature Barcoding Technology. For specific details consult published materials on protocols.io (https://doi.org/10.17504/protocols.io.kxygx3xzkg8j/v1.) Deconvolution of MULTI-Seq barcodes was performed as described previously^108^ using the MULTI-seq package at https://github.com/chris-mcginnis-ucsf/MULTI-seq/.

*Cortical Spheroids:* single-cell suspensions were FACS-sorted to remove debris and aggregates, then 10x single-cell RNA sequencing was performed according to manufacturer’s instructions using the Chromium Single Cell 3ʹ Reagent Kits v3, targeting 2,000 cells per irradiation condition using one 10x lane per condition. Single-cell analysis of all irradiated samples was performed using Seurat v4 (RRID:SCR_016341)^109^ according to default parameters for normalization and integration of data sets. Droplets with more than 15% mitochondrial reads detected were excluded as poor analysis candidates due to likelihood of cell death resulting in poor RNA representation. Plots were generated using ggplot2 (RRID:SCR_014601) in R.

### Dopaminergic neuron differentiation

Feeder-free adapted WIBR3 hESCs were differentiated into dopaminergic neurons as per previously reported protocols with slight modifications (DOI: 10.1016/j.stem.2021.01.004, DOI: 10.1016/j.stem.2021.01.005)^62,63^. Briefly, hESC colonies were dissociated into single cells and seeded onto matrigel coated plates at a density of 400-600k cells per well of a 6 well plate in mTeSR (Stem Cell Technologies) containing 10 µM Rock inhibitor (Y27632, ToCris). Differentiation was induced sequentially with media A - 3 days (Neurobasal media (Gibco) + N2 supplement (Gibco; 1% vol/vol) + B27 supplement without vitamin A (Gibco; 2% vol/vol) + L-Glutamine (Gibco; 2 mM) + Penicillin-Streptomycin (Gibco; 100U/ml) + SHH C25II (R&D systems; 100-200 ng/ml) + CHIR99021 (ToCris; 0.7 µM) + LDN (Stemgent; 250 nM) + SB431542 (SelleckChem; 10 µM)), B -3 days (Neurobasal media (Gibco) + N2 supplement (Gibco; 1% vol/vol) + B27 supplement without vitamin A (Gibco; 2% vol/vol) + L-Glutamine (Gibco; 2 mM) + Penicillin-Streptomycin (Gibco; 100U/ml) + SHH C25II (R&D Systems; 100-200 ng/ml) + CHIR99021 (ToCris; 7.5 µM) + LDN (Stemgent; 250 nM) + SB431542 (SelleckChem; 10 µM)), C - 3 days (Neurobasal media (Gibco) + N2 supplement (Gibco; 1% vol/vol) + B27 supplement (Gibco; 2% vol/vol) + L-Glutamine (Gibco; 2 mM) + Penicillin-Streptomycin (Gibco; 100U/ml) + CHIR99021 (SelleckChem; 7.5 µM)) and D -1 day (Neurobasal media (Gibco) + B27 supplement (Gibco; 2% vol/vol) + L-Glutamine (GIbco; 2 mM) + Penicillin-Streptomycin (Gibco; 100U/ml) + BDNF (PeProtech; 20 ng/ml) + GDNF (PeProtech; 20 ng/ml) + Ascorbic acid (Sigma; 200 µM) + Dibutyryl-cAMP (SelleckChem; 0.5 mM) + TGFꞵ3 (R&D Systems; 1 ng/ml) + CHIR99021 (SelleckChem; 3 µM)) over an 10 day period. On day 11, cells were dissociated and plated (1:2 ratio) at high density and maintained in maturation media (Neurobasal media (Gibco) + B27 supplement (Gibco; 2% vol/vol) + L-Glutamine (GIbco; 2 mM) + Penicillin-Streptomycin (Gibco; 100U/ml) + BDNF (PeProtech; 20 ng/ml) + GDNF (PeProtech; 20 ng/ml) + Ascorbic acid (Sigma; 200 µM) + Dibutyryl-cAMP (SelleckChem; 0.5 mM) + TGFꞵ3 (R&D Systems; 1 ng/ml) + DAPT (ToCris; 10 µM)) until day 16, when they were replated at the similar high density in 12 well plate and left to mature until day 24. On day 25, cells were dissociated for the final time with accutase and replated at no less than 1-2 × 10^6^ cells per well of 12 well plate and left to mature until post-differentiation experiments were carried out. For specific details consult published materials on protocols.io: https://doi.org/10.17504/protocols.io.3byl4q8yovo5/v1.

### scRNA-Seq of dopaminergic neurons - 10x Genomics library preparation

Dopaminergic neurons were harvested with Accutase on days 35-37 of culture and subsequently labeled with 10x Genomics CellPlex reagents, following the manufacturer recommendation (10x Genomics CG000391 Rev B). After labeling with Cell Multiplexing Oligos (CMOs), samples were pooled and taken for 10x Genomics library preparation, following manufacturer recommendations with target capture of 30,000 cells per 10x lane (Chromium Single Cell 3’ Reagent Kits v3.1, User Guide CG000388 Rev C).

### scRNASeq of dopaminergic neurons – Data analysis

After next-generation sequencing of 10x Genomics libraries (NovaSeq 6000), FASTQ files were processed with 10x Genomics CellRanger pipeline (v7.0.1, RRID:SCR_017344) to demultiplex and generate count matrices for each sample. Data for each sample was first filtered to remove low-quality cells (cells with fewer than 1,500 genes detected, greater than 30,000 RNA counts, and greater than 10% mitochondrial reads were removed). Filtered datasets were each processed individually with Seurat v4 (RRID:SCR_016341)^109^, using the SCTransform function for normalization and variance stabilization. Integration of the SCTransformed data was performed to generate a combined dataset of 10,097 cells.

### Microglia differentiation

To generate in vitro differentiated microglia cells (iMGs), we adapted a previously published protocol (DOI: 10.1016/j.stemcr.2017.04.023)^69^. Undifferentiated feeder free hESC colonies maintained in mTeSR (Stem Cell Technology) were seeded at low density into cell culture flasks coated with reduced growth factor matrigel (30 colonies/T75 flask (Fisher)) using manual passaging. In vitro differentiation was achieved by sequential culture of the cells in the following media: Step 1 (mTeSR (Stem Cell Tech) + 80 ng/ml BMP4 (PeProtech) - 4 days), Step 2 (StemPro-34 SFM (Gibco) + 2 mM GlutaMAX (Gibco) + 80 ng/ml VEGF (PeProtech), 25 ng/ml FGF (PeProtech) + 100 ng/ml SCF (PeProtech) - 3 days), Step 3 (StemPro-34 SFM (Gibco) + 2 mM GlutaMAX + 50 ng/ml SCF (PeProtech) + 50 ng/ml IL-3 (PeProtech) + 5 ng/ml TPO (PeProtech) + 50 ng/ul M-CSF (PeProtech) + 50 ng/ul Flt3 (PeProtech) - 9 days) and Step 4 (StemPro-34 SFM (Gibco) + 2 mM GlutaMAX + 50 ng/ml M-CSF (PeProtech) + 50 ng/ml Ftl3 (PeProtech) + 25 ng/ml GM-CSF (PeProtech) - 14 days). After ∼28 days, microglia progenitors are ready to be isolated and plated on Primaria plates (Corning) for maturation (at least 2 weeks) in microglia maturation media (Neurobasal media (Gibco) + N2 Neuroplex (Gemini; 1x final concentration) + GEM21 Neuroplex (Gemini; 1x final concentration) + 20% AlbuMAX I (Gibco; 0.2% final concentration) + NaCl (Fisher; 5M) (50mM final concentration) + sodium pyruvate 100x (Gibco’ 1x final concentration) + glutaMAX 100x (Gibco; 1x final concentration) + Penicillin-Streptomycin (Gibco; 100U/ml) + 50 ng/ml TGF-β1 (PeProtech) + 100 ng/ml IL- 34 (PeProtech) + 12.5 ng/ml M-CSF (PeProtech)). Detailed protocols for microglial differentiation can be found on protocols.io (https://doi.org/10.17504/protocols.io.4r3l22zbjl1y/v1). Microglia cells were evaluated by immunocytochemistry (https://doi.org/10.17504/protocols.io.yxmvm3146l3p/v1) and FACS-based analysis (https://doi.org/10.17504/protocols.io.81wgbxokqlpk/v1) in order to confirm expression of precursor and mature microglia markers (CD16, CD45, CX3CR1, P2RY12, CD11b, CD14, IBA1, PU.1). FACS data was analyzed using FlowJo software (version 10.8.0).

#### Immunocytochemistry

Immunocytochemistry was used to assess biomarker expression to characterize each of the cell types shown in this publication. Briefly, samples were fixed in PFA and permeabilized (0.03% triton when necessary) and blocked (BSA or serum) as required depending on the biomarker being analyzed on hESCs (OCT4 (DSHB), SSEA4 (DSHB)) or our differentiated cell cultures: dopaminergic neurons (TH (Pelfreeze), FOXA2 (R&D Systems)) or microglia (IBA1 (Abcam), P2RY12 (Sigma), CX3XR1 (Biolegend), PU.1 (Cell Signaling Tech)). Fluorochrome conjugated secondary antibodies were used to image our samples in an epifluorescence or confocal microscope. Alkaline phosphatase activity was measured using Vector® Black Substrate Kit, Alkaline Phosphatase (Vector Laboratories). Specific details on the protocol used can be found in protocols.io (https://doi.org/10.17504/protocols.io.yxmvm3146l3p/v1). For specific details on our staining of pluripotency markers in our hESCs consult: https://doi.org/10.17504/protocols.io.b4yyqxxw. OCT4, SSEA4 and AP Images from our hESC cultures were captured using a 10X objective on a fluorescence microscope (Zeiss ZEN 3.8). Magnification may differ depending on which microscope-camera set was used to capture the images. This was a result of which team within the collaboration was in charge of generating a specific cell line and its analysis through the QC steps.

### RNA Isolation and qRT-PCR

Total RNA was isolated from cell pellets with the RNeasy kit (Qiagen). 1-2µg of RNA was used for cDNA synthesis using High-capacity reverse transcriptase kit (ThermoFisher Scientific). Real-time qRT-PCR was performed on the QuantStudio 6 Flex thermocycler using PowerUp SYBR green master mix (ThermoFisher Scientific). All reactions were prepared according to the manufacturer instructions. Results were normalized to GAPDH and compared against human fibroblast samples (MRC-9, BJ1-hTERT and GM01660). All primer sequences used for qRT-PCR were listed in Supplemental Table 9. For a detailed protocol consult: https://doi.org/10.17504/protocols.io.4r3l22r9pl1y/v1. Plots were generated using GraphPad Prism (RRID:SCR_002798, version 10.1.2 [324]).

### Southern blot

Southern blotting was performed following standard protocols (https://cshprotocols.cshlp.org/content/2021/7/pdb.top100396#cited-by) to validate the structural integrity and exclude the loss of heterozygosity (LOH) at a genomic locus of interest (PRKN, DJ1/PARK7, FBXO7 and SYNJ1) resulting from CRISPR/Cas9 or prime editing-based genome editing experiments in hESCs. Southern blot probes were generated by PCR amplification (AccuPrime™ Taq DNA Polymerase, high fidelity (ThermoFisher)) of a 150bp to 600bp large genomic region 3’- and 5’ to the targeted genomic region. Southern blot probes were radiolabeled using the Prime-it Random Primer Labeling Kit (Agilent) according to the manufacturer’s instructions. Restriction digested genomic DNA isolated from clonally expanded genome edited hESC lines was separated on a 0.8% agarose (Sigma) gel, transferred to a nylon membrane (Amersham), and hybridized with 32P random primers labeled southern blot probes. Oligonucleotide sequences and restriction enzyme information can be found in Supplemental Table 9. Detailed protocols for southern blot analysis can be found on protocols.io (https://doi.org/10.17504/protocols.io.bp2l6xe6dlqe/v1).

---

**Supplemental Figure 1.**
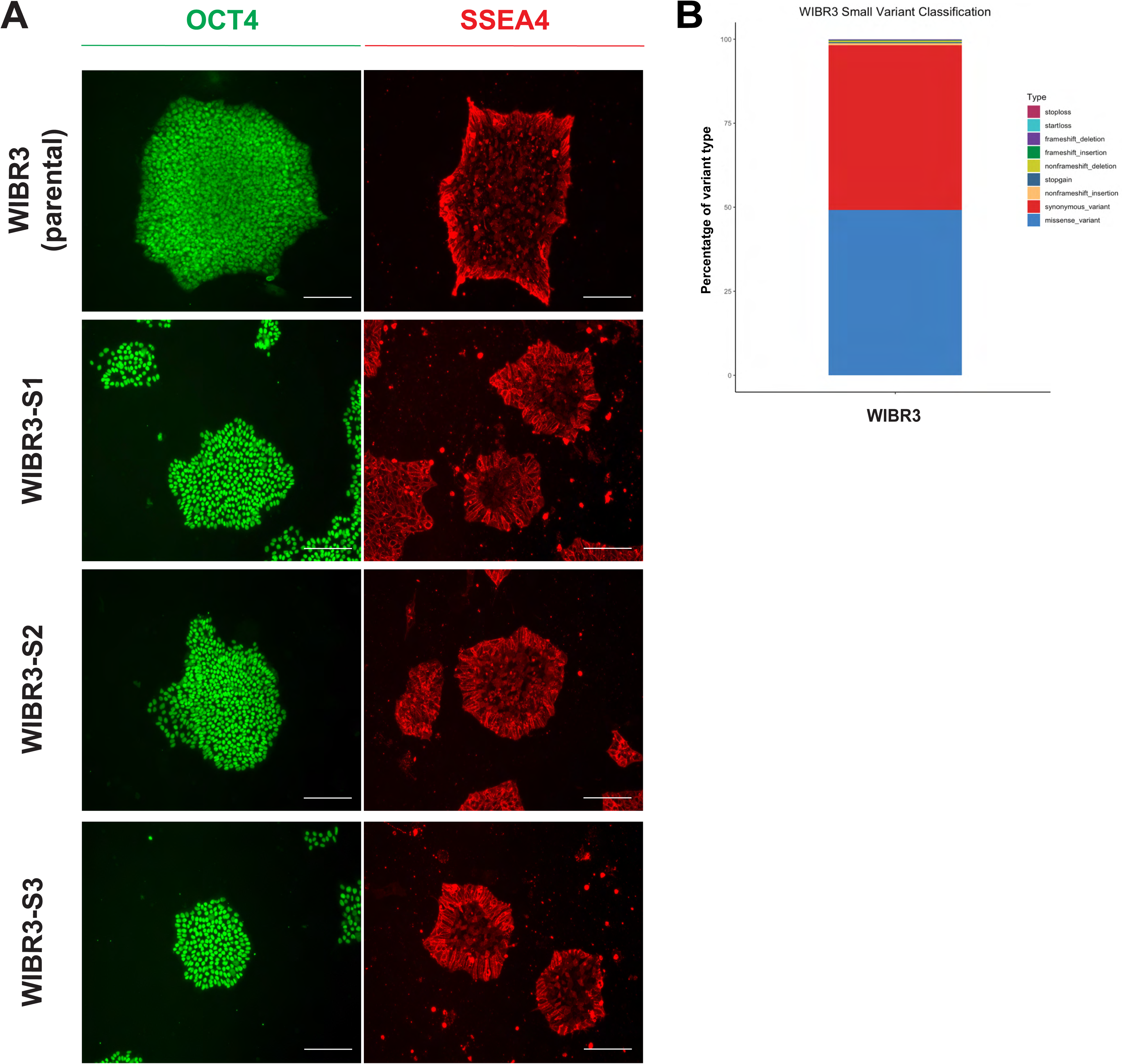
Characterization of WIBR3 in feeder free conditions. (A) Immunocytochemistry for pluripotency markers OCT4 (green) and SSEA4 (red) on hESC colonies in feeder free cultures from parental WIBR3 hESC and the clonal lines WIBR3-S1, WIBR3-S2 and WIBR3-S3. Scale bar 100 µm. (B) The percentage of genetic variant types present in WIBR3 grouped by their predicted consequences on coding sequences.

**Supplemental Figure 2.**
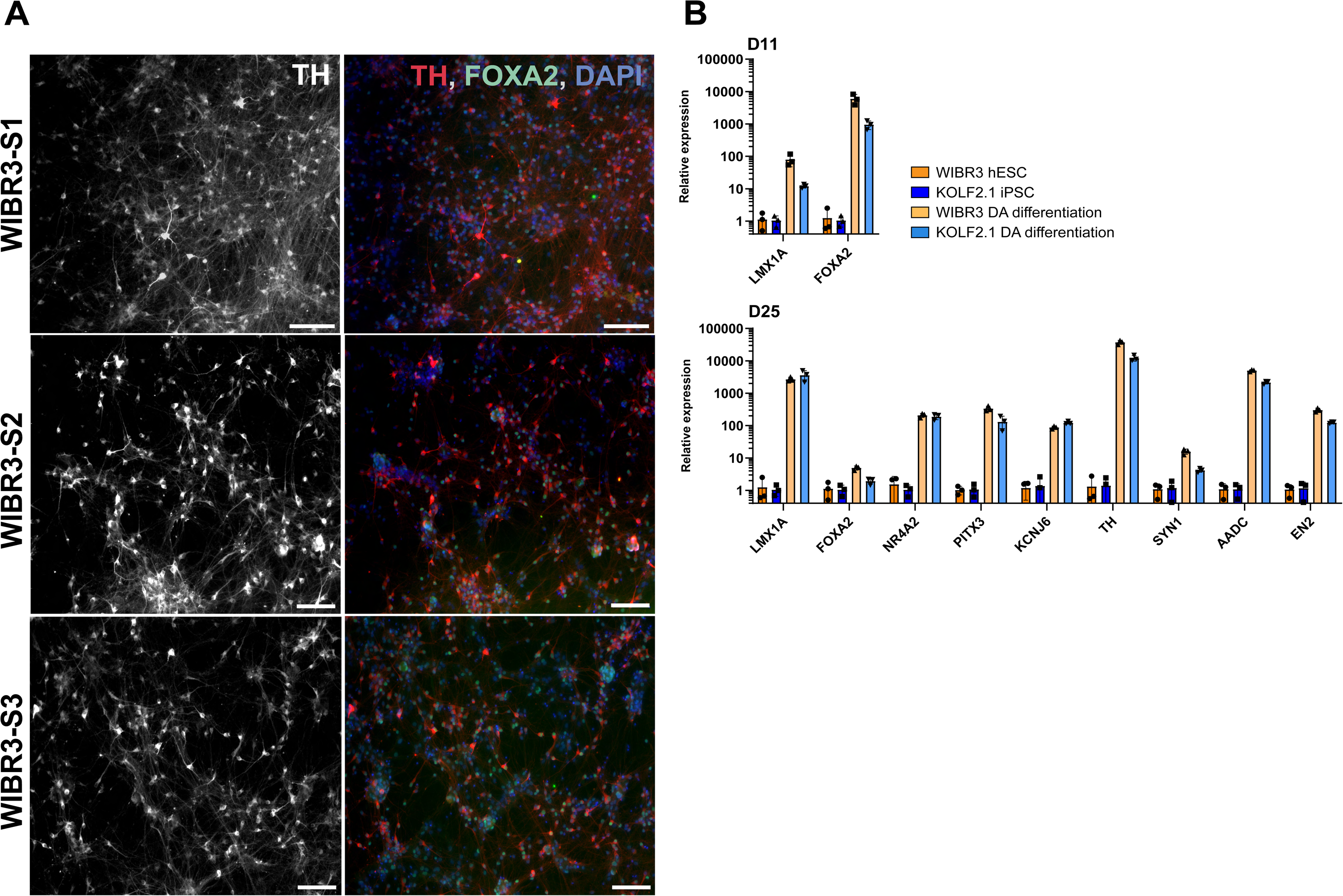
Dopaminergic neuron immunostaining and gene expression analysis. (A) Immunofluorescence of TH and FOXA2 in dopaminergic neurons (at day 35) derived from three subclones (S1, S2, S3) of WIBR3, scale bar 100 µm. (B) qRT-PCR quantification of midbrain floor plate progenitor and dopaminergic neuron markers at day 11 and day 25 of differentiation of WIBR3 and KOLF2.1 - derived cells. Relative gene expression is calculated relative to the expression of GAPDH and compared against their corresponding hPSC expression levels. (N = 3; MEAN ± SD).

**Supplemental Figure 3.**
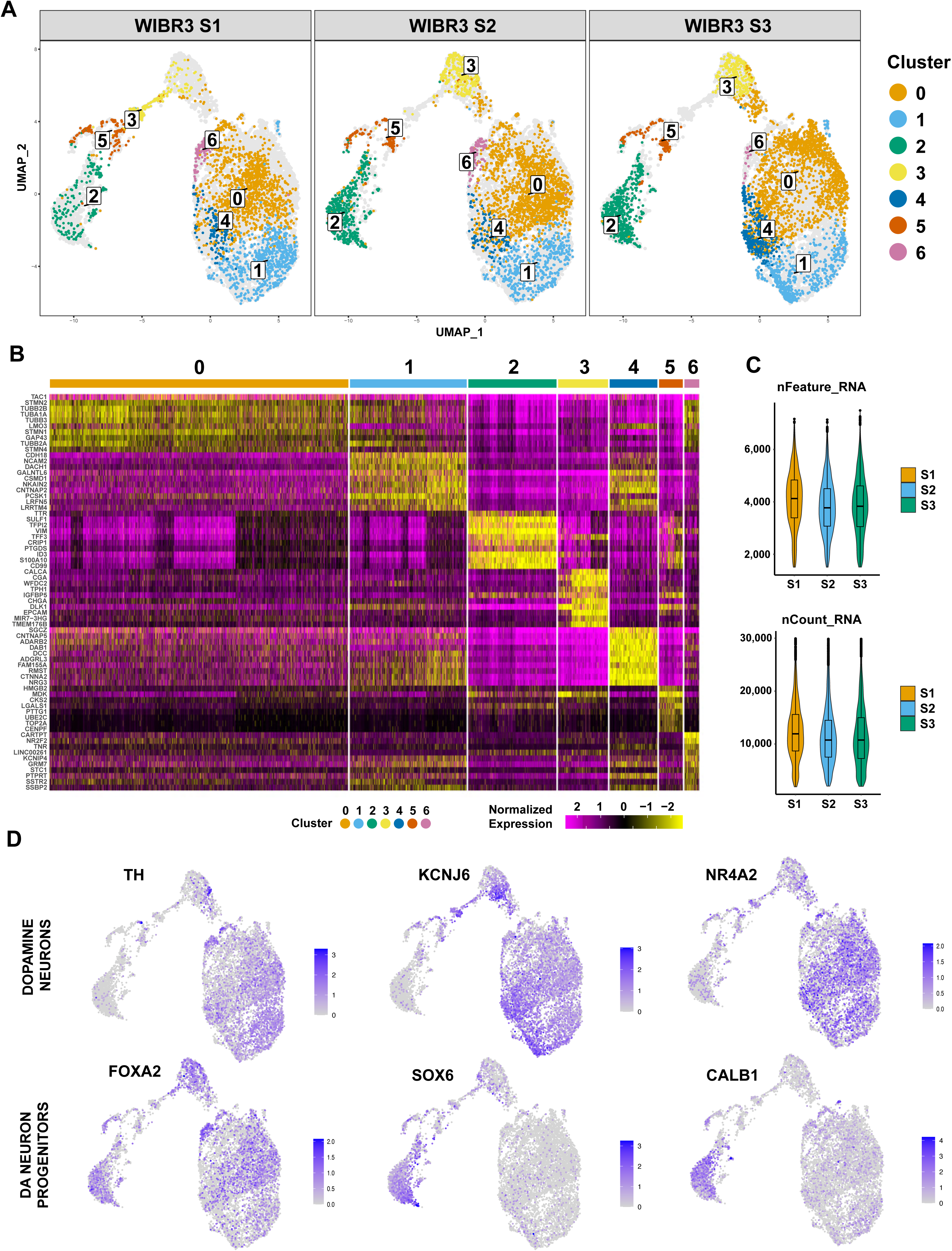
Quality control metrics of single cell data. (A) UMAPs of WIBR3 dopamine neurons of three subclones, to visualize distribution of cells in each cluster from an integrated seurat single cells dataset. (B) Heatmap showing top 10 genes differentially expressed in each cluster. (C) Quality control plots of nCount RNA reads (UMIs) and feature RNA reads (genes) for each subclone. (D) UMAP feature plots showing expression patterns across cell clusters of dopamine neuronal progenitor and mature dopamine neuron specific marker genes.

**Supplemental Figure 4.**
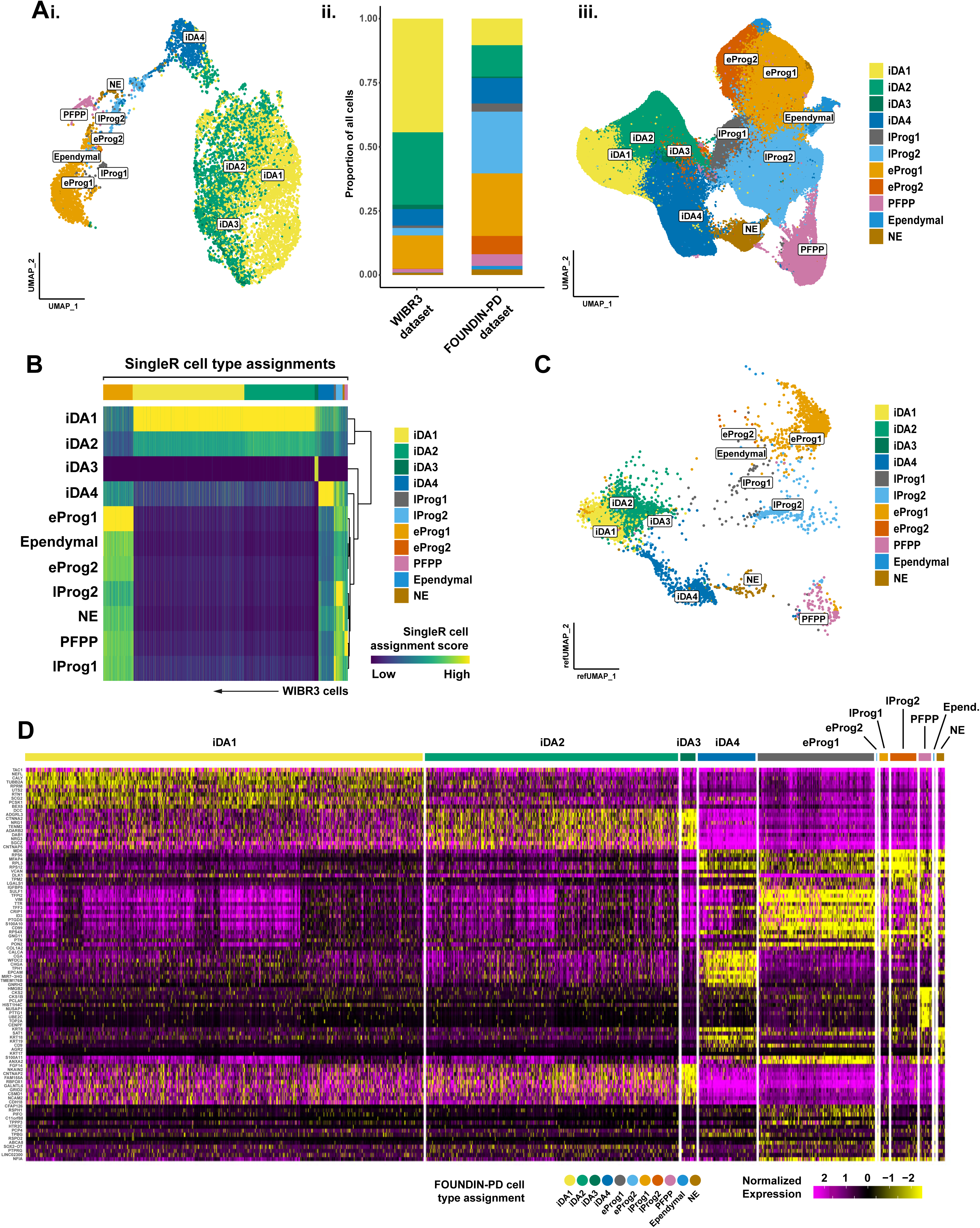
scRNASeq comparison of WIBR3 cells to the FOUNDIN-PD reference. (A) *(i)* UMAP plot of all 10,097 WIBR3 cells profiled, with labels representing a cell’s assignment to a corresponding cell type as defined in the FOUNDIN-PD reference dataset. Labels for each cell are identified using the SingleR classifier in R (Wilcoxon rank sum test). *(ii)* Stacked barplot depicting the proportion of cell type identities across the entire 10,097 cell WIBR3 dataset or the 416,216 cells in the FOUNDIN-PD reference dataset. *(iii)* UMAP of the 416,216 cells in the FOUNDIN-PD reference dataset, with the author’s default cell type labels applied. (B) Heatmap depicting the cell type classification of 10,097 WIBR3 cells (rows) into a list of cognate cell type labels as defined in the FOUNDIN-PD reference dataset. Plotted values represent normalized cell type assignment scores as calculated by the SingleR package in R (Wilcoxon rank-sum test). Any cells retaining undefined identities after SingleR classification (“NA”) are omitted from downstream analyses. (C) All 10,097 WIBR3 cells depicted in the UMAP space of the FOUNDIN-PD dataset shown in panel *A iii*, with each cell depicted with a FOUNDIN-PD cell type label applied by SingleR. (D) Heatmap of WIBR3 cells (columns) grouped by FOUNDIN-PD labels as applied by SingleR. Rows depict the top 10 marker genes that define the WIBR3 cells assigned to each FOUNDIN-PD cell type identity. Markers lists are identified with the “FindAllMarkers” function in Seurat, using the “MAST” test.

**Supplemental Figure 5.**
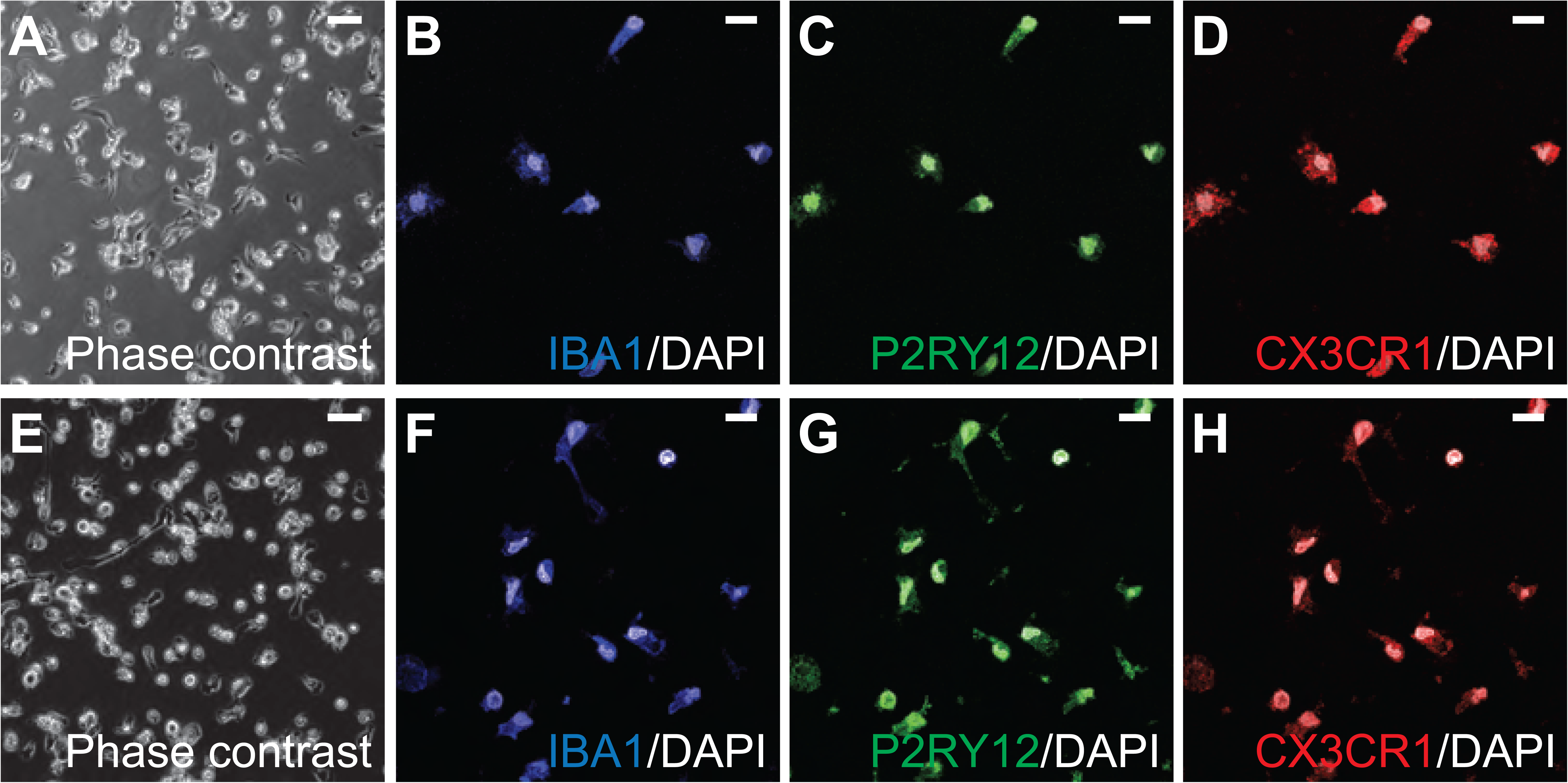
I*n vitro* differentiation of microglia from WIBR3-S2 and WIBR3-S3. (A-D) Representative phase contrast (A) and immunostaining (B-C) images of *in vitro* differentiated microglia derived from subclone WIBR-S2 for microglia-specific markers IBA1, P2RY12, and CX3CR1 (terminal diff day 14). (E-H) Representative phase contrast (E) and immunostaining (F-H) images of *in vitro* differentiated microglia derived from subclone WIBR-S3 for microglia-specific markers IBA1, P2RY12, and CX3CR1 (terminal diff day 14). Scale bar (phase contrast): 50 µm; Scale bar (ICC): 10 µm.

**Supplemental Figure 6.**
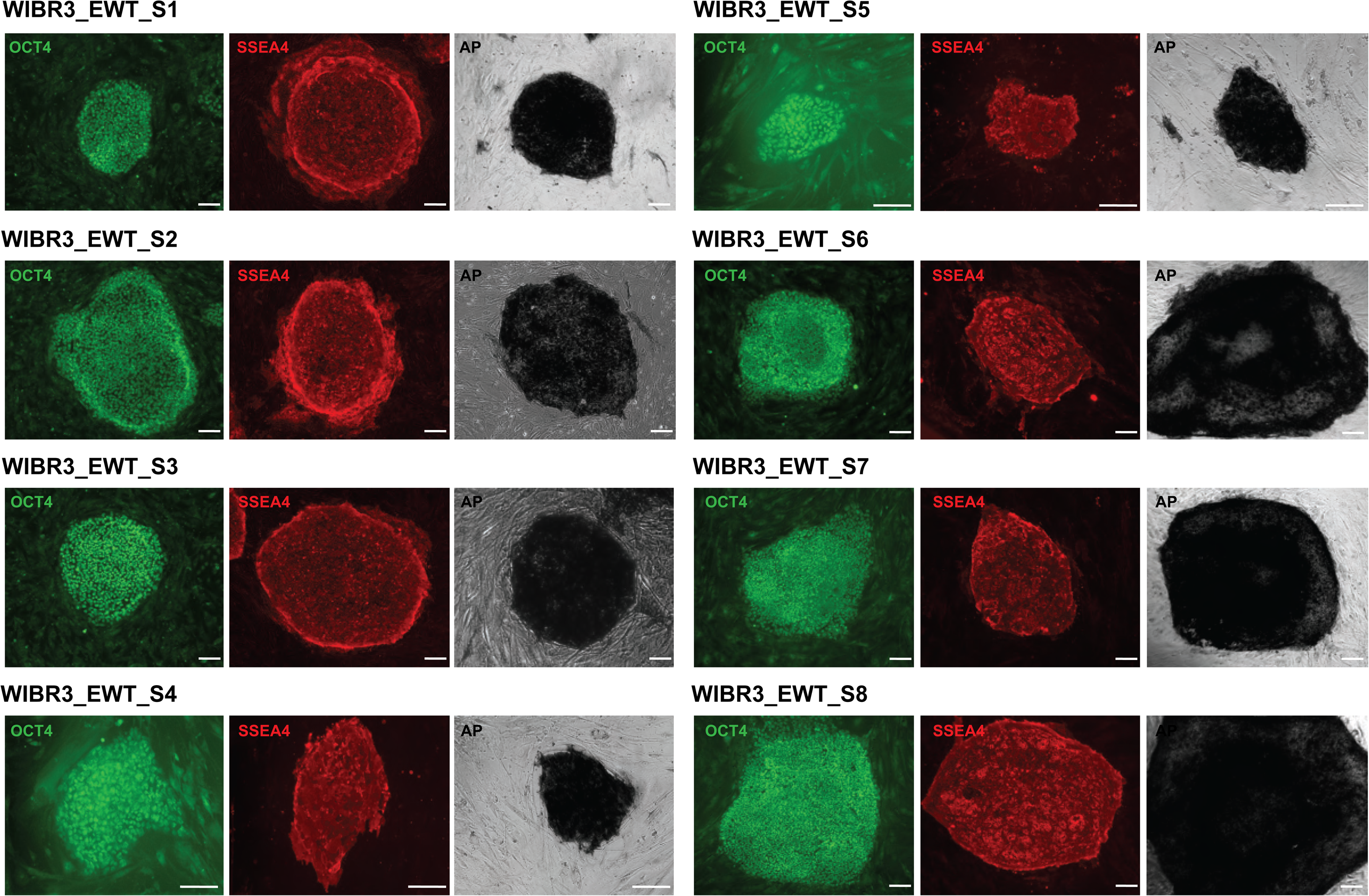
Characterization of edited wild type cell lines. Immunocytochemistry of edited wild-type (EWT) cells lines for pluripotency markers OCT4 (green), SSEA4 (red) and alkaline phosphatase (black). EWT refers to cell lines that have undergone the editing pipelines but were not genetically modified and remain genotypically wild type. WIBR3_EWT_S1-3 were isolated from a prime editing experiment using Pipeline B, WIBR3_EWT_S4-5 were isolated from CRISPR/Cas9-facilitated HDR experiments using Pipeline B and WIBR3_EWT_S6-8 were isolated from a prime editing experiment using Pipeline A. Scale bar 100 µm.

**Supplemental Figure 7.**
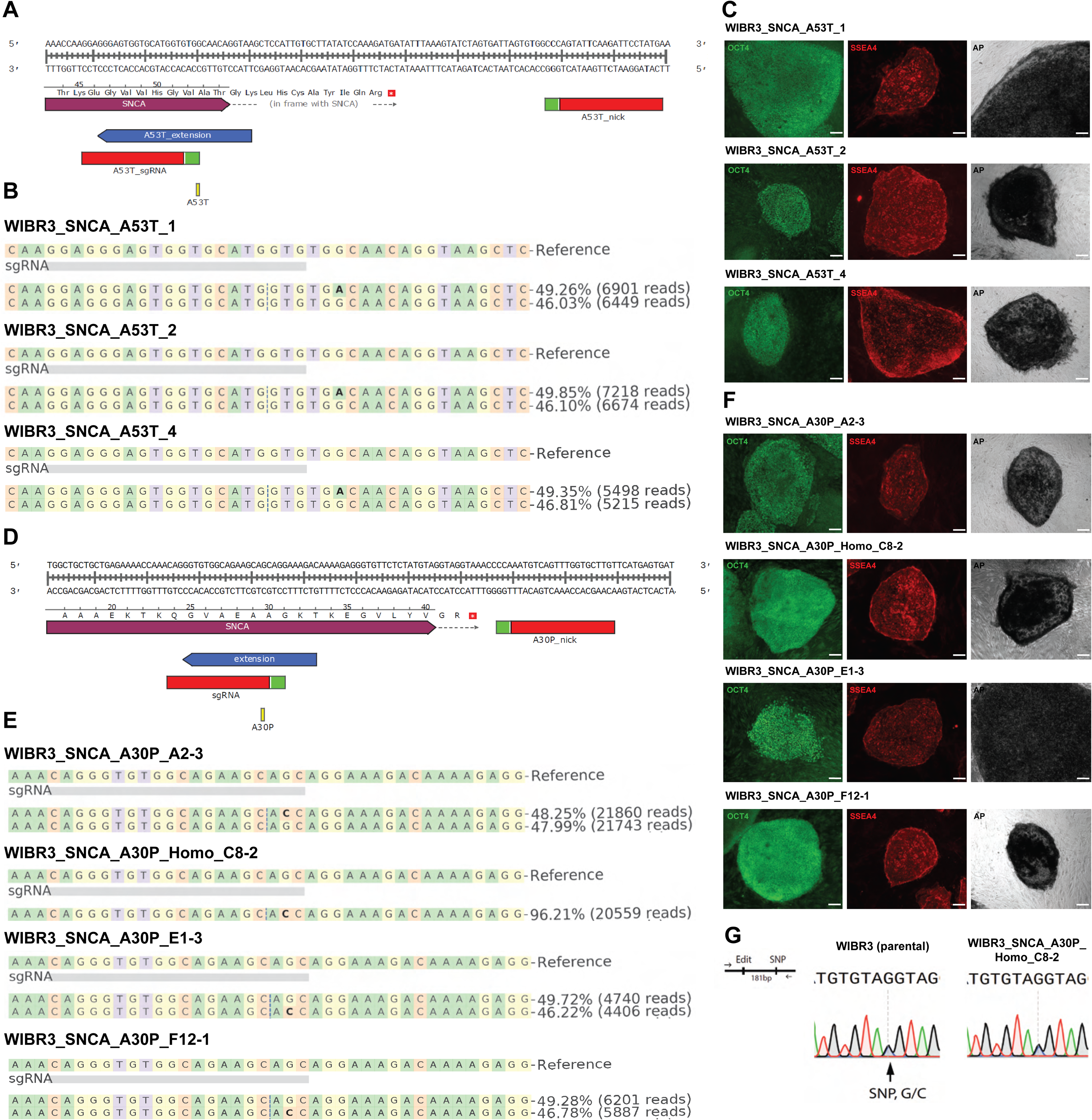
Genome editing and quality control of WIBR3 hESCs carrying PD-associated mutations in *SNCA*. (A) Targeting strategy to generate SNCA A53T mutation using prime editing. (B) NGS-based genotyping to confirm correct editing for SNCA A53T in clones WIBR3_SNCA_A53T_1, WIBR3_SNCA_A53T_2 and WIBR3_SNCA_A53T_4. Bold bases indicate base substitutions. (C) Immunocytochemistry of hESC cultures for pluripotency markers OCT4 (green), SSEA4 (red), and alkaline phosphatase (black). Scale bar 100 µm. (D) Targeting strategy to generate SNCA A30P mutation by prime editing. (E) NGS-based genotyping to confirm correct editing for SNCA A30P in clones WIBR3_SNCA_A30P_A2-3, WIBR3_SNCA_A30P_Homo_C8-2, WIBR3_SNCA_A30P_E1-3 and WIBR3_SNCA_A30P_F12-1. Bold bases indicate base substitution. The generation of the WIBR3_SNCA_A30P cell lines was previously reported^70^ (F) Immunocytochemistry of hESC cultures for pluripotency markers OCT4 (green), SSEA4 (red) and alkaline phosphatase (black). Scale bar 100 µm. (G) Zygosity analysis using Sanger sequencing to detect heterozygous neighboring SNV to exclude LOH in clone WIBR3_SNCA_A30P_Homo_C8-2.

**Supplemental Figure 8.**
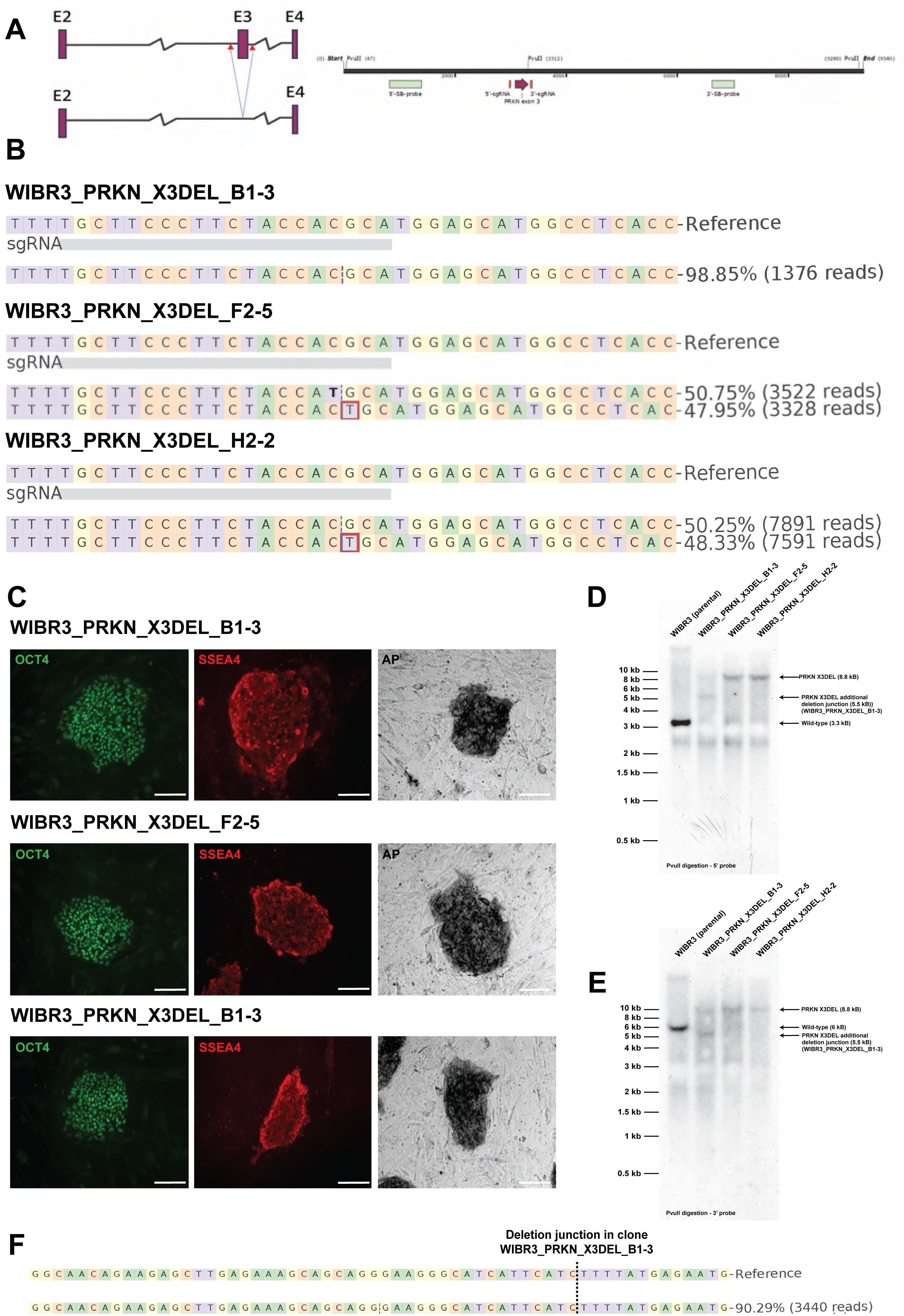
Genome editing and quality control of WIBR3 hESCs carrying PD-associated deletions in *PRKN*. (A) Schematic illustrating targeting strategy to generate PRKN X3del mutation by CRISPR/Cas9 dual guide strategy. Included are genomic location of sgRNAs, Southern blot (SB) probes and restriction enzymes used for southern blot. (B) NGS-based genotyping to confirm correct editing for PRKN X3del in clones WIBR3_PRKN_X3DEL_B1-3, WIBR3_PRKN_X3DEL_F2-5 and WIBR3_PRKN_X3DEL_H2-2 (deletion junction). Red box indicates single base insertion. Bold bases indicate base substitution. (C) Immunocytochemistry of hESC cultures for pluripotency markers OCT4 (green), SSEA4 (red) and alkaline phosphatase (black). Scale bar 100 µm. (D, E) Southern blot analysis of WIBR3_PRKN_X3DEL cell lines to exclude LOH. Genomic DNA was digested with indicated enzymes and hybridized with 3’ and 5’ probes indicated in (A). Expected fragment size for wild type and PRKN_X3DEL allele are indicated for each digest. This analysis indicates that the clone WIBR3_PRKN_X3DEL_B1-3 carries a larger deletion on one allele. (F) NGS analysis of the new deletion junction in the clone WIBR3_PRKN_X3DEL_B1-3.

**Supplemental Figure 9.**
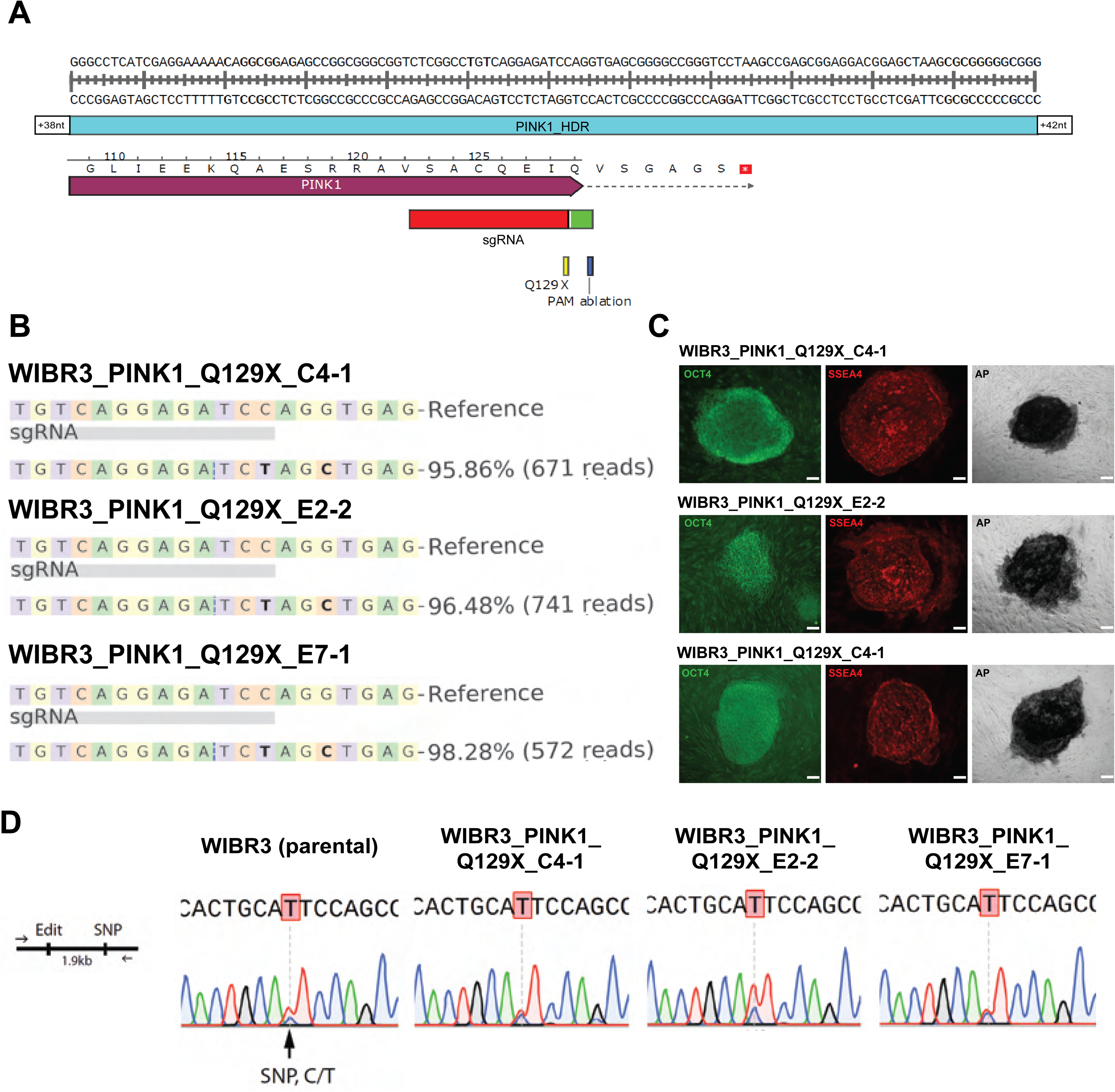
Genome editing and quality control of WIBR3 hESCs carrying PD-associated mutations in *PINK1*. (A) Schematic illustrating targeting strategy to generate PINK1 Q129X mutation by CRISPR/Cas9 facilitated HDR. (B) NGS-based genotyping to confirm correct editing for PINK1 Q129X in clones WIBR3_PINK1_Q129X_C4-1, WIBR3_PINK1_Q129X_E2-2 and WIBR3_PINK1_Q129X_E7-1. Bold bases indicate base substitution. (C) Immunocytochemistry of hESC cultures for pluripotency markers OCT4 (green), SSEA4 (red) and alkaline phosphatase (black). Scale bar 100 µm. (D) Zygosity analysis using Sanger sequencing to detect heterozygous SNV to exclude LOH in any of the PINK1 Q129X clones.

**Supplemental Figure 10.**
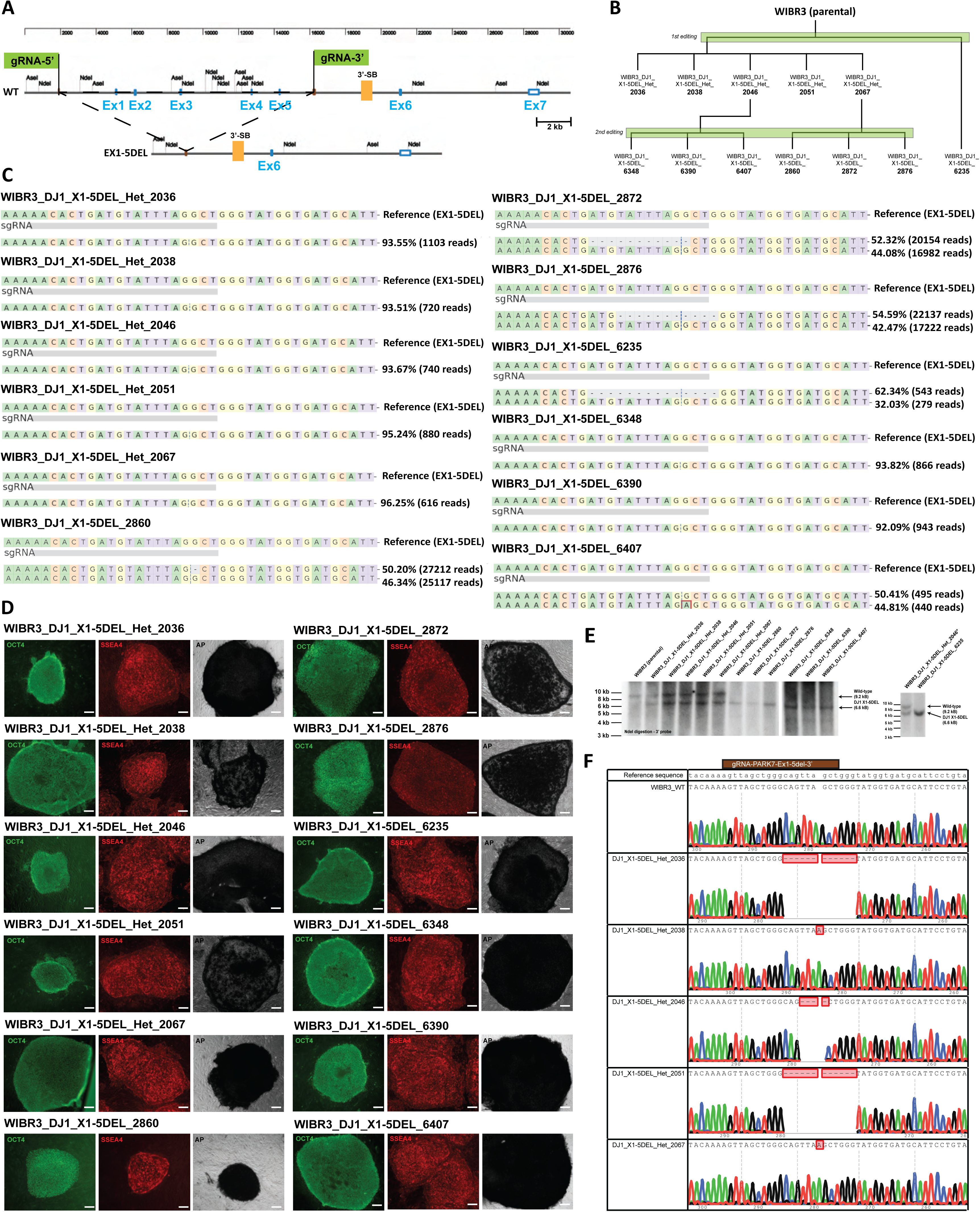
Genome editing and quality control of WIBR3 hESCs carrying PD-associated deletions in DJ1/*PARK7*. (A) Schematic illustrating targeting strategy to generate DJ1/PARK7 Ex1-5del mutation by CRISPR/Cas9 dual guide strategy. Included are genomic location of sgRNAs, Southern blot (SB) probe and restriction enzyme used for southern blot. (B) Schematic representation of the parental lineage of heterozygous and homozygous clones carrying Ex1-5del mutation in the iSCORE-PD collection. (C) NGS-based genotyping to confirm correct editing for DJ1/PARK7 Ex1-5del in heterozygous clones WIBR3_DJ1_X1-5DEL_Het_2036/2038/2046/2051/2067 and homozygous clones WIBR3_DJ1_X1-5DEL_2860/2872/2876/6235/6348/6390/6407 (deletion junction). “-” indicates base deletion. Red box indicates single base insertion. (D) Immunocytochemistry of hESC cultures for pluripotency markers OCT4 (green), SSEA4 (red), and alkaline phosphatase (black). Scale bar 100 µm. (E) Southern blot analysis of homozygous and heterozygous WIBR3_DJ1_X1-5DEL cell lines to exclude LOH. Genomic DNA was digested with indicated enzymes and hybridized with 3’ probes indicated in (A). Expected fragment size for wild type and DJ1_X1-5DEL alleles are indicated. (F) Sanger sequencing results for secondary WT allele alteration in heterozygous cell lines after 1^st^ editing experiment.

**Supplemental Figure 11.**
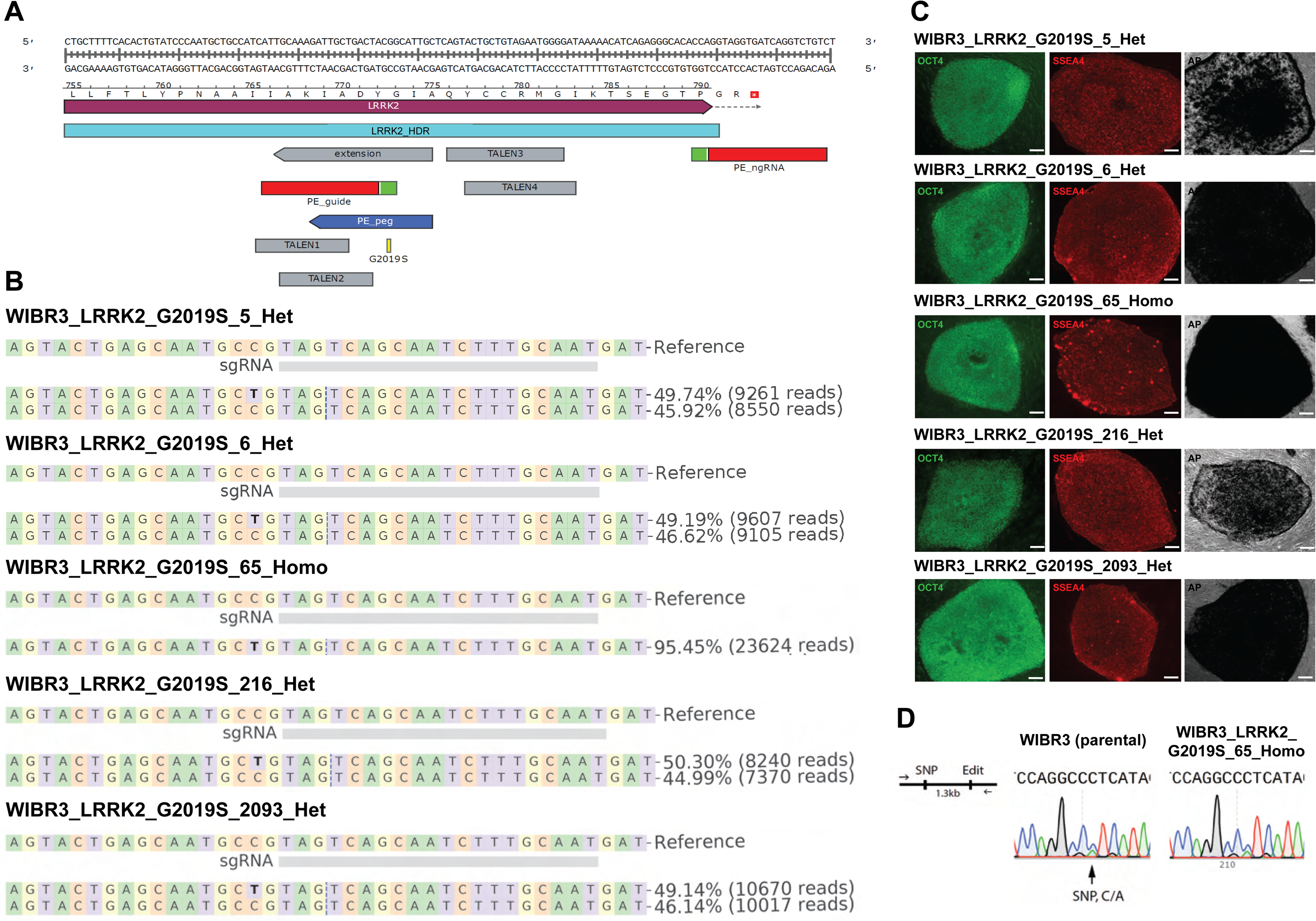
Genome editing and quality control of WIBR3 hESCs carrying PD-associated mutations in *LRRK2*. (A) Schematic illustrating targeting strategy to generate LRRK2 G2019S mutation by prime editing and TALEN or CRISPR/Cas9 facilitated HDR. (B) NGS-based genotyping to confirm correct editing for LRRK2 G2019S in clones WIBR3_LRRK2_G2019S_5_Het, WIBR3_LRRK2_G2019S_6_Het, WIBR3_LRRK2_G2019S_65_Homo, WIBR3_LRRK2_G2019S_216_Het and WIBR3_LRRK2_G2019S_2093_Het. Bold bases indicate base substitution. The generation of the WIBR3_LRRK2_G2019S_5_Het and WIBR3_LRRK2_G2019S_6_Het cell lines was already reported^70^. (C) Immunocytochemistry of hESC cultures for pluripotency markers OCT4 (green), SSEA4 (red) and alkaline phosphatase (black). Scale bar 100 µm. (D) Zygosity analysis using Sanger sequencing to detect heterozygous SNV to exclude LOH in cell line WIBR3_LRRK2_G2019S_65_Homo.

**Supplemental Figure 12.**
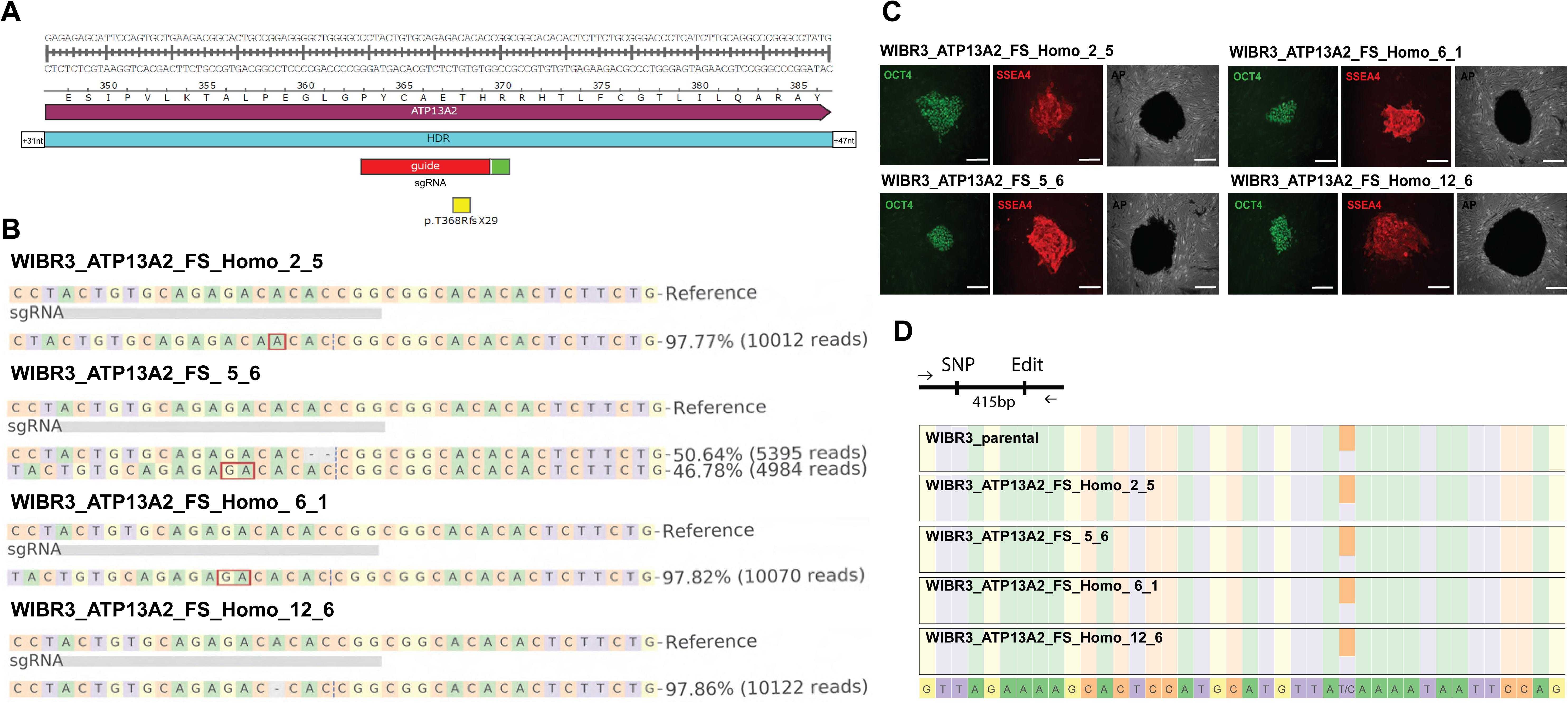
Genome editing and quality control of WIBR3 hESCs carrying PD-associated alterations in *ATP13A2*. (A) Schematic illustrating targeting strategy to generate CRISPR/Cas9 mediated frameshift mutation in ATP13A2. (B) NGS-based genotyping to confirm a frameshift mutation in ATP13A2 in clones WIBR3_ATP13A2_FS_Homo_2_5, WIBR3_ATP13A2_FS_5_6, WIBR3_ATP13A2_FS_Homo_6_1, and WIBR3_ATP13A2_FS_Homo_12_6. Red box indicates base insertion. “-” indicates base deletion. (C) Immunocytochemistry of hESC cultures for pluripotency markers OCT4 (green), SSEA4 (red) and alkaline phosphatase (black). Scale bar 100 µm. (D) Zygosity analysis using NGS sequencing to detect heterozygous SNV to exclude LOH in all ATP13A2 clones.

**Supplemental Figure 13.**
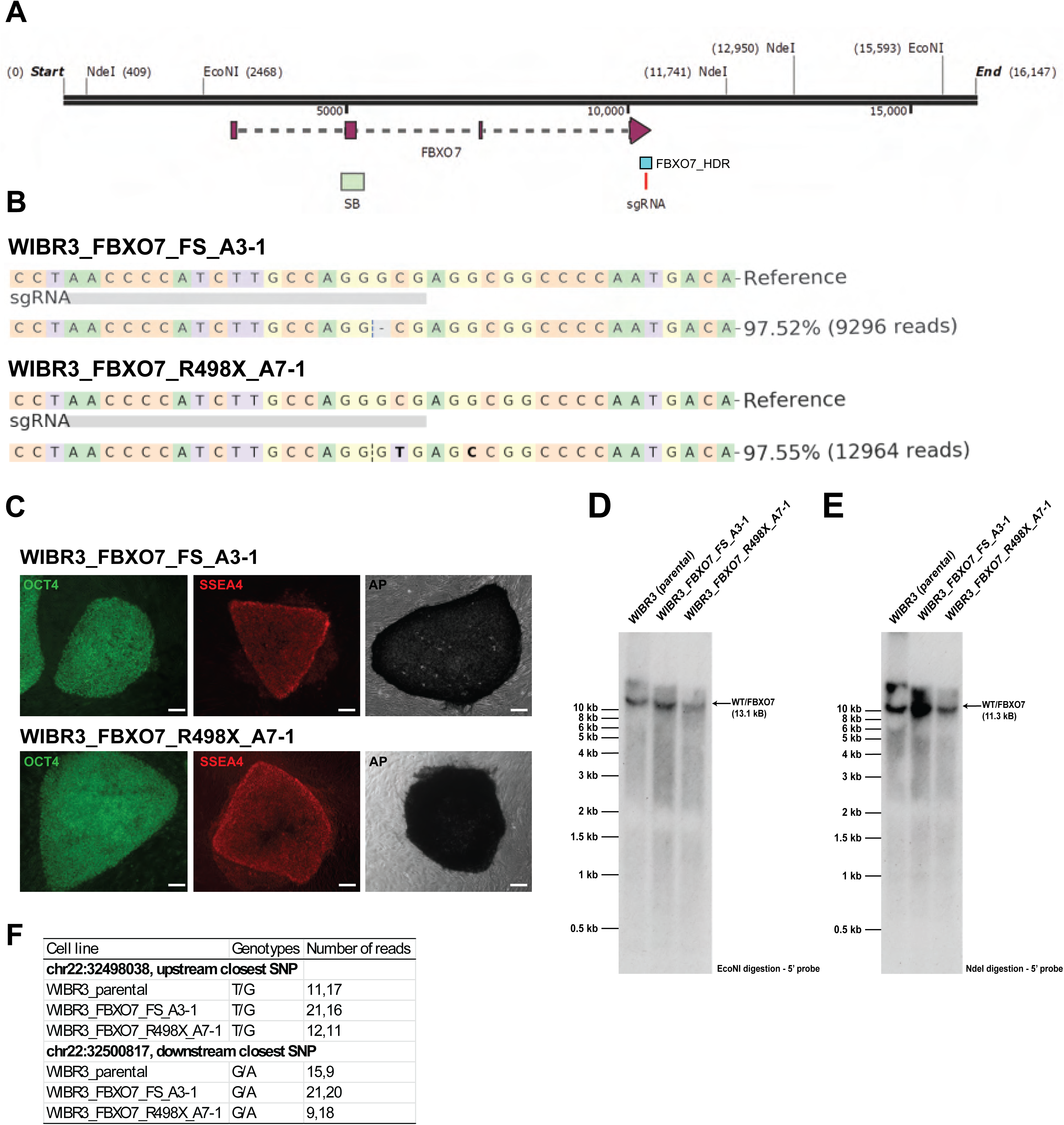
Genome editing and quality control of WIBR3 hESCs carrying PD-associated alterations in *FBXO7*. (A) Schematic illustrating targeting strategy to generate CRISPR/Cas9-mediated R498X/frameshift mutation in FBXO7. Included are genomic location of sgRNA, Southern blot (SB) probe and restriction enzymes used for southern blot. (B) NGS-based genotyping to confirm a frameshift in mutation in FBXO7 in clones WIBR3_FBXO7_FS_A3-1 and WIBR3_FBXO7_R498X_A7-1. “-” indicates base deletion. Bold bases indicate base substitution. (C) Immunocytochemistry of hESC cultures for pluripotency markers OCT4 (green), SSEA4 (red) and alkaline phosphatase (black). Scale bar 100 µm. (D, E) Southern blot analysis to exclude LOH. Genomic DNA was digested with indicated enzymes and hybridized with a 5’ probe indicated in (A). Expected fragment size for wild type and FBXO7 frameshift alleles are indicated for each digest. (F) WGS data was used to exclude LOH of FBXO7 clones by assessing closest upstream and downstream SNVs.

**Supplemental Figure 14.**
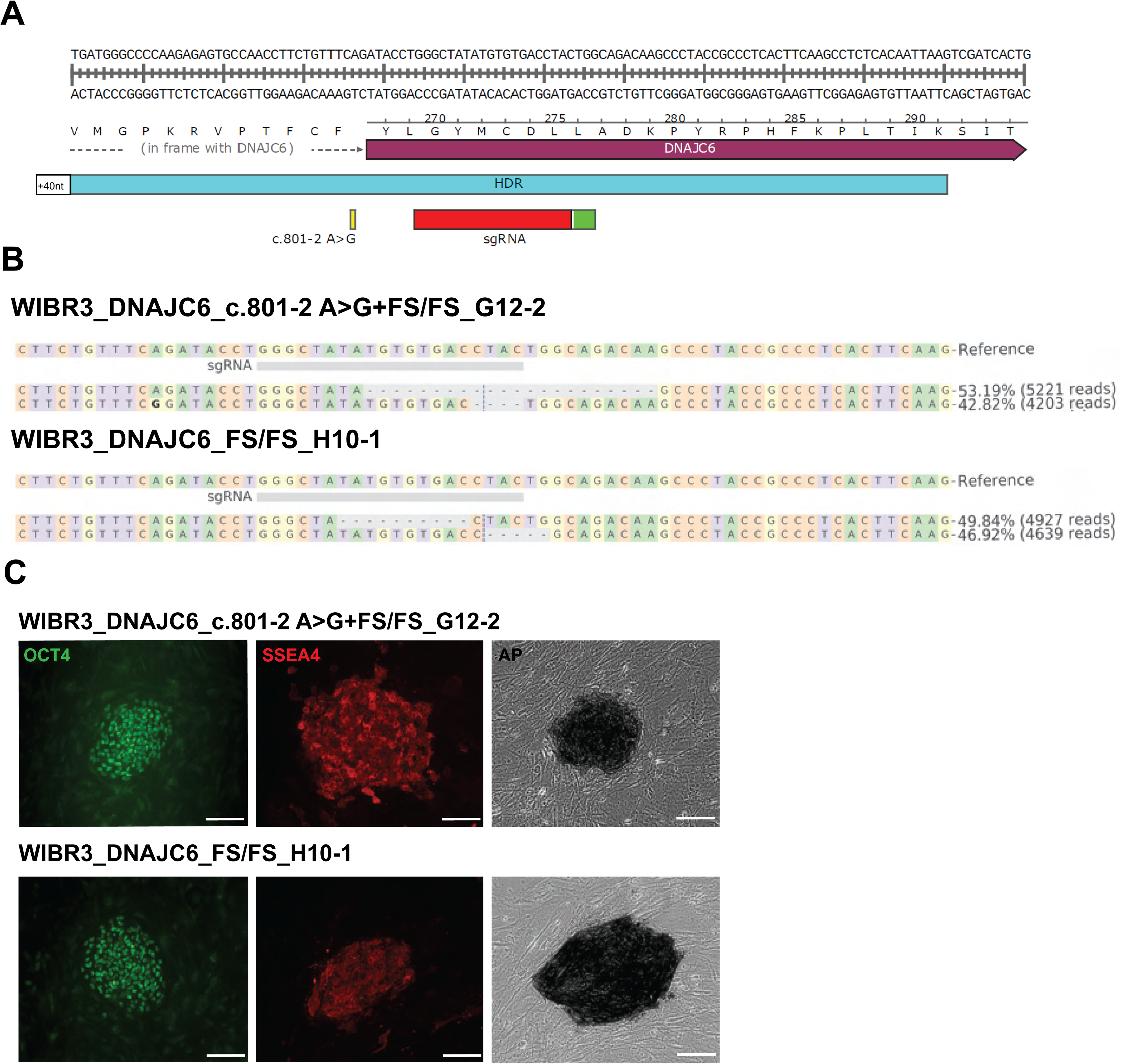
Genome editing and quality control of WIBR3 hESCs carrying PD-associated alterations *DNAJC6*. A) Schematic illustrating targeting strategy to generate DNAJC6 c.801-2 A>G/FS mutation by CRISPR/Cas9 facilitated HDR. (B) NGS-based genotyping to confirm correct editing for DNAJC6 c.801-2 A>G and frameshift mutation in clones WIBR3_DNAJC6_c.801-2 A>G+FS/FS_G12-2 and WIBR3_DNAJC6_FS/FS_H10-1. “-” indicates base deletion. (C) Immunocytochemistry of hESC cultures for pluripotency markers OCT4 (green), SSEA4 (red) and alkaline phosphatase (black). Scale bar 100 µm.

**Supplemental Figure 15.**
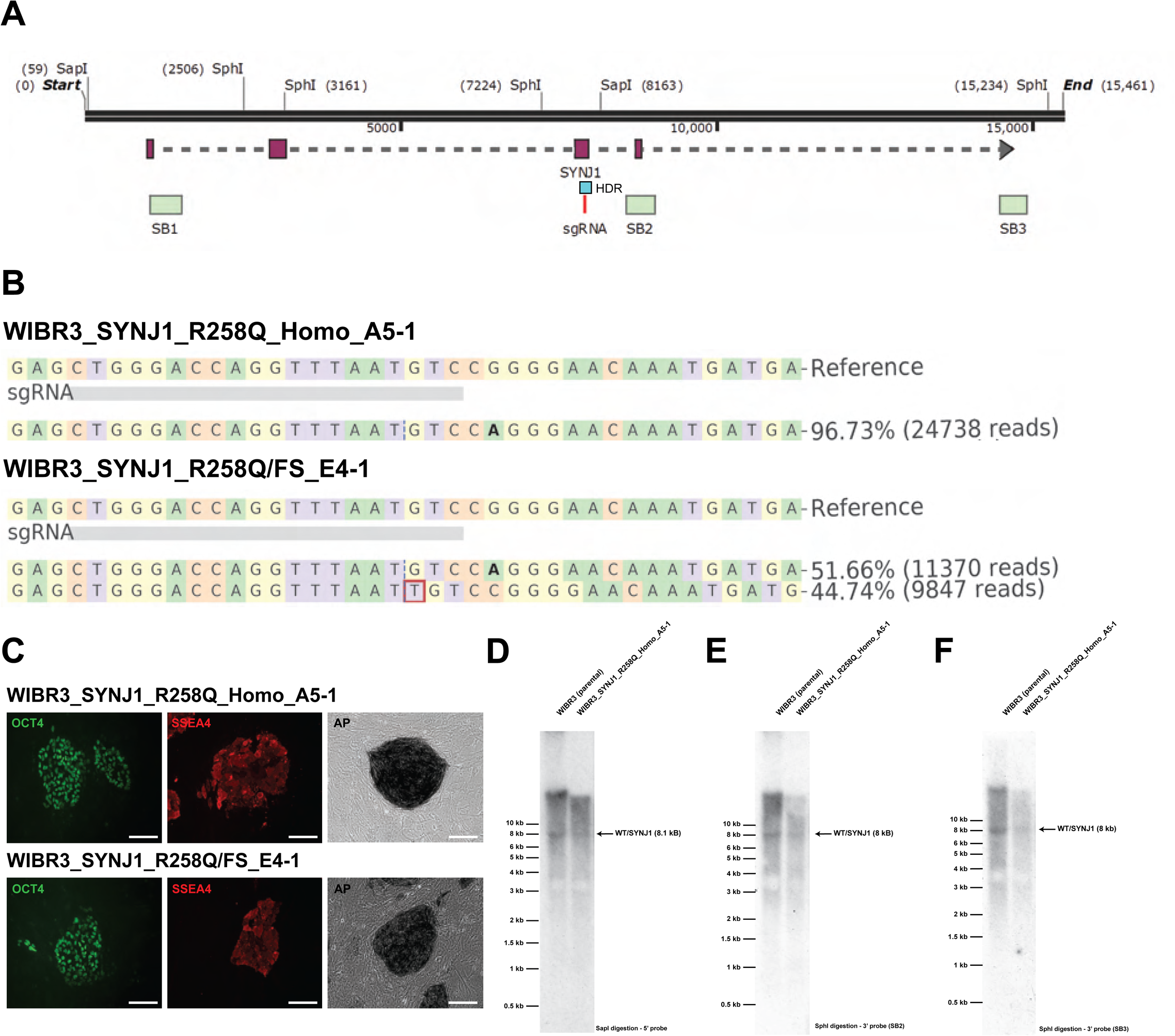
Genome editing and quality control of WIBR3 hESCs carrying PD-associated alterations *SYNJ1*. (A) Schematic illustrating targeting strategy to generate SYNJ1 R258Q/FS mutation by CRISPR/Cas9 facilitated HDR. (B) NGS-based genotyping to confirm correct editing for SYNJ1 R258Q and frameshift mutation in clones WIBR3_SYNJ1_R258Q_Homo_A5-1 and WIBR3_SYNJ1_R258Q/FS_E4-1. Bold bases indicate base substitution. Red box indicates base insertion. (C) Immunocytochemistry of hESC cultures for pluripotency markers OCT4 (green), SSEA4 (red) and alkaline phosphatase (black). Scale bar 100 µm. (D-F) Southern blot analysis of WIBR3_SYNJ1 cell lines to excluded LOH. Genomic DNA was digested with indicated enzymes and hybridized with probes indicated in (A). Expected fragment size for wild-type and SYNJ1 alleles are indicated for each digest.

**Supplemental Figure 16.**
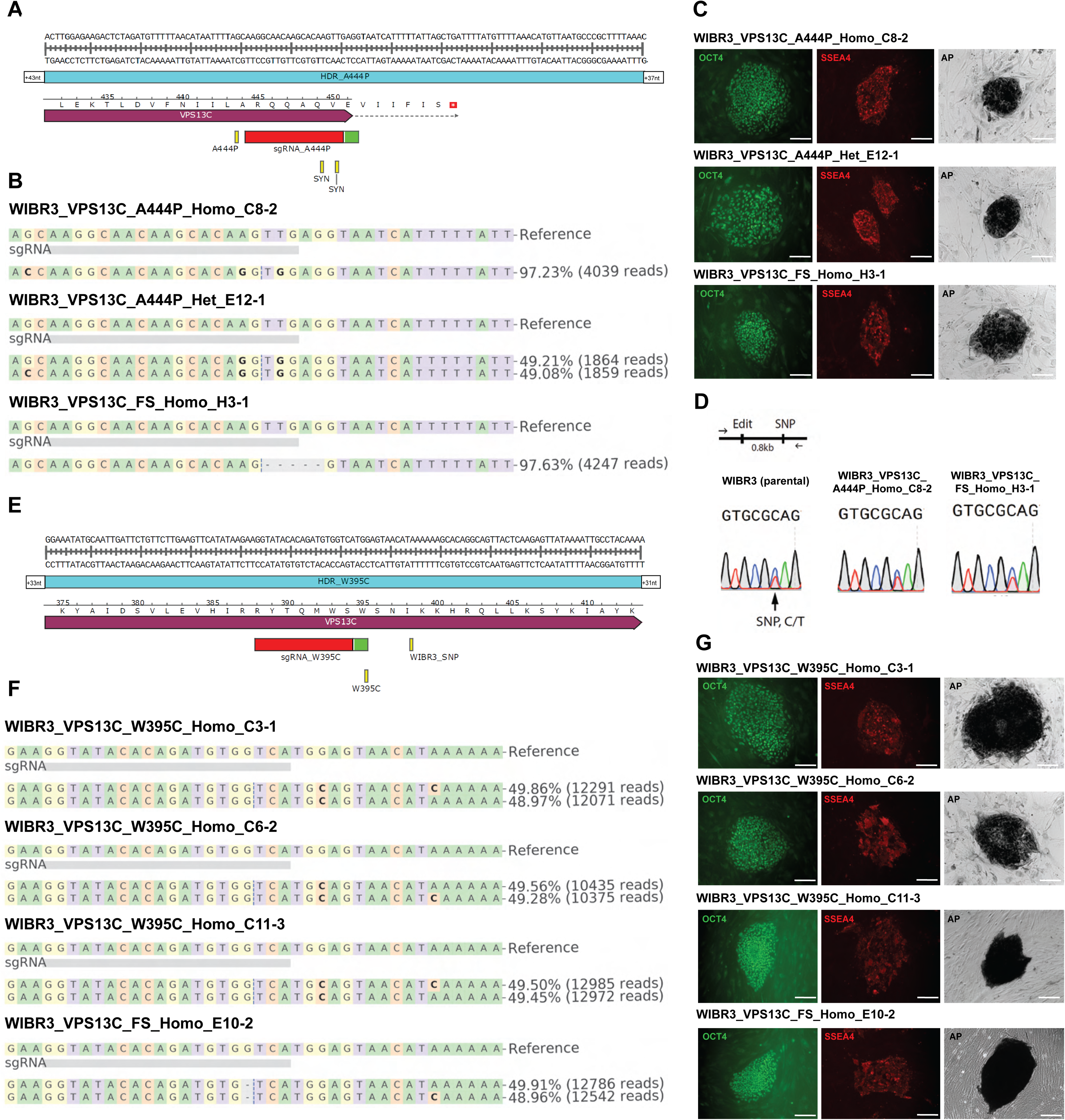
Genome editing and quality control of WIBR3 hESCs carrying PD-associated alterations *VPS13C*. (A) Schematic illustrating targeting strategy to generate VPS13C A444P mutation by CRISPR/Cas9 facilitated HDR. Included are genomic location of the A444P mutation, sgRNA, ssODN template including synonymous ssODN mutations (SYN) in sgRNA-target site to prevent re-cutting of edited alleles. (B) NGS-based genotyping to confirm correct editing for A444P and frameshift mutation in VPS13C in the clones WIBR3_VPS13C_A444P_Homo_C8-2, WIBR3_VPS13C_A444P_Het_E12-1 and WIBR3_VPS13C_FS_Homo_H3-1. Bold bases indicate base substitution. “-” indicates base deletion. (C) Immunocytochemistry of hESC cultures for pluripotency markers OCT4 (green), SSEA4 (red) and alkaline phosphatase (black). Scale bar 100 µm. (D) Zygosity analysis using Sanger sequencing to detect heterozygous SNV to exclude LOH in clones WIBR3_VPS13C_A444P_Homo_C8-2 and WIBR3_VPS13C_A444P_Homo_H3-1. (E) Schematic illustrating targeting strategy to generate VPS13C W395C and frameshift mutation by CRISPR/Cas9 facilitated HDR. (F) NGS-based genotyping to confirm correct editing for VPS13C W395C and frameshift mutation in the clones WIBR3_VPS13C_W395C_Homo_C3-1, WIBR3_VPS13C_W395C_Homo_C6-2, WIBR3_VPS13C_W395C_Homo_C11-3 and WIBR3_VPS13C_FS_Homo_E10-2. Bold bases indicate base substitution. “-” indicates base deletion. (G) Immunocytochemistry of hESC cultures for pluripotency markers OCT4 (green), SSEA4 (red) and alkaline phosphatase (black). Scale bar 100 µm.

**Supplemental Figure 17.**
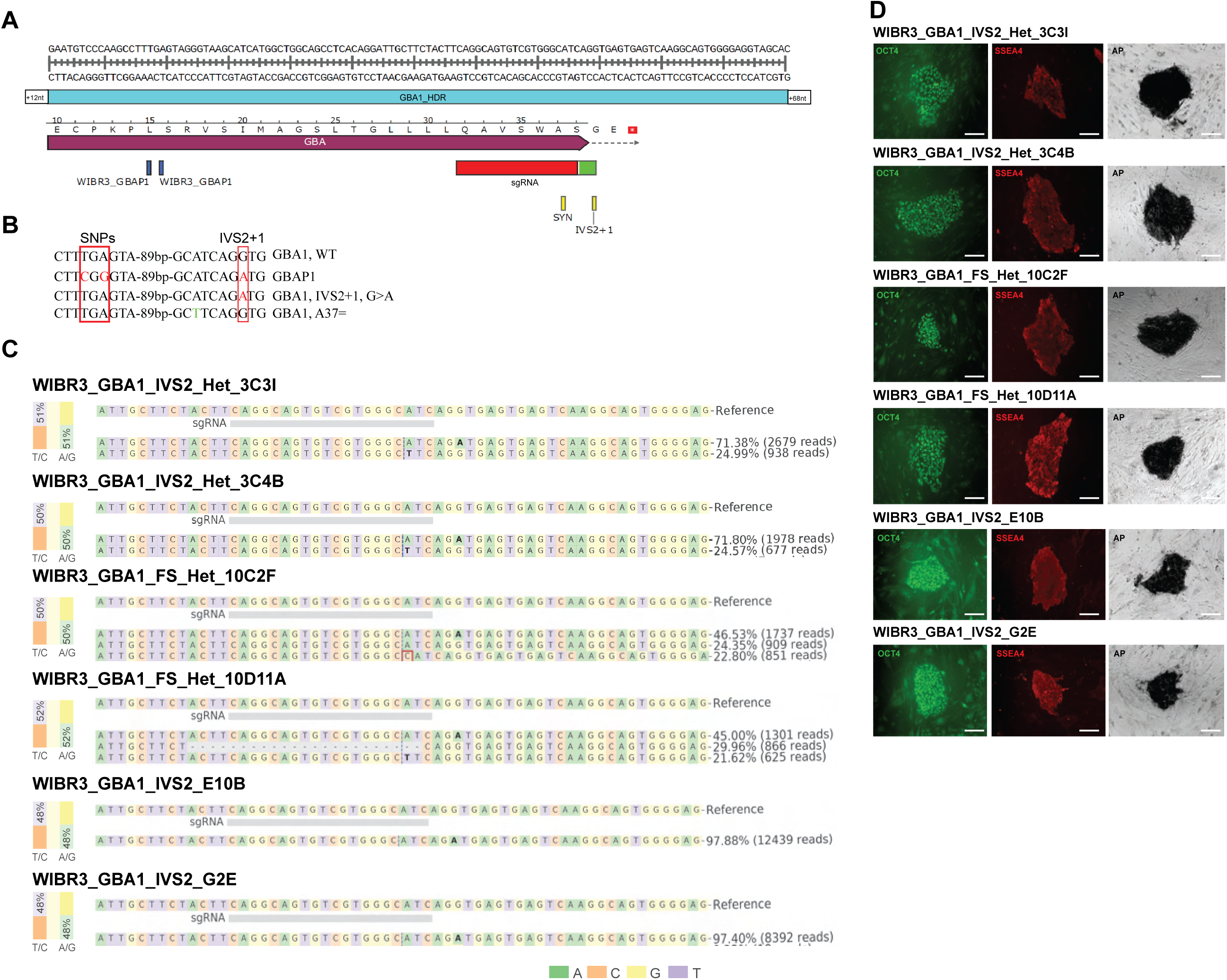
Genome editing and quality control of WIBR3 hESCs carrying PD-associated alterations *GBA1*. (A) Schematic illustrating targeting strategy to generate GBA1 IVS2+1 mutation by CRISPR/Cas9 facilitated HDR. To generate the heterozygous mutations a competing HDR template (ssODNs) containing synonymous mutations (SYN) in the sgRNA-target site was used. (B) Sequences alignment of the wild type GBA1 gene (GBA1, WT), its pseudogene (GBAP1), the engineered GBA1 IVS2+1 mutation (GBA1, IVS2+1 G>A) and synonymous mutation (GBA1, A37=) in the targeted region. The mutated nucleotides and SNVs used to calculate the allelic balance between GBA1 and GBAP1 are highlighted in red boxes. (C) NGS-based genotyping to confirm correct editing of the GBA IVS2+1 mutation and evaluate proper zygosity balance with GBAP1 for Clones WIBR3_GBA_IVS2_Het_3C3I, WIBR3_GBA_IVS2_Het_3C4B, WIBR3_GBA_FS_Het_10C2F, WIBR3_GBA_IVS2_E10B and WIBR3_GBA_IVS2_G2E. Bold bases indicate base substitution. Red box indicates base insertion. (D) Immunocytochemistry of hESC cultures for pluripotency markers OCT4 (green), SSEA4 (red) and alkaline phosphatase (black). Scale bar 100 µm.

**Supplemental Figure 18.**
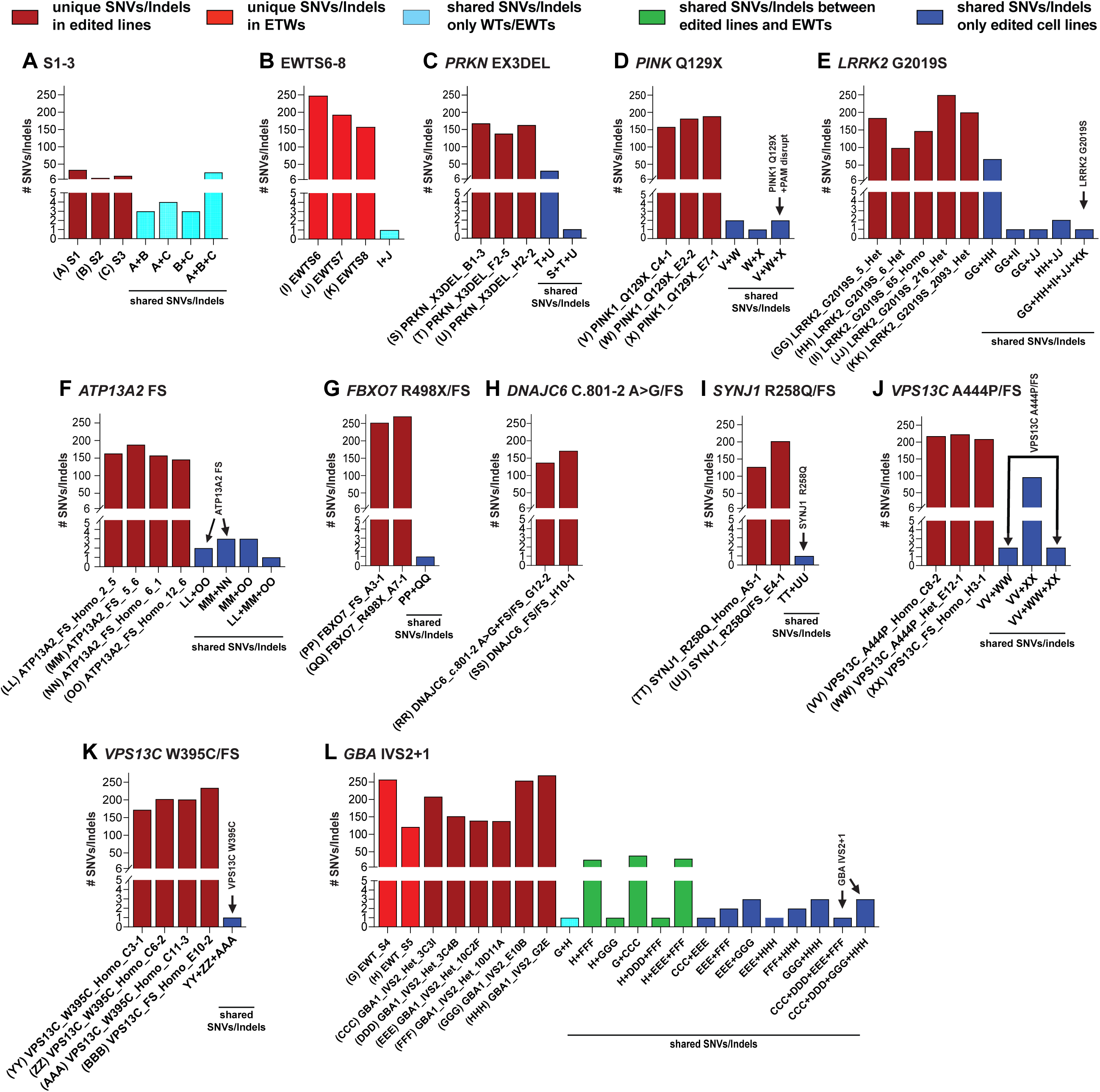
Graph showing number of unique and shared SNVs/indels in each editing experimental group: (A) WIBR3-S1/S2/S3, (B) WIBR3_EWTS6/7/8, (C) *PRKN* EX3DEL, (D) *PINK1* Q129X, (E) *LRRK2* G2019S, (F) *ATP13A2* FS, (G) *FBXO7* R498X/FS, (H) *DNAJC6* C.801-2 A>G/FS, (I) *SYNJ1* R258Q/FS, (J) *VPS13C* A444P/FS, (K) *VPS13C* W395C/FS and (L) *GBA* IVS2+1. Different color bars indicate unique or shared SNVs/Indels between different cell lines. Dark red indicates unique SNVs/Indels in edited lines, light red indicates unique SNVs/Indels in EWT lines, light blue indicates shared SNVs/Indels only found on WTs/EWTs, green indicates shared SNVs/Indels between edited lines and EWTs and dark blue indicates shared SNVs/Indels only found in edited cell lines.

## Code availability

All scripts used for the mapping and analysis of scRNASeq data can be accessed at https://github.com/ASAP-Team-Rio/iSCORE-PD_DAN_scRNASeq

## Data availability

All Genotyping and sequencing data is available at https://www.amp-pd.org/ via GP2 data sharing agreements This includes *(i)* Array genotyping data, *(ii)* Oxford Nanopore Technologies long-read sequencing *(iii)* Pacific Biosciences long-read sequencing, *(iv)* NGS and Sanger sequencing data for targeted genotyping and zygosity analysis, *(v)* single cell RNA-seq data from dopaminergic neuron differentiation and p53 pathway analysis, (vi) whole genome sequencing data

All other materials can be found in the Zenodo repository: 10.5281/zenodo.14907986. This includes *(i)* aCGH reports, *(ii)* ICC images (microglia, DA neurons, hESC), *(iii)* southern blot raw film and gel images, *(iv)* qRT-PCR result files/analysis for dopaminergic neuron differentiation, and *(v)* FACS-analysis results for microglia.

## Cell line availability

All cell lines in this isogenic collection will be banked and distributed by WiCell (https://www.wicell.org/) using material transfer agreements (MTAs).

## Acknowledgements.

We thank all the members of the Hockemeyer, Soldner, Rio, Blauwendraat and Bateup lab for helpful discussions and comments on the manuscript. This work was funded by Aligning Science Across Parkinson’s (ASAP-024409, ASAP-000486) through the Michael J. Fox Foundation for Parkinson’s Research (MJFF) to the Bateup, Rio, Hockemeyer, Blauwendraat and Soldner laboratories. This work was further supported in part by the Intramural Research Program of the National Institute on Aging (NIA, AG000542), and the Center for Alzheimer’s and Related Dementias (CARD), within the Intramural Research Program of the NIA and the National Institute of Neurological Disorders and Stroke. Some of the Flow Cytometry and Genomics shared resources at Albert Einstein College of Medicine were supported by the Cancer Center Support Grant (P30 CA013330). This work utilized the computational resources of the NIH HPC Biowulf cluster (http://hpc.nih.gov) and the Savio HPC cluster at UC Berkeley. This research has been conducted using the UK Biobank Resource under Application Number 33601. For the purpose of open access, the author has applied a CC-BY public copyright license to the Author Accepted Manuscript version arising from this submission. Long-read PacBio data generation was performed by Janet Aiyedun, Jackson Mingle, Jeff Burke, and Michelle Kim from PacBio.

## Author contribution

DH, HSB, DCR and FS conceived the project, supervised data analysis, and interpreted results. OB, HL, DH, and FS designed and supervised the genome engineering of cell lines and interpreted results. This cell line generation was assisted by KMS, JD, YV, GRP, JS, YD, VMS. Cell line expansion, sample collection and quality control experiments (NGS, ICC, aCGH array, LOH detection) were carried out by OB, HL, KMS, JD, RLB, YV, GRP, JS, YD, VMS, SP, ZB, JD, AS, JG. RLB performed microglia differentiation and analysis (ICC and FACS). KMS, JD performed differentiation of dopaminergic neurons and scRNA-seq analysis and data interpretation and graphing. AM performed the analysis of the p53 response in WIBR#3 cells. KMS carried out qPCR experiments. PAJ, DGH, KSL and CB performed long-read sequencing and high-density array analysis. EOB coordinated the deposition of the isogenic cell line collection with WiCell and edited the paper for open access requirements. MB performed the computational analysis of WGS data to assess genetic variability and off-target effects within the iSCORE-PD collection. OB and HL performed all other experiments. OB, HL, HSB, DH, CB and FS wrote the paper with input from all authors.

## Declaration of generative AI and AI-assisted technologies in the writing process

During the preparation of this work the authors used ChatGPT in order to improve the readability and language of the manuscript. After using this tool/service, the authors reviewed and edited the content as needed and take full responsibility for the content of the published article.

## Supplementary Note 1

### iSCORE-PD collection of isogenic hPSC lines carrying PD-associated mutations

For our initial iSCORE-PD collection, we prioritized engineering cell lines carrying mutations in high confidence PD genes^1^. The specific modifications for each gene were selected based on information in the MDSgene database^2^ (https://www.mdsgene.org) and the currently available literature as outlined for each gene below. Overall, 65 clonal cell lines carrying PD-associated mutations in 11 genes linked to PD along with isogenic control lines passed all the above-described quality control steps and have become part of the iSCORE-PD collection (Table 1, Supplemental Table 3). The current iteration of the iSCORE-PD includes the following cell lines:

#### Control cell lines

Together with the parental WIBR3 cell line, we provide a set of subclones with this collection (WIBR3-S1, S2, S3) (Figure 1, Supplemental figure 1). In addition, we included WIBR3 cell lines that were isolated as part of the standard genome editing pipeline but did not exhibit any genetic modifications at the targeted locus. We consider these as “edited wild-type” cells (EWT), which are the best experimental control to account for any non-specific changes caused by the gene editing process (EWT1-3: prime editing controls - Pipeline B, EWT4-5: CRISPR/Cas9 controls - Pipeline B and EWT6-8: prime editing controls - Pipeline A) (Table1, Supplemental Table 3, Supplemental Figure 6).

#### *SNCA* (PARK1)

The *SNCA* gene encodes the alpha-synuclein (ɑ-Syn) protein. The discovery that mutations and copy number increases of the *SNCA* gene linked to familial forms of PD, along with the identification of non-coding variants in the *SNCA* locus as a risk factor for sporadic PD, indicate a central pathogenic role for this protein^3^. Moreover, fibrillar ɑ-Syn is the major component of Lewy bodies, and changes in the dosage, aggregation, and clearance properties of ɑ-Syn are thought to be central pathogenic drivers of PD^1,3–5^. We established prime editing reagents to efficiently introduce the A30P or A53T mutation in the *SNCA* gene, both of which are linked to autosomal dominant forms of PD^6–8^. Using this approach (Supplemental Figure 7A,D), we generated cell lines that include three cell lines carrying the A53T mutation (heterozygous) (Supplemental Figure 7B,C) and four cell lines carrying the A30P mutation (3 heterozygous and 1 homozygous) (Supplemental Figure 7E,F). All cell lines passed the described quality control steps. In addition, we performed SNV-PCR followed by Sanger sequencing-based zygosity analysis to exclude LOH in the homozygous *SNCA* A30P line (Supplemental Figure 7G). Based on the described WGS analysis, we recommend including EWT_S1-3 as controls in disease modeling experiments for the cell lines carrying the A30P mutation.

#### *PRKN* (PARK2)

Mutations in the *PRKN* gene, which encodes the E3 ubiquitin-protein ligase parkin (Parkin), are the most frequent cause for autosomal recessive PD^9^. The first genetic alterations in *PRKN* linked to PD were distinct large-scale deletions^10^. Since then, a wide range of deletions and point mutations have been identified^11^. Molecular alterations in Parkin impact a wide range of molecular and cellular functions including responses to oxidative stress, mitochondrial membrane potential, mitochondrial motility, contact sites with the endoplasmic reticulum, and the regulation of inflammatory responses^12,13^. Homozygous and compound heterozygous deletions of Exon3 (Ex3del) are repeatedly found in families with PD, and Ex3del is now a confirmed pathogenic variant of *PRKN*^11,14^. We used a dual CRISPR/Cas9 approach (Figure 3C[v]) targeting each side of Exon3 to generate 3 cell lines that are homozygous for the Ex3del (Supplemental Figure 8A-C). Subsequent southern blot analysis to exclude LOH revealed that one cell line (WIBR3_PRKN_X3DEL_B1-3) carries a larger Ex3del (∼3.8 kb, Supplemental Figure 8D,E), which we fully characterized by NGS sequencing (Supplemental Figure 8F). We included this cell line in the iSCORE-PD collection since the flanking exons 2 and 4 are not affected by the deletion and thus, no significant consequences are expected of the extended deletion. No heterozygous clones have been included in this collection since these genotypes are frequent in control populations and are not associated with a higher risk of PD^15^.

#### *PINK1* (PARK6)

Mutations in *PINK1* were first associated with PD in 2004 following earlier studies linking the PARK6 genomic region with increased risk for PD^16^. Subsequently, a large number of point mutations, frameshift mutations and deletions have been identified, predominantly affecting the activity of the kinase domain^11^. This indicates that *PINK1* loss-of-function is the cause for early-onset, autosomal recessive PD. The serine/threonine-protein kinase PINK1 plays a crucial role in mitochondrial quality control by regulating mitochondrial homeostasis and clearance^17^. Based on the segregation of the Q129X and a related Q129fsX157 mutation in *PINK1* observed in two large families with PD^18^, we generated 3 cell lines that are homozygous for the Q129X mutation in *PINK1* (Supplemental Figure 9A-C). To exclude LOH, we performed SNV-PCR followed by Sanger sequencing based zygosity analysis (Supplemental Figure 9D). Heterozygous *PINK1* mutant cell lines were not included in the iSCORE-PD collection, as the heterozygous genotype is not considered a risk factor for PD.

#### *DJ1* (PARK7)

Since its initial association with PD^19^, multiple point mutations and genomic rearrangements in *PARK7* (*DJ1*), encoding the Parkinson disease protein 7, have been identified as a rare cause for autosomal recessive PD^11^. While the exact function of this enzyme remains largely unknown, Parkinson disease protein 7 is implicated in regulating transcription, cell growth and oxidative stress response pathways linked to cell survival and apoptosis^20–22^. Based on the identification of homozygous deletions of either exon 5^23^ or exon 1 to 5^24^ in families with PD, we used two step dual gRNA CRISPR/Cas9 approach to closely recapitulate the Exon 1 to 5 deletion (Supplemental Figure 10A-D). As outlined in detail above, the WGS analysis revealed a high number of shared SNVs/indels between the initially established clones (WIBR3_DJ1_X1-5DEL_2860/2872/2876).

To account for the potential impact of these shared variants on phenotypical analyses, we screened for an additional homozygous clone that was generated in a single targeting step (WIBR3_DJ1_X1-5DEL_6235). Additionally, we included three other homozygous DJ1/PARK7 clones (WIBR3_DJ1_EX1-5DEL_6348/6390/6407), which were generated by retargeting a second heterozygous cell line (WIBR3_DJ1_X1-5DEL_2046) that did not share SNVs/indels with the previously described homozygous clones (WIBR3_DJ1_X1-5DEL_2860/2872/2876). As there is currently no evidence that heterozygous genotypes confer an increased risk of developing PD^25^, we included several heterozygous DJ1/PARK7 lines as experimental controls (WIBR3_DJ1_X1-5DEL_Het_2036/2038/2046/2051/2067) that should account for the genetic variability of the homozygous targeted DJ1/PARK7 clones (Supplemental Figure 10). Southern blot analysis was performed to exclude LOH in the homozygous exon 1-5 deleted cell lines (Supplemental Figure 10E). Sanger sequencing of the wild-type allele in the heterozygous clones (WIBR3_DJ1_X1-5DEL_Het_2036/2038/2046/2051/2067) revealed additional SNVs/Indels at the target sites of the gRNAs used to generate these cell lines (Supplemental Figure 10F).

#### *LRRK2* (PARK8)

Mutations in the *LRRK2* gene, encoding the leucine rich repeat serine/threonine-protein kinase 2, were first linked to PD in 2004^26^. Over the years, more than 100 different variants of the gene have been described^27^, establishing *LRRK2* coding variants as the most frequently mutated gene linked to dominant and sporadic forms of PD. Among these mutations, G2019S is the most common substitution identified across populations^28^, found in approximately 4% of dominantly inherited familial PD cases, in both heterozygous and homozygous forms, and in around 1% of sporadic PD cases^29^. This variant increases the kinase activity of leucine rich repeat serine/threonine-protein kinase 2, affecting a wide range of cellular and molecular processes including vesicular trafficking and cytoskeleton dynamics, autophagy and lysosomal degradation, neurotransmission, mitochondrial function, and immune and microglial responses^30,31^. We recently established CRISPR/Cas9, TALEN and prime editing reagents to efficiently introduce the G2019S mutation in the *LRRK2* gene^6^. Using this approach (Supplemental Figure 11A), we generated additional cell lines so that the iSCORE-PD collection now includes 5 clones carrying G2019S (4 heterozygous and 1 homozygous) (Supplemental Figure 11B-C). We performed SNV-PCR followed by Sanger sequencing based zygosity analysis to exclude LOH in the homozygous *LRRK2* G2019S line (Supplemental Figure 11D).

#### ATP13A2 (PARK9)

Mutations in the *ATP13A2* gene, encoding the polyamine-transporting ATPase 13A2 (ATP13A2) protein, were identified as a cause for PD in the PARK9 locus in 2006, with this genomic region previously associated with Kufor Rakeb disease^32^. Various mutations in the gene result in a truncated ATP13A2 protein causing its mis-localization from the lysosome to the endoplasmic reticulum, where it accumulates before being eventually targeted for proteasomal degradation in the cytoplasm^32,33^. Loss of ATP13A2 is linked to mitochondrial and lysosomal dysfunction, as well as to the accumulation of ɑ-Syn due to the dysregulation of proteasomal and autophagy-mediated protein degradation^34–38^. Based on the identification of a homozygous frameshift mutation (NP_071372.1, T368RfsX29) in a family with recessive PD^38^, we used CRISPR/Cas9-based genome editing to engineer the frameshift mutation around the amino acid T368 in the *ATP13A2* gene (Supplemental Figure 12A). Given the recessive inheritance pattern of the gene, our collection contains 4 clones with biallelic frameshift mutations, predicted to result in a truncated loss of function protein similar to that observed in PD patients (Supplemental Figure 12B,C). We performed SNV-PCR followed by NGS sequencing-based zygosity analysis to exclude LOH in the homozygous edited cell lines (Supplemental Figure 12D).

#### *FBXO7* (PARK15)

Mutations in the *FBXO7* gene, encoding the F-box only protein 7 (FBOX7), were first linked to autosomal recessive PD in 2008^39^. The most notable homozygous *FBXO7* variant, found in multiple family pedigrees with PD, is the truncating R498X premature stop mutation located in the proline-rich region of the FBOX7 protein^40^. This mutation disrupts the interaction of FBOX7 with PINK1 and Parkin^41–43^, resulting in abnormal localization and reduced stability of the truncated FBXO7 protein. This causes disruption of mitophagy and leads to mitochondrial aggregation^41,44,45^. We used CRISPR/Cas9-based genome editing to engineer R498X and a frameshift mutation which leads to a premature stop and is predicted to result in truncated FBXO7, similar to the protein in patients carrying the R498X mutation (Supplemental Figure 13A-C). We performed Southern blot, SNV-PCR followed by NGS sequencing and analysis of WGS data to exclude LOH (Supplemental Figure 13D-F).

#### *DNAJC6* (PARK19)

The DNAJC6 gene encodes for the protein putative tyrosine-protein phosphatase AUXILIN, a member of the DNAJ/HSP40 family of proteins, which regulate molecular chaperone activity. DNAJC6 was initially linked to very early onset, autosomal recessive PD in 2012^46^ through the identification of a mutation that affects splicing and the expression of Auxilin. Consistent with its role as a co-chaperone that recruits HSC70 to clathrin-coated vesicles, hESC models show that *DNAJC6* alterations result in loss of Auxilin protein. This leads to the accumulation of clathrin, reduced vesicular transport, and the degeneration of midbrain dopaminergic neurons^47^. Furthermore, these alterations are associated with ɑ-Syn aggregation, mitochondrial and lysosomal dysfunction, and lipid defects^47–49^. We used a CRISPR/Cas9-based approach to recapitulate the effect of the c.801 -2A>G splice acceptor site mutation by either precise insertion of this mutation or a frameshift modification (Supplemental Figure 14). Similar frameshift modifications were previously shown to recapitulate the loss of *DNAJC6* expression in hESC-derived neuronal cells^47^, consistent with observations in *DNAJC6* variant carriers^46^.

#### *SYNJ1* (PARK20)

*SYNJ1* encodes Synaptojanin-1 and was first linked to early onset autosomal recessive PD in 2013^50,51^. Synaptojanin-1 is predominantly expressed in neurons and is concentrated in presynaptic terminals. Homozygous *SYNJ1* mutations are linked to alterations in lipid metabolism and vesicle trafficking^50,51^, as well as defects in autophagosome maturation^52^. We used a CRISPR/Cas9 approach to insert the R258Q substitution, which was identified in a homozygous state in several independent families with PD^50,51,53^ (Supplemental Figure 15A). The iSCORE-PD collection includes clones that are homozygous for the R258Q mutation in *SYNJ1* or compound heterozygous for the R258Q and a frameshift allele at the same location. Both genotypes are expected to recapitulate the modification in *SYNJ1* associated with disease (Supplemental Figure 15B,C). We performed Southern blot based zygosity analysis to exclude LOH in the homozygous edited cell line (Supplemental Figure 15D-F).

#### *VPS13C* (PARK23)

Mutations in *VPS13C* were initially identified as the cause of autosomal recessive early-onset PD in 2016^54^. The initial functional analysis revealed that disruption of intermembrane lipid transfer protein vacuolar protein sorting 13 homolog C (VPS13C) causes decreased mitochondrial membrane potential, mitochondrial fragmentation, increased respiration rates, exacerbated PINK1/Parkin-dependent mitophagy, and transcriptional upregulation of *PRKN* in response to mitochondrial damage^54^. As the VPS13C protein is critical for the transport of lipids between the ER and endosome/lysosome, as well as for lipid droplet formation^55,56^, loss of VPS13C causes the accumulation of lysosomes and altered lipid profiles^57^. We used CRISPR/Cas9-based editing to introduce the W395C and A444P variants into the *VPS13C* gene (Supplemental Figure 16A,E), both of which are found as homozygous or compound heterozygous mutations in PD patients^58,59^. While *VPS13C* is thought to cause PD through a loss of function mechanism, we included additional frameshift alleles in the iSCORE-PD collection to allow further investigation of the loss of function mechanism (Supplemental Figure 16B,C,F,G). We performed SNV-PCR followed by Sanger sequencing based zygosity analysis to exclude LOH in the homozygous *VPS13C* A444P lines (Supplemental Figure 16D).

#### GBA1

The *GBA1* gene codes for the enzyme Lysosomal acid glucosylceramidase, which is essential for maintaining glycosphingolipid homeostasis. While homozygous or compound heterozygous pathogenic variants in *GBA1*, associated with reduced glucosylceramidase activity, cause autosomal recessive Gaucher disease, heterozygous carriers of pathogenic *GBA1* variants have an elevated risk of developing PD^60,61^. *GBA1* mutations are currently considered the strongest risk factor for PD and Lewy body dementia with odd ratios between 1.4 to >10^62^. Given that *GBA1* mutations are present in approximately 3-20% of sporadic PD patients across different populations, *GBA1* represents the most prevalent genetic risk factor for PD^62^. Over 300 mutations in *GBA1* with variable risks for developing PD have been reported^61^. Among them, the splice site IVS2+1 mutation, causing missplicing and loss of *GBA1* expression, represents one of the most pathogenic alleles for PD^60,61^. To insert the IVS2+1 in the *GBA1* gene, we devised a CRISPR/Cas9-based targeting strategy that allows specific targeting of the *GBA1* gene and not the nearby highly homologous *GBAP1* pseudogene (Supplemental Figure 17A). To identify correctly targeted clones, we used a genotyping strategy that can conclusively distinguish between the *GBA1* and *GBAP1* pseudogene based on small sequence variation (Supplemental Figure 17B-D). Using this approach, we generated 2 heterozygous and 2 homozygous edited cell lines carrying the IVS2+1 mutation in *GBA1*. In addition, to allow comparison between the IVS2+1 and a loss of function allele, we also included cell lines with a frameshift mutation at the same genomic location in the iSCORE-PD collection.

